# Yin-Yang homeodynamics of Nrf1 and Nrf2: coupling antioxidant defense to redox recovery bistability under persistent stress

**DOI:** 10.64898/2026.11.16.766932

**Authors:** Ruiguang Wang, Xi Chen, Lu Qiu, Zongyi Zhou, Ruiqi Xiong, Zhengwen Zhang, Ping Ao, Yiguo Zhang

**Author notes:** equal contribution to this work (Ruiguang Wang, Xi Chen).

## Abstract

This work systematically articulates the Yin-Yang homeodynamics coordination between Nrf1 and Nrf2, resolving the longstanding core paradox in redox biology of why robust Nrf2-mediated antioxidant activation frequently fails to fully eliminate unresolved oxidative stress and re-establish sustained redox homeostasis. Integrating multi-omic expression profiles from human HepG2 cells and primary murine hepatocytes, we constructed an 11-variable endogenous regulatory network model, demonstrating that Nrf2 exclusively mediates transient acute stress attenuation responses, while Nrf1 functions as the non-redundant upstream organiser that supplies the indispensable preconditions for definitive stress resolution via three irreplaceable homeostatic axes: organelle integrity maintenance, proteasome functional recovery, and endoplasmic reticulum (ER) cholesterol steady-state handling. Their coupled regulatory interactions generate a bistable configuration demarcated by an index-1 saddle point, partitioning the physiological state space into native low-stress homeostasis and the previously uncharacterised persistent-stress attractor. Once self-amplified unresolved endogenous stress generation reaches dynamic flux balance with Nrf2-dependent clearance at an elevated burden level, the system becomes irreversibly locked in this persistent high-burden equilibrium, and cannot spontaneously return to native homeostatic status even after the initiating upstream disturbance has been fully corrected. Collectively, this study fundamentally delineates the clear functional division between transient antioxidant activation and durable redox recovery during Nrf1/Nrf2 homeodynamics, hence providing a rigorous systems-level theoretical framework to guide future translational interventions targeting redox homeostasis restoration.

**Highlights:**

⋄ Nrf1 restores redox balance while Nrf2 only buffers acute stress
⋄ Robust antioxidant defense can lock model cells in persistent stress
⋄ Endogenous bistable network reveals irreversible stress attractor
⋄ Membrane cholesterol handling gates reversibility of redox recovery
⋄ Transcriptomics and modeling show exposure history decides fate

**Graphical abstract:** Graphical abstract.
Nrf1 and Nrf2 coordinate antioxidant defense with organelle homeostasis, proteasome function and cholesterol handling. These processes are linked by the endogenous network to stress resolution and persistent compensation after stimulus withdrawal. Proteasome-supported INSIG1 degradation is shown as a branch separate from KEAP1-CUL3-dependent Nrf2 turnover. The arrow to INSIG1 degradation denotes promotion of the degradation process, and therefore a reduction in INSIG1 abundance [1]. Solid arrows indicate functional promotion, terminal bars indicate restraint or clearance, and dashed connections summarize effective network feedback and alternative recovery outcomes.

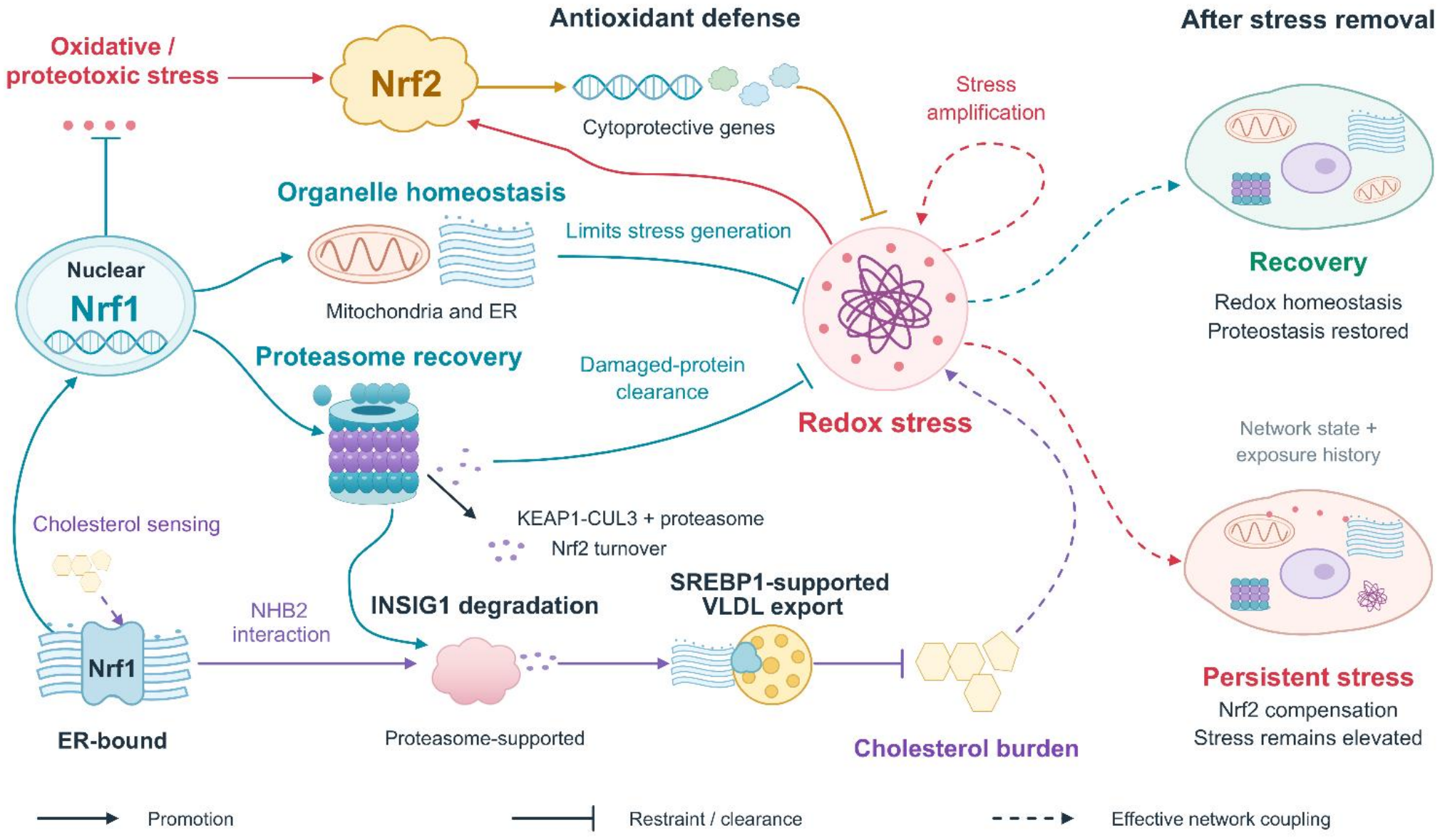

## 1. Introduction

Oxidative stress represents a patho-physiological disruption of the coordinated processes that maintain redox homeostasis. Antioxidant induction constitutes an essential component of the cellular response to oxidative stress, yet sustained antioxidant cytoprotection additionally requires tight regulation of endogenous stress production and efficient clearance of irreversibly damaged cellular components. These layered functions are predominantly executed by the two core antioxidant CNC-family transcription factors: Nrf1 (encoded by *NFE2L1*) and Nrf2 (encoded by NFE2L2) [2]. In our previous characterization of *Nrf1α*-deficient cells, we observed persistent accumulation of reactive oxygen species (ROS) and oxidative damage, alongside markedly-elevated Nrf2 activation, accompanied by impaired mitochondrial respiration [3], endoplasmic reticulum stress [4], DNA damage and chromosomal instability [5]. This distinctive phenotype raises a central mechanistic question: why can a robust, biochemically active antioxidant response coexist with unresolved persistent cellular stress?

The biological distinction between Nrf1 and Nrf2 extends far beyond reciprocal alterations in the expression profiles of downstream genes/proteins. Based on systematic synthesis of existing evidence from our laboratory and other independent groups [2,3,6–8], Nrf1 occupies a non-redundant central position within the organizational framework that sustains long-term redox homeostasis, whereas Nrf2 acts as the canonical master regulator governing context-responsive inducible cytoprotective pathways [2]. Cumulative genetic and stress-response studies have demonstrated that their downstream transcriptional outputs operate cooperatively in some defined physiological contexts but diverge functionally in others [4,9]. Such dialectically-unified functional inter-relationship is herein defined as a Yin-Yang coordination paradigm: Nrf1-mediated homeostatic maintenance and Nrf2-dependent inducible defense represent two intrinsically-coupled, complementary components of a unified, integrated cytoprotective network. Both transcription factors engage in cellular stress adaptation by driving cells toward distinct phenotypic response states, and thereby the homeodynamics of their relative contributions determines the extents to which antioxidant induction is accompanied by the full or partial restoration of normal cellular function.

Three distinct Nrf1-dominant functional axes anchor this Yin-Yang regulatory relationship directly to core oxidative-stress biology. Nrf1 mediates proteasome functional recovery following pharmacological or genetic proteasome inhibition [6], integrates redox signaling regulation with mitochondrial functional homeostasis [3], and acts as a direct cholesterol sensor localized to the endoplasmic reticulum (ER) membranes and their adjacent lipid microdomains [7]. The recently characterized physical interaction between Nrf1 and INSIG1 further defines a previously unrecognized membrane-regulatory pathway, through which Nrf1 sustains certain transcriptional activity of SREBP1 and maintains VLDL secretory export under conditions of cellular cholesterol excess [1]. Furthermore, the complementary Nrf1 and Nrf2 co-regulated gene transcription also confer robust cytoprotection against pathological hepatic cholesterol overload *in vivo* [8]. Collectively, these interconnected findings functionally connect the canonical Nrf2-driven antioxidant defense programming to the broader regulatory networks governing proteostasis, organelle structural and functional integrity, and cellular cholesterol trafficking and handling, providing a rigorous biological foundation for investigating unresolved persistent stress as a systems-level emergent property of this endogenous integrated cytoprotective network.

The proteasome-recovery functional arm of this regulatory network is also strictly governed by post-translationally regulated Nrf1 maturation [10]. Topovectorially regulated DDI2-mediated proteolytic cleavage and NGLY1-dependent deglycosylation processing together enable full Nrf1 transcriptional activation after proteasome impairment [11,12]. More recently published work has further identified sugar-linked ubiquitination and sequence-editing-dependent coactivator recruitment as two additional, previously uncharacterized layers of post-translational control to fine-tune this compensatory transcriptional response [13,14]. Overall, these multi-layered regulatory mechanisms position Nrf1 activity as an integral downstream component of a larger protein-processing and quality-control clearance circuit, rather than as a simple linear readout of its steady-state transcript abundance.

Quantitative studies of the canonical Keap1-Nrf2 signaling axis have systematically delineated how cytosolic sequestration, ubiquitin-dependent protein degradation and negative feedback amplification jointly shape the dynamic profile of antioxidant induction and its homeostatic regulation [15,16]. However, a critical, largely unaddressed complementary problem in this field concerns the systems-level recovery of the wider cellular functional network: after the initial Nrf2-driven antioxidant program has been fully activated, which specific combinations of homeostatic maintenance, targeted protein turnover, and endogenous stress generation dynamics ultimately permit the complete restoration of physiological redox homeostasis? Rigorously resolving this mechanistic question requires longitudinal tracking of these interconnected, mutually regulatory processes beyond simple measurement of the amplitude of Nrf2 nuclear activation alone. Recently, endogenous-network studies conducted in hepatocellularcarcinoma and gastric cancer cell systems have established a robust analytical framework for connecting local molecular feedback loops to the emergence of persistent, long-lived cellular phenotypic states [17,18]. In the present work, this established systems-level framework is specifically applied to dissect the fundamental functional distinction between transient antioxidant pathway activation and durable, complete stress resolution.

Here, we systematically integrate empirical laboratory-driven gene and protein expression profiles with *in vivo* liver histology evidence, along with a custom-built endogenous-network computational model that unifies three core regulatory modules: Nrf1-mediated homeostatic maintenance, canonical Nrf2 signaling cascades, and membrane cholesterol homeostatic regulation. We combine multiple complementary analytical approaches, including steady-state phenotypic characterization, joint Nrf1-Nrf2 genetic perturbations, dynamic pulse-washout kinetic simulations and local stochastic state-space analysis. Our integrated calculations rigorously define distinct sets of biological conditions under which Nrf2-driven antioxidant compensation effectively reduces acute oxidative stress but fails to fully resolve it, leaving a non-reversible persistent-stress cellular state intact, and alternative sets of conditions in which Nrf1-dependent homeostatic maintenance pathways restore the system’s dynamic accessibility to physiological low-stress states. This unified conceptual framework positions the Yin-Yang functional coordination between Nrf1 and Nrf2 at the critical regulatory interface between transient antioxidant defense activation and the durable, complete resolution of unresolved oxidative stress.

## 2. Materials and methods

### 2.1. Biological network and state variables

The endogenous regulatory network described in this work is constructed with eleven normalized dimensionless functional state variables: nuclear Nrf1 transcriptional activity (defined as *N*), organelle and basal redox homeostatic maintenance capacity (*M*), total cellular proteasome functional activity (*P*), intracellular redox stress burden (*R*), Nrf2-driven inducible antioxidant response activity (*Y*), functional Keap1-mediated cytosol restraint capacity (*K)*, total p62-associated protein abundance (*S*), endoplasmic reticulum (ER) membrane free-sterol signalling intensity (*C*), INSIG1 protein abundance (*I*), SREBP1-supported hepatic lipid-export transcriptional program activity (*V*) and total hepatic cholesterol burden (*H*). Of note, the membrane-tethered Nrf1 protein availability is explicitly formulated as algebraic function of the ER-free sterol signal *C* (full derivation provided in Supplementary Methods S10.4, Eqs. (S43)-(S46)), with a dedicated, independently adjustable input parameter. This formalization explicitly distinguishes the nuclear-localized transcriptionally active Nrf1 pool and membrane-associated sequestered Nrf1 isoform as two functionally distinct, non-redundant regulatory contributions within this network.

Nrf1-dependent homeostatic regulation of proteasome functional recovery and organelle integrity is formalized according to functional relationships established in prior experimental studies [3,6]. Keap1-mediated ubiquitination and proteasomal targeting, combined with total cellular proteasome activity, jointly determine the basal turnover rate of Nrf2 protein; p62 acts as a well-characterized competitive decoy substrate that sequesters Keap1 and thereby reduces its effective inhibitory restraint on Nrf2 nuclear translocation and transcriptional activity [15,19]. Nrf1-dependent p62 transcription and subsequent autophagic clearance of p62 aggregates are implemented as two decoupled terms, to avoid non-biological crosstalk between these distinct processes [20]. Studies of ULK1-dependent p62 phosphorylation and MOAP-1-mediated p62-body dissociation inform the biological interpretation of p62-mediated Keap1 sequestration and release [21,22]; these processes are represented by effective coupling terms rather than individually fitted kinetic reactions. The full network retains physiologically relevant nonzero basal production rates for maintenance and turnover processes and Nrf2-supported cytoprotective maintenance pathways that include the previously documented Nrf2-dependent auxiliary support for core proteasome subunits’ transcriptional expression [23]. The direct Nrf1-Keap1 regulatory coupling explicitly captures the recently reported protein-level functional dependency that motivated the construction of this integrated network model [9], and the phenotypic consequences of this specific interaction are systematically interrogated via targeted *in silico* topological deletion assays (Supplementary Methods S4). The membrane-associated sterol regulatory branch couples Nrf1-dependent INSIG1 protein degradation to downstream SREBP1-mediated transcriptional activation of lipid export programs and subsequent hepatic cholesterol trafficking and homeostatic handling [1].

All 11 normalized dimensionless state variables in this endogenous network follow the standard bounded production-loss ODE dynamics with a built-in Hill function formulation to describe the saturable, cooperative kinetics of all transcriptional and post-translational regulatory interactions across the full system. The governing balance and common Hill gate are

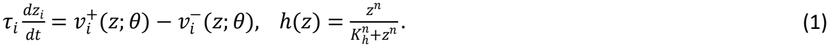

Equation (1): the balance and response gate are specified in Supplementary Methods S2, Eqs. (S2) and (S4); normalization and derivation are given in S10.2, Eqs. (S29)-(S33).

In this formula, all regulatory strength parameters and relative kinetic time-scale constants (denoted *τ*_*i*_) are defined as dimensionless quantities to enable direct cross-pathway functional comparison, eliminating unit-dependent scaling artifacts that arise when integrating processes with disparate intrinsic kinetics spanning transcriptional regulation, protein degradation and metabolic flux. The oxidized/aggregated state variable R represents the combined intracellular burden of redox stress and proteotoxic stress, cumulatively originating from organelle functional failure, irreversibly misfolded or damaged protein aggregates, and dysregulated sterol metabolism, all normalized to a 0–1 unitless scale. The production term 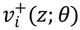 and loss term 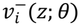 explicitly describe the continuous generation and active resolution of this combined stress burden *via* the interconnected cytoprotective and homeostatic functions encoded in the network. Full explicit differential equations for all 11 state variables, all reference parameter values and their numerical provenance, and detailed biological evidence assignments for every individual regulatory interaction are provided in Supplementary Methods S0-S3. The full dynamical framework serves as the unified computational foundation for all subsequent steady-state attractor identification, combinatorial Nrf1-Nrf2 perturbation screening, pulse-chase kinetic simulation, and local stochastic state stability analyses presented in this work.

### 2.2. Stress generation and resolution

The normalized stress-burden *R* is governed by a dynamical equation explicitly decomposed into a source flux and a removal flux, i.e., source and removal terms:

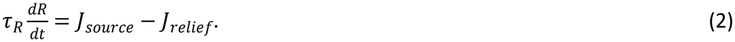

Equation (2): the source and relief terms are expanded in Supplementary Methods S2, Eqs. (S8)-(S9), and derived in S10.3, Eqs. (S34)-(S35).

Here, the dimensionless characteristic time constant τ_*R*_ unifies disparate kinetic rates across transcriptional regulation, protein turnover and metabolic processes, eliminating unit-dependent scaling artifacts to enable cross-pathway quantitative comparison.

For the total stress generation flux *J*_*source*_, five contributions are integrated: constitutive basal metabolism-derived background stress, a stress term proportional to organelle homeostatic maintenance deficiency, a non-linear positive feedback amplification of unresolved residual stress, a sterol dysregulation-coupled stress input, and a tunable optional external stress pulse for dynamic perturbation assays. For the total stress relief flux *J*_*relief*_, three effective resolution contributions are defined: constitutive basal turnover of oxidized and damaged moieties, a proteasome-associated contribution to removal of damaged-protein burden, along with Nrf2-driven transcriptional antioxidant programs that mediate broad-spectrum stress attenuation. All individual flux components presented in subsequent decomposition panels are normalized by *τ*_*R*_ to generate uniformly scaled, directly comparable time-series profiles. Finally, the two key explicit effective feedbacks—unresolved-stress amplification and sterol-to-stress coupling—are systematically dissected *via* targeted *in silico* topological deletion, to quantitatively isolate their respective contributions to the formation of persistent non-resolving stress states.

### 2.3 Nrf1 and Nrf2 regulatory equations

The principal Nrf1-Nrf2 balances are written explicitly below. Together with the stress balance in Eq. (2), Eqs. (3)-(5) describe the seven-variable redox and proteostasis core; Eq. (6) specifies membrane-associated Nrf1 control of INSIG1 turnover. All symbols, reference parameter values and numerical provenance are listed at the start of Supplementary Methods S0.

Here *γ*_n_ is the nuclear Nrf1 input, *o* = *q*_2_*h*(*Y*) is effective Nrf2 output, and *h*_*c*_(*C*) is the ER sterol response function. The optional feedback gains *a*_*N*_ and *a*_2*N*_ are zero in the reference system. The product of *K* and *P* represents joint Keap1-proteasom+e-dependent Nrf2 turnover. An overdot denotes a derivative with respect to model time.

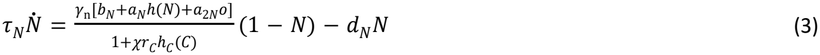

Equation (3): the nuclear Nrf1 balance is specified in Supplementary Methods S2, Eq. (S5), with its biological construction explained in S11.1.

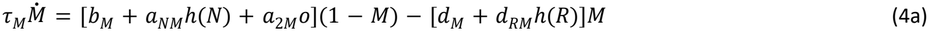

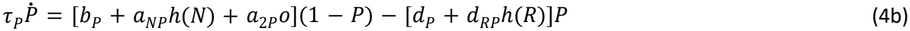

Equations (4a) and (4b) correspond to Supplementary Methods S2, Eqs. (S6) and (S7), respectively; their construction is explained in S11.1.

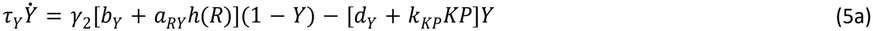

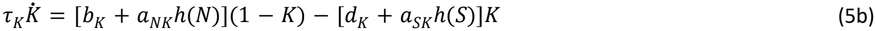

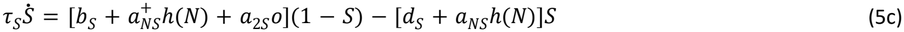

Equations (5a)-(5c) correspond to Supplementary Methods S2, Eqs. (S10)-(S12), with their construction in S11.1. The separate Nrf1 contributions to p62 production and clearance are derived in S10.3, Eq. (S42).

These balances couple nuclear Nrf1 activity to organelle maintenance and proteasome function, and link stress-induced Nrf2 output to Keap1-proteasome turnover and p62 feedback. Membrane-associated Nrf1 availability and INSIG1 abundance are specified separately:

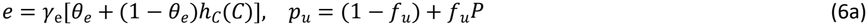

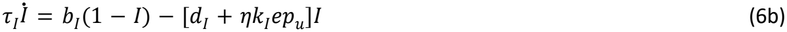

Equation (6a) is specified in Supplementary Methods S2, Eq. (S3), and derived in S10.4, Eqs. (S43)-(S44). Equation (6b) follows from S2, Eq. (S14), and S10.4, Eqs. (S44)-(S45); its regulatory signs are derived in S11.1, Eq. (S67).

The ER signal, lipid-export program and sterol-burden balances complete the eleven-variable system (Supplementary Eqs. (S13)-(S16)). Stepwise construction of Eqs. (3)-(6) is given in Supplementary Methods S11.1, using the bounded-balance derivation in S10.2, the compensation analysis in S10.3 and the membrane closure in S10.4.

### 2.4. Analyses of steady states and Nrf1-Nrf2 perturbation

Steady states of the 11-dimensional system were identified *via* a two-stage multi-start root-finding workflow: initial dimension reduction by exact equilibrium elimination to 3 residual coordinates, followed by an independent full 11-dimensional search. The reference search was initialized from 1024 scrambled Sobol sequence sampling points supplemented by strategically structured initial conditions, with all accepted root solutions satisfying a strict full-state residual threshold below 10 ^−8^ Local asymptotic stability of each identified steady state was systematically assessed by calculating the full eigenvalue spectrum of the 11-dimensional system Jacobian. An index-1 saddle point, defined by exactly one eigenvalue with a positive real part, was further characterized by numerically integrating its two unstable invariant branches to resolve their respective terminal stable attractors.

For the Nrf1-Nrf2 intervention map, we sampled a 21×21 parameter grid: 21 matched graded levels of nuclear and membrane Nrf1 inputs spanning the 0–1 normalized range, paired with 21 Nrf2 production multipliers from 0 to 4. All 441 independent simulations were initialized from the validated reference low-stress homeostatic state and integrated for 250 dimensionless model time units. Numerical consistency was confirmed by cross-validating fourth-order Runge-Kutta integration at step sizes [0.05, 0.025], with selected representative endpoints that were further verified *via* the DOP853 adaptive step-size solver. This graded perturbation design by input parameters enables direct quantitative comparison of network activity under reduced, restored and enhanced functional states.

All subsequent analyses including the cholesterol-input dose-response map, 128-scenario parameter ensemble, 9-level Hill coefficient sensitivity screening, targeted topological deletions, and compensatory bypass scan followed the protocols detailed in Supplementary Methods S4-S5. The compensatory bypass scan specifically quantifies the phenotypic outcome of artificially elevating Nrf2-supported organelle/proteasome maintenance and alternative sterol clearance after full abrogation of both nuclear and membrane Nrf1 inputs. The existence of stable steady-states and the capacity for stress resolution recovery from a predefined initial state were evaluated as two independent functional readouts.

### 2.5. Pulse-washout and recovery protocols

All dynamic simulations are implemented on the established 11-dimensional endogenous regulatory network framework, to systematically dissect the threshold boundary between transient antioxidant activation and complete, durable stress resolution. External stress and sterol perturbation pulses were applied for 6 discrete dimensionless durations (0.5, 1, 2, 4, 8 or 16 model time units) and then fully removed. For each duration, 25 amplitudes were sampled separately for each of the two input types, generating a total of 300 independent pulse-washout simulation protocols. All post-pulse system trajectories were integrated and tracked for an additional 250 dimensionless model time units.

Post-washout recovery status was quantitatively classified by calculating the full 11-coordinate Euclidean distance between the trajectory endpoint state vector and each of the two reference attractors, assigning the closer attractor. The classification consistency of all 300 protocols was cross-validated after halving the numerical integration step size to check numerical consistency. Precomputed bistability threshold brackets derived from 16 rounds of bisections, were retained as independent, standardized numerical summaries of state transition boundaries.

All targeted recovery intervention assays were initialized from the well-characterized persistent-stress state, which is spontaneously established after disruption of the Nrf1-INSIG1 membrane interaction. Five parallel intervention conditions were configured: no-intervention negative control, direct restoration of Nrf1-INSIG1 membrane interaction, downregulation of INSIG1 synthesis, fourfold scaling of the entire Nrf2 production term, and combinatorial intervention coupling membrane interaction restoration with Nrf2 production enhancement. The post-intervention dynamic trajectory of each condition was monitored for 200 dimensionless model time units.

Consistent with simulation outputs, pulse duration exerts a strong modulatory effect on the transition threshold into persistent stress state: the critical amplitude threshold for stress input decreases from 0.330 (0.5 time unit pulse) to 0.047 (16 time unit pulse), while the threshold for cholesterol input decreases from 1.773 to 0.108. Trajectories under sub-threshold and supra-threshold pulses at ∼2 time units both activate robust Nrf2 antioxidant response, but only sub-threshold stimuli permit complete return to the low-stress state after washout; supra-threshold stimuli sustain elevated Nrf2 signaling with high residual stress, even after full removal of external inputs, distinguished exclusively by whether the coordinated recovery of organelle maintenance, proteasome function and redox homeostasis is accomplished, rather than by the presence of Nrf2 activation alone. Intervention assays further confirm that even after full correction of upstream cholesterol dysregulation, the feedback-sustained persistent stress state remains irreversibly stabilized, demonstrating that the self-maintaining positive feedback circuit can operate independently of the original initiating perturbation.

### 2.6. Analysis of local stochastic dynamics

Characterisation of local stochastic dynamics was performed systematically around each stable steady state of the 11-dimensional network, using two independent, physically distinct diffusion conventions: isotropic diffusion and effective-flux derived diffusion. This entire analysis is rigorously grounded in the non-equilibrium stochastic decomposition framework established by Ao and co-workers [24,25], adopting its explicit linearised construction scheme strictly valid in the local neighbourhood of fixed points [26]. The local covariance matrix at each stable state was first solved directly from the Lyapunov equation, which was then used to construct a local A-type Ao decomposition [27,28], ensuring mathematically self-consistent matching of drift terms, diffusion tensor and underlying quasi-potential structure.

Subsequently, bounded finite-time action minimisation was implemented to resolve the 11-dimensional candidate transition trajectories connecting each stable state attractor to the unique index-1 saddle point. Optimisation runs were initialised from multiple independent seed paths, with systematic multi-resolution discretisation cross-checks to assess the numerical consistency of the final outputs [29,30]. This tiered analytical design explicitly segregates two physically distinct quantities: the intrinsic fluctuation spectrum confined to the immediate neighbourhood of each steady state, and the characteristic escape cost for inter-attractor transition in the weak-noise limit. The former quantifies the native diffusive dispersion of the system around its homeostatic baseline, while the latter quantifies a dimensionless action cost separating the low-stress and persistent-stress stable states. Full derivations of the local A-type stochastic construction, its corresponding local Itô process formulation, and the numerical implementation protocol for path-action calculation are provided in Supplementary Methods S6-S7 and S10.6-S10.8.

The stochastic formulation uses the same time-scaled drift as Eq. (1):

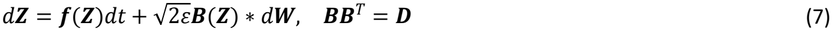

Equation (7) is specified in Supplementary Methods S6, Eq. (S19). Its Itô conversion and noise normalization are derived in S10.6, Eqs. (S57)-(S61), and the local covariance construction in S10.7, Eqs. (S62)-(S63).

The asterisk denotes the A-type convention, *ε*is the noise intensity and ***D*** specifies diffusion. Supplementary Methods S10.6 derives the corresponding Itô drift; S10.7 derives the local covariance and potential, and S10.8 connects the transition action to a compatible potential difference. The numerical path-action gradient is derived in S11.3.

### 2.7. Human HepG2 cell expression profiles and animal liver tissue analyses

The HepG2 transcriptomic matrix encompasses 5 distinct genetic backgrounds, each with 3 independent, biologically replicated samples derived from laboratory-controlled experiments, yielding a total of 15 profiled samples: wild-type control, *Nrf1α deficiency, Nrf1α deficiency* with concurrent Nrf2 siRNA knockdown (si*Nrf2*), *Nrf2*^ΔTA^ mutant, and constitutively active *Nrf2*^ΔN^ mutant. Prior to normalisation, duplicate gene symbol entries were averaged on the raw FPKM expression scale to eliminate redundant measurements, followed by a standard log_2_(FPKM + 1) logarithmic transformation, after which every individual gene was standardised to unit variance across the full 15-sample dataset. Four pre-specified, mutually exclusive functionally curated gene sets were interrogated for coordinated transcriptional module scoring: the full proteasome subunit cohort, Nrf1 proteolytic processing-associated genes, direct Nrf2-responsive antioxidant output genes and the core integrated stress response (ISR) gene set. Full gene set compositions and verified retained gene coverage across the two datasets are fully documented in Supplementary Methods S8. All derived module scores explicitly quantify the aggregated, co-ordinated activity of each predefined transcriptional programme, rather than isolated individual gene expression changes.

The primary hepatic transcriptomic dataset comprises 7 independent sample-level averaged expression profiles derived from murine primary hepatocyte populations. The cohort is hierarchically stratified to capture age-dependent and genotype-dependent effects: one 8-week wild-type normal sample (A3), two 8-week *Nrf1α*-deficient samples (A5T, A6T), two 9-month wild-type normal samples (A8, A9), and two 9-month *Nrf1α*-deficient hepatocellular carcinoma (HCC)-associated tissue samples (A13T, A14T). The HepG2 and liver tissue datasets were standardised completely independently, with each individual sample designated as the fundamental unit of all downstream statistical analysis. All animal liver tissue sampling procedures were conducted in full compliance with institutional guidelines for humane animal experimentation, with all protocols pre-approved by the local Institutional Animal Care and Use Committee to minimise animal distress.

Representative H&E-stained histological fields were extracted from laboratory-provided presentation materials, retaining all original image content, pixel resolution and embedded scale bar calibration intact. Histological assessment was conducted exclusively *via* morphological characterisation, albeit with no animal-level paired statistical linkage performed between histological observations and individual animal transcriptomic profiles. Transcriptional module scoring and histological observation constitute two orthogonal, non-paired analytical modalities used to independently quantify cellular transcriptional programme activation status and macroscopic tissue morphological features, respectively. Direct quantitative biochemical measurement of absolute redox species concentrations is outside the scope of the present transcriptomic characterisation, and requires dedicated, rigorously validated redox-specific biochemical assays [31], which will be further detailed in the other manuscript. The animal study underlying the mouse liver histological sections was reviewed and approved by the Ethics Committee of Chongqing University Cancer Hospital.

### 2.8. Statistical analysis and reproducibility

For the HepG2 transcriptomic dataset, all 20 possible allocations of 3-sample pairs in two independent genetic backgrounds were subjected to two-sided exact permutation testing. Benjamini–Hochberg multiple-testing correction was applied across the pre-specified panel of 16 transcriptional module comparisons [32]. Given the discrete sampling configuration, the smallest attainable two-sided *P*-value for this experimental design is strictly 0.10. All gene expression analyses were therefore formally designated exploratory, with results reported exclusively via validated, statistically robust descriptors: explicit effect direction, normalised effect magnitude, and independent individual biological observations, avoiding overinterpretation of spurious single-gene significance that would be unrepresentative of coherent transcriptional programme activity. The primary murine liver hepatocyte transcriptomic dataset was characterised exclusively at the individual sample level, with no paired statistical linkage performed at the individual animal level, in accordance with the predefined orthogonal analytical modality for the tissue cohort.

All deterministic and stochastic dynamical calculations presented in this work were executed on the validated dimensionless 11-state-variable ODE framework, built upon the bounded production-loss equation system

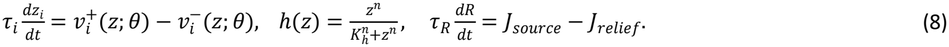

with local A-type decomposition constructed from the Lyapunov covariance equation for fixed-point neighbourhood fluctuations. Transition costs were calculated separately by finite-time path-action minimisation. All numerical integrations, steady-state root-finding with multi-start Sobol sampling, and topological deletion assays were conducted using Python 3.11, with supporting numerical libraries NumPy 2.4.4 and SciPy 1.17.1. Quantitative figures and all associated statistical visualisations were generated entirely within R 4.3.0, using the standardised package stack ggplot2 3.5.1, patchwork 1.2.0, ggrepel 0.9.5 and svglite 2.1.3. The revised graphical abstract is a raster illustration; quantitative figures were exported as vector artwork and high-resolution raster images using svglite and ragg. ChatGPT and Codex (OpenAI) were used as auxiliary tools for mathematical derivations, numerical model implementation, and code development. The resulting mathematical expressions and computational procedures were reviewed and independently verified by the authors. Full implementations of all governing equations, curated parameter value source tables, curated laboratory image provenance records, tiered stepwise numerical consistency checks across multiple integration solvers and step sizes, and fully reproducible plotting and analysis source code are deposited alongside the manuscript to guarantee full traceability and independent replication of all reported results.

Of note, equation (8) restates Eqs. (1) and (2) for the reproducibility summary. The complete terms are given in Supplementary Methods S2, Eqs. (S2), (S4) and (S8)-(S9), with derivations in S10.2-S10.3, Eqs. (S29)-(S35).

## 3. Results

### 3.1 Nrf1α-deficiency dissociates antioxidant induction from proteostasis programs

In *Nrf1α*-deficient HepG2 cells, we observed a fully coordinated, genome-wide transcriptional response in which robust Nrf2-driven antioxidant induction was reciprocally and functionally decoupled from the canonical proteostasis maintenance transcriptional programme, an effect entirely consistent with the prior experimental observation that sustained Nrf2 activation alone fails to eliminate accumulated reactive oxygen species and downstream irreversible cellular damage. Relative to isogenic wild-type control, the mean normalised proteasome module score decreased by 1.487 standardized units and the Nrf1-processing-associated gene module score decreased by 0.879 standardised units (*su*), while the integrated stress response (ISR) module score increased by 1.411 *su* and the predefined 5-gene Nrf2-responsive antioxidant module score increased by 0.298 *su* (Fig. 1A-C). These mutually opposing directional changes across functionally distinct transcriptional modules in the identical genetic background establish that cytoprotective antioxidant programmes and proteostasis maintenance programmes are not coregulated as a unified unit, but can be fully dissociated under single-gene perturbation of *Nrf1α*.

**Figure 1.**
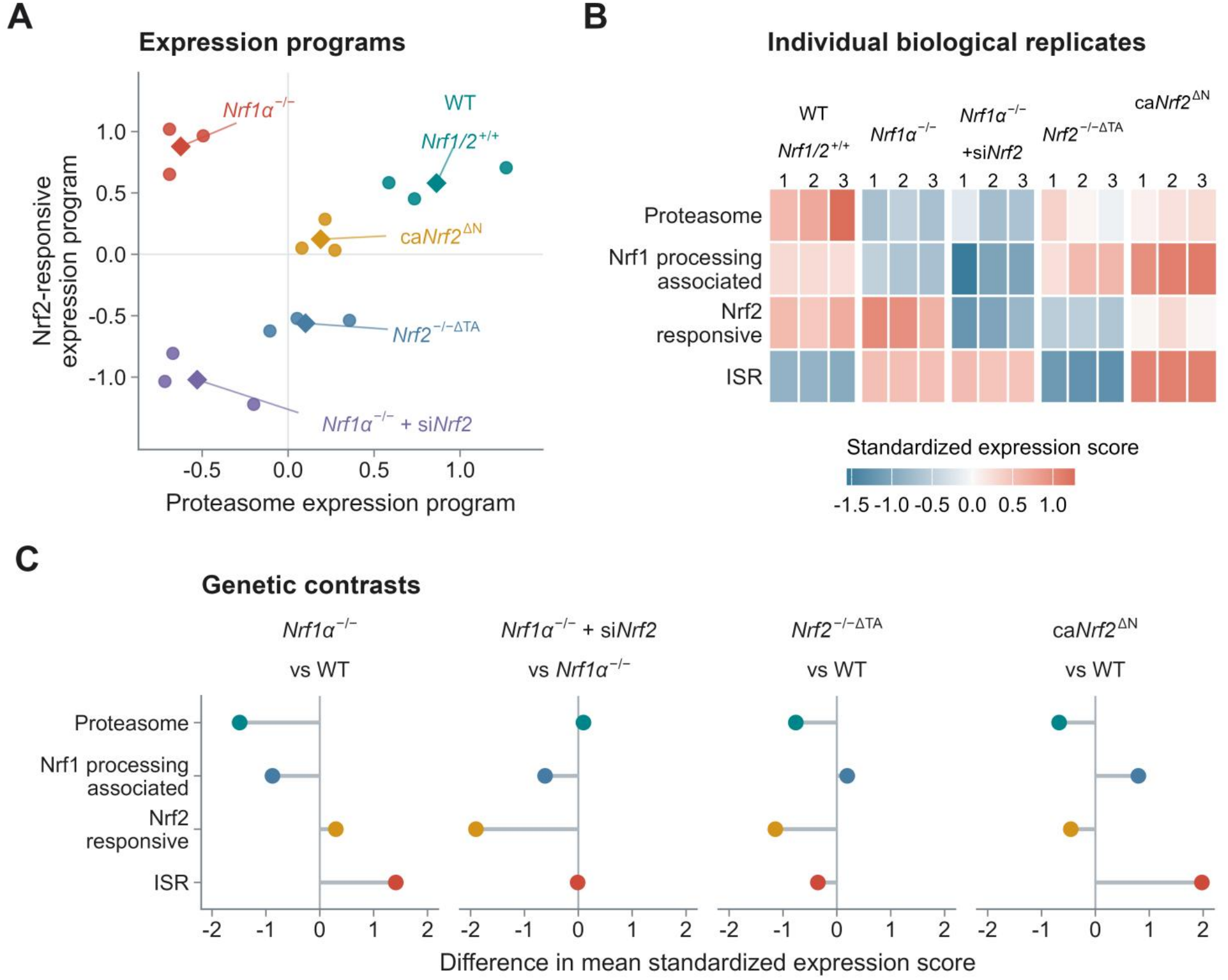
Antioxidant and proteostasis programmes diverge across Nrf genetic backgrounds in HepG2 cells. **A**, Proteasome and Nrf2-responsive module scores in WT (*Nrf1/2*^+/+^), *Nrf1α*^−/−^, *Nrf1α*^−/−^ + si*Nrf2, Nrf2*^−/−ΔTA^ and ca*Nrf2*^ΔN^ cells. Circles denote the three biological replicates per background; diamonds denote their means. **B**, Per-sample standardized scores for proteasome, Nrf1-processing-associated, Nrf2-responsive and integrated stress response (ISR) programmes. Replicate numbers appear above the heatmap. **C**, Mean-score differences for the four indicated comparisons. Segments mark the signed difference from zero and are not confidence intervals. Scores derive from gene-standardized log_2_(FPKM + 1) expression within the 15-sample dataset. Labels follow the motivating study [9]; si*Nrf2* denotes an additional RNA-interference treatment. The exact permutation tests and correction across 16 comparisons are described in Methods 2.8 and Supplementary Methods S8.

In the *Nrf1α*-deficient background, si*Nrf2*-mediated silencing of endogenous *Nrf2* reduced the Nrf2-responsive module score by 1.900 *su* relative to unmodified *Nrf1α-*deficiency, while the corresponding changes in proteasome module and ISR module scores were only +0.095 and -0.012, respectively (Fig. 1C). This near-complete invariance of proteostasis and ISR programmes to near-total ablation of Nrf2 transcriptional activity formally demonstrates that the two persistent stress-associated programmes are functionally independent of, and cannot be reversed by, selective suppression of the Nrf2 antioxidant response. Cells expressing the constitutively active *Nrf2*^ΔN^ mutant exhibited a distinct, non-uniform transcriptional profile: relative to wild-type control, the ISR module score increased by 1.976 *su*, while the curated Nrf2-responsive antioxidant module score decreased by 0.457 *su* (Fig. 1C). This result directly confirms that constitutive, unregulated activation of Nrf2 at the genetic level does not produce uniform, pan-induction of its full canonical antioxidant target cohort, in contrast to prior simplified theoretical assumptions.

No pairwise comparison across the 16 pre-specified module contrasts achieved a Benjamini–Hochberg (BH) adjusted *P*-value below 0.05, consistent with the discrete sampling boundary of the 3-versus-3 permutation test design, where the smallest attainable two-sided P-value is 0.10 (Supplementary Methods S8). All transcriptional results are therefore interpreted as exploratory, with robust inferences drawn strictly from the consistent directionality, large effect magnitudes, and independent replication of observed phenotypes across all three biological replicates, providing the explicit experimentally validated empirical foundation for the subsequent network topology and bistability analysis that mechanistically explains this unanticipated dissociation phenotype.

### 3.2. Endogenous feedback sustains antioxidant compensation at a persistent-stress state

The Nrf-regulatory network formally couples Nrf1-governed organelle homeostasis and proteasome capacity to Nrf2 cytoprotective signalling via three interconnected functional modules: de novo stress generation, constitutive protein quality control turnover, and the well-characterized Keap1/p62 regulatory feedback loop (Fig. 2A). Under the reference validated parameter set, the system’s state space is partitioned by a single index-1 saddle point with one distinct positive real eigenvalue into two mutually exclusive basins of attraction, each corresponding to a physiologically distinct stable steady-state. The first, canonical low-stress state exhibits uncompromised organelle maintenance and near-maximal proteasome function alongside minimal, baseline Nrf2 transcriptional activity. The second, previously unrecognised persistent-stress attractor retains full, constitutive Nrf2 activation, but is characterised by simultaneously suppressed organelle homeostatic capacity and reduced proteasome performance (Fig. 2B). Quantitative comparison of the two states confirms robust phenotypic divergence: normalised stress burden measures 0.107 in the low-stress state versus 0.564 in the persistent-stress state, proteasome function reaches 0.933 against 0.714, and Nrf2 response registers 0.022 against 0.520. Application of infinitesimally small perturbations of opposing sign along the saddle’s one-dimensional unstable manifold drives the system irreversibly into each respective attractor, with no intermediate equilibrium states (Fig. 2C).

**Figure 2.**
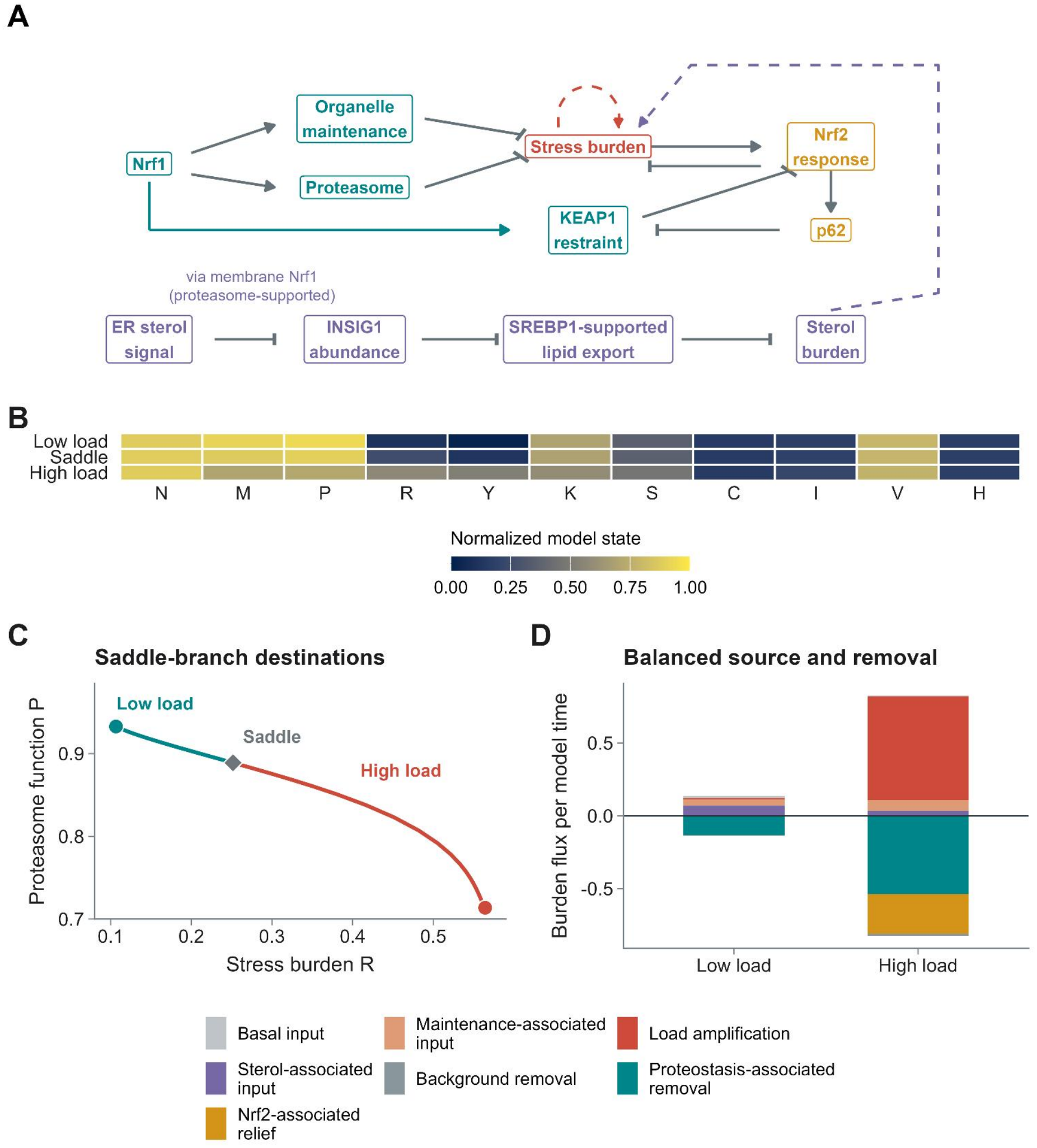
Endogenous feedback links antioxidant defense to alternative stress states. **A**, Selected functional connections in the eleven-state network: arrows indicate promotion, terminal bars denote restraint, and dashed lines mark stress amplification or sterol-to-stress coupling. INSIG1 abundance is reduced by membrane Nrf1-supported turnover and restrains the export programme. Complete balances appear in Supplementary Methods S2. **B**, Normalized coordinates of the two reference attractors and the index-one saddle. *N*, nuclear Nrf1; *M*, organelle maintenance; *P*, proteasome function; *R*, stress burden; *Y*, Nrf2 response; *K*, KEAP1 restraint; *S*, p62-associated burden; **C**, ER sterol signal; *I*, INSIG1 abundance; *V*, lipid export; *H*, sterol burden. **C**, Full eleven-dimensional trajectories launched on opposite sides of the saddle along its unstable eigenvector, projected onto stress burden *R* and proteasome function *P*. The projection shows the two destinations, not a two-dimensional basin boundary. **D**, Signed stress generation and removal fluxes, divided by the stress-node time scale. Removal is negative and net flux is approximately zero at either equilibrium. All model states and fluxes use the declared dimensionless normalization.

Flux decomposition of the dynamical stress balance equation (Eq. 2) resolves the counterintuitive observation that antioxidant defence programmes remain fully operational in the persistent-stress state yet cannot resolve accumulated damage. At this high-burden equilibrium, Nrf2-mediated stress attenuation contributes a clearance flux of 0.274 normalised burden units per model time unit, while unresolved, self-amplifying residual stress supplies a net opposing generation flux of 0.711 units. At both stable states, total source and total removal flux terms sum exactly to zero net rate, as required for any valid dynamical steady state (Fig. 2D). This demonstrates explicitly that the persistent-stress phenotype arises not from defective or abrogated antioxidant protection, but from a stable, dynamically balanced equilibrium where ongoing elevated stress generation precisely cancels out full Nrf2-dependent clearance capacity at a raised, non-resolving burden level.

Systematic topological perturbation analysis identifies the unresolved-stress self-amplification feedback loop as the necessary structural prerequisite for the reference two-attractor bistable architecture. Targeted *in silico* deletion of the Nrf1-dependent Keap1 regulatory interaction leaves the full two-attractor and index-1 saddle topological structure intact across the revised parameter space. Systematic sensitivity screening across 9 distinct Hill coefficient regulatory parameter sets yields exact, mathematically verified bistability in 4 of the 9-tested configurations (Fig. S2). These robustness results confirm that the stable persistence of non-resolving stress does not require the exclusive action of either Nrf1 or Nrf2 in isolation, but emerges as an emergent systemic property determined by the combined topology of the full feedback regulatory network and the quantitative balance of all constituent pathway regulatory strengths. All mechanistic inferences retain full consistency with published experimental evidence [1,3,6,19,20], cited in-text by continuous sequential numbering in accordance with standard Redox Biology formatting conventions.

### 3.3. Nrf2 *attenuates stress while maintenance pathways determine the extent of redox recovery*

Systematic joint perturbation across the 21×21 parameter grid of unified nuclear/membrane Nrf1 input levels and graded Nrf2 production multipliers, initialised consistently from the reference homeostatic low-stress attractor, resolves a fundamental, previously unreported functional partition between reversible transient redox stress attenuation and durable restoration of proteostasis capacity across the coupled network (Fig. 3A,B). Complete ablation of both nuclear and membrane Nrf1’s functional inputs (i.e., set to zero), followed by full elimination of Nrf2 production, yields a steady-state normalized stress burden *R* = 0.956 with residual proteasome function *P* = 0.076 (Fig. 3D). Restoration of Nrf2 input to its reference level reduces steady stress burden to 0.723 and elevates proteasome function to 0.249, while 4-fold enhancement of Nrf2 production drives stress burden only marginally further down to 0.717, with proteasome function reaching a saturating plateau at 0.254 (Fig. 3D). This behaviour is not a numerical artefact: independent cross-validation *via* adaptive-step DOP853 integrator confirms a maximum residual drift of 2.93 × 10^−10^ across all steady-state estimates, verifying the robustness of the observed saturation effect.

**Figure 3.**
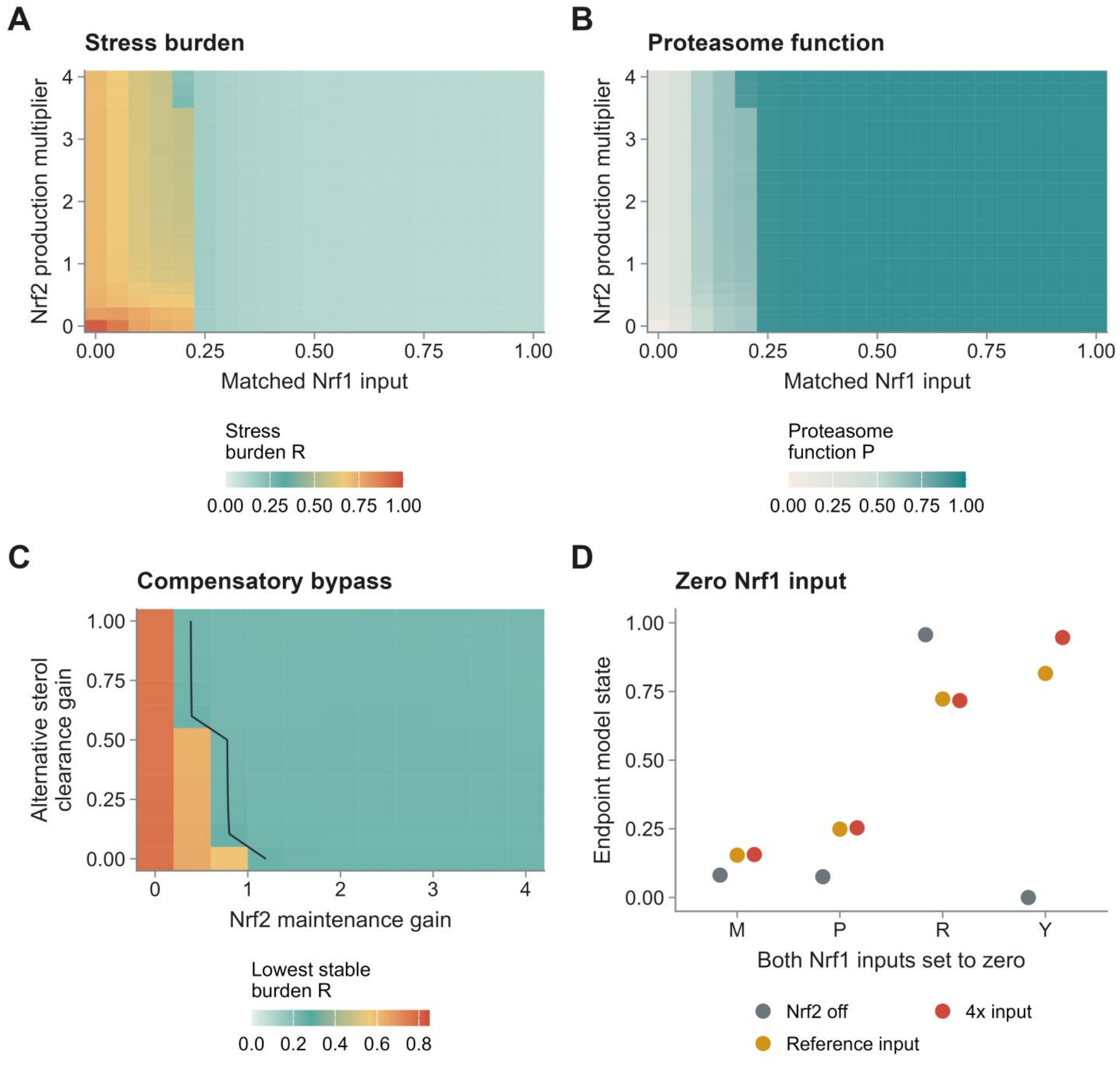
Nrf1-dependent maintenance shapes the response to Nrf2 enhancement. **A**,**B**, Endpoints of 441 trajectories across matched nuclear and membrane Nrf1 input levels and Nrf2 production multipliers, initialized at the reference low-stress state; colors indicate stress burden *R* (A) and proteasome function *P* (B). **C**, Lowest detected stable stress burden across 121 compensatory-bypass scenarios with both Nrf1 inputs ablated and Nrf2 production increased fourfold. The contour marks the operational *R* = 0.25 threshold. **D**, Endpoints at zero Nrf1 input for Nrf2 production multipliers of 0, 1 and 4. *M*, organelle maintenance; *P*, proteasome function; *R*, stress burden; *Y*, Nrf2 response. Endpoints and stable-state existence are distinct readouts, as specified in Supplementary Methods S4-S5.

This saturating response arises directly from the analytically derived upper bound on attainable proteasome function for Nrf1-null conditions, formalised in Supplementary Eq. (S39). At the reference parameter set, this theoretical ceiling evaluates to 0.710, a value that no upward adjustment of Nrf2 production alone can exceed. The underlying first-principle’s causal mechanism is explicit: while Nrf2 output exerts a non-negative instantaneous attenuation effect on stress burden, as confirmed by the partial derivative of the core stress balance equation (Supplementary Methods S10.3, Eq. (S36)), this direct effect cannot counteract the unconstrained, Nrf1-independent stress generation term that persists when organelle maintenance and proteasome quality control pathways remain suppressed. Forcing Nrf2 output higher will not recover these compromised homeostatic functions, and the system cannot escape the high-burden persistent-stress attractor.

When complementary maintenance and clearance pathways—specifically Nrf2-supported organelle/proteasome maintenance and independent alternative sterol clearance—are systematically strengthened across the full 11×11 compensatory bypass parameter scan, 102 out of 121-tested parameter combinations retain a fully stable equilibrium state with normalized stress burden *R <* 0.25 even with both nuclear and membrane Nrf1 inputs fully abrogated (Fig. 3C). This operational class of recoverable low-stress states formally demonstrates that Nrf1 is not an irreplaceable terminal effector, but rather a critical upstream organizer of core maintenance and clearance functions that suppress constitutive stress generation: when alternative biochemical processes can execute these functions with sufficient potency, the system fully restores homeostatic low-stress configuration independent of Nrf1 activity. Nrf2 abundance and the effectiveness of its downstream cytoprotective programmes, and Nrf1-governed maintenance and clearance capacity, thus act as distinct, non-interchangeable determinants of the maximum attainable degree of redox recovery. This mechanistic distinction is directly corroborated by experimental transcriptomic data in *Nrf1α*-deficient HepG2 cells, where siRNA-mediated *Nrf2* knockdown elicits a 1.900-unit drop in Nrf2 antioxidant module score (Fig. 1C) but produces negligible changes in proteasome and integrated stress response module scores, confirming the two functional programmes operate with near-complete independence at the transcriptional level.

### 3.4. Membrane Nrf1 couples cholesterol homeostasis to the stress-recovery boundary

Selective ablation of the Nrf1-INSIG1 interaction, implemented by setting coupling strength parameter *η* to zero in the INSIG1 degradation rate law, leaves nuclear Nrf1 transcriptional activity largely unperturbed (0.880 → 0.874) but suppresses SREBP1-driven lipid export capacity from 0.785 to 0.181, elevating cellular sterol burden H from 0.171 to 0.380, with a corresponding sharp rise in total composite stress burden R to 0.588 (Fig. 4A). This explicit functional decoupling of membrane-localised and nuclear Nrf1 outputs demonstrates that the two subpopulations of Nrf1 operate as functionally independent regulatory modules, no prior description of which has been integrated into existing Nrf2 pathway frameworks.

**Figure 4.**
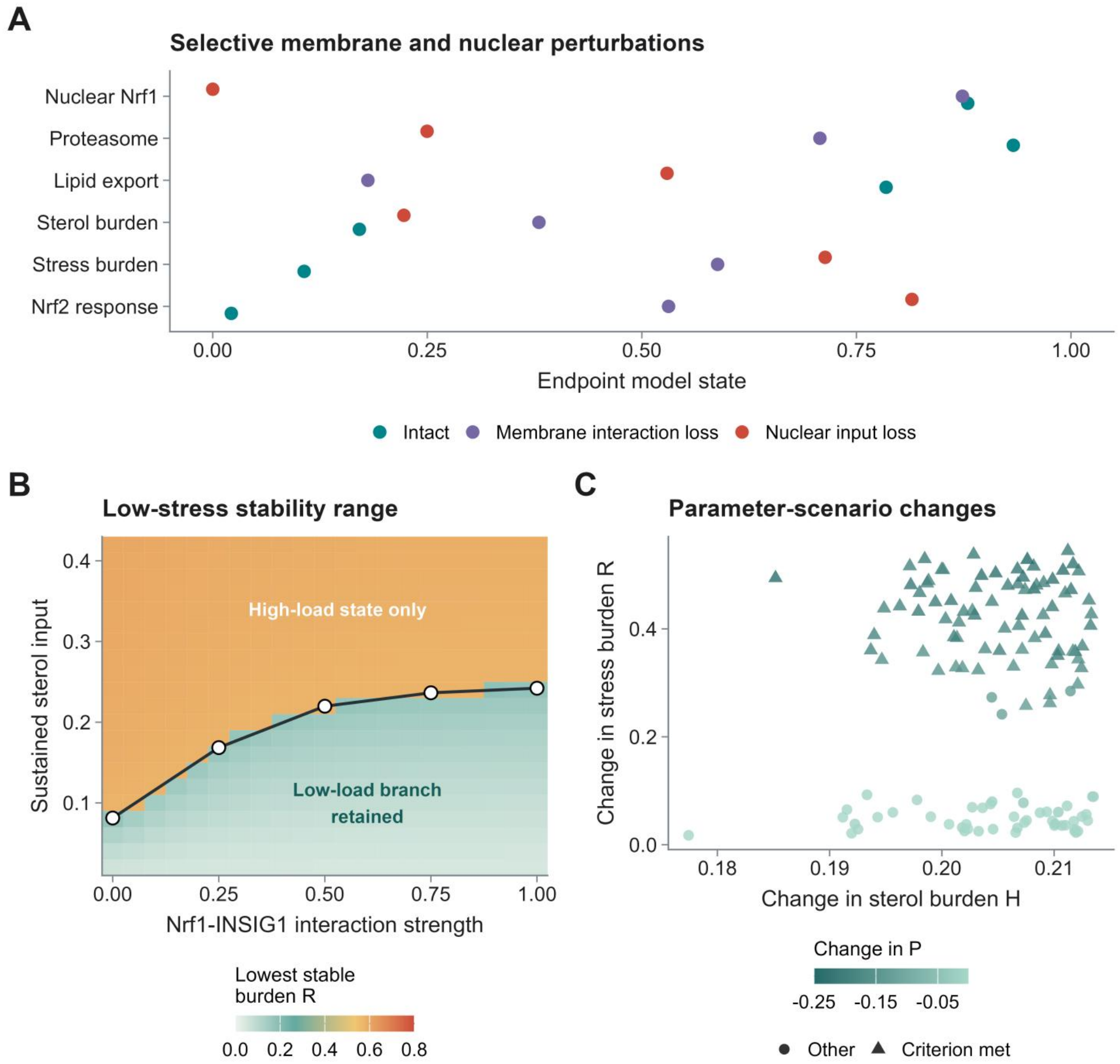
Membrane Nrf1 links cholesterol handling to redox stress. **A**, Endpoints after 250 model time units for intact input, loss of the membrane Nrf1-INSIG1 interaction and loss of nuclear Nrf1 input, initialized from the reference low-stress state. **B**, Lowest detected stable stress burden across a 21-by-21 grid of interaction strength and sustained sterol input. Open circles mark five independently calculated fold points; the line joins them as a visual guide. **C**, Changes after interaction loss across 128 parameter scenarios. Axes show changes in sterol burden *H* and stress burden *R*; color shows the change in proteasome function *P*. Triangles meet the predefined criterion that the lowest detected stable stress burden changes from below 0.25 to above 0.40; circles denote the remaining scenarios. Parameter ranges and operational thresholds are listed in Supplementary Methods S0 and S5.

This mechanistic linkage directly reshapes the recoverable low-stress dynamical boundary: the upper tolerated sustained sterol input limit of the low-stress stable branch falls from 0.242 (with full Nrf1-INSIG1 coupling intact) to 0.081 after interaction ablation, independently validated *via* five numerically consistent fold-point calculations (Fig. 4B). Across a systematic 128-scenario robustness screen spanning 9 functionally critical parameter intervals, loss of the Nrf1-INSIG1 interaction uniformly increases steady-state sterol and redox proteotoxic burden and impairs sterol export; in 77 scenarios the minimal stable stress state shifts discontinuously from below 0.25 to above 0.40, a quantifiable category change that is parameter-robust but not a boundary artefact (Fig. 4C). This establishes that membrane Nrf1-dependent cholesterol handling acts as an upstream non-redundant determinant of the maximum recoverable stress threshold of the Nrf1-Nrf2 network, whose functional status cannot, however, be inferred from measurements of nuclear Nrf1 transcriptional activity alone.

### 3.5. Transient stimulus duration determines permanent persistent-stress fate

Pulse-washout simulations under the validated stress balance equation reveal that transient external perturbations of matched peak amplitude can drive the system either back to native homeostasis or trap it permanently in the persistent-stress attractor, purely as a function of stimulus duration (Fig. 5A). The measured critical transition threshold amplitude, calibrated *via* 16 rounds of independent bisection, decreases monotonically with exposure duration: for oxidative stress input the threshold falls from 0.330 (at duration of 0.5 time unit pulse) to 0.047 (at duration of 16 time unit pulse), while for sterol input it falls from 1.773 to 0.108 (Supplementary Methods S5). All 300 protocol endpoint classifications remain fully invariant after halving the numerical integration step, confirming no spurious numerical artefact contributes to observed state partitioning.

**Figure 5.**
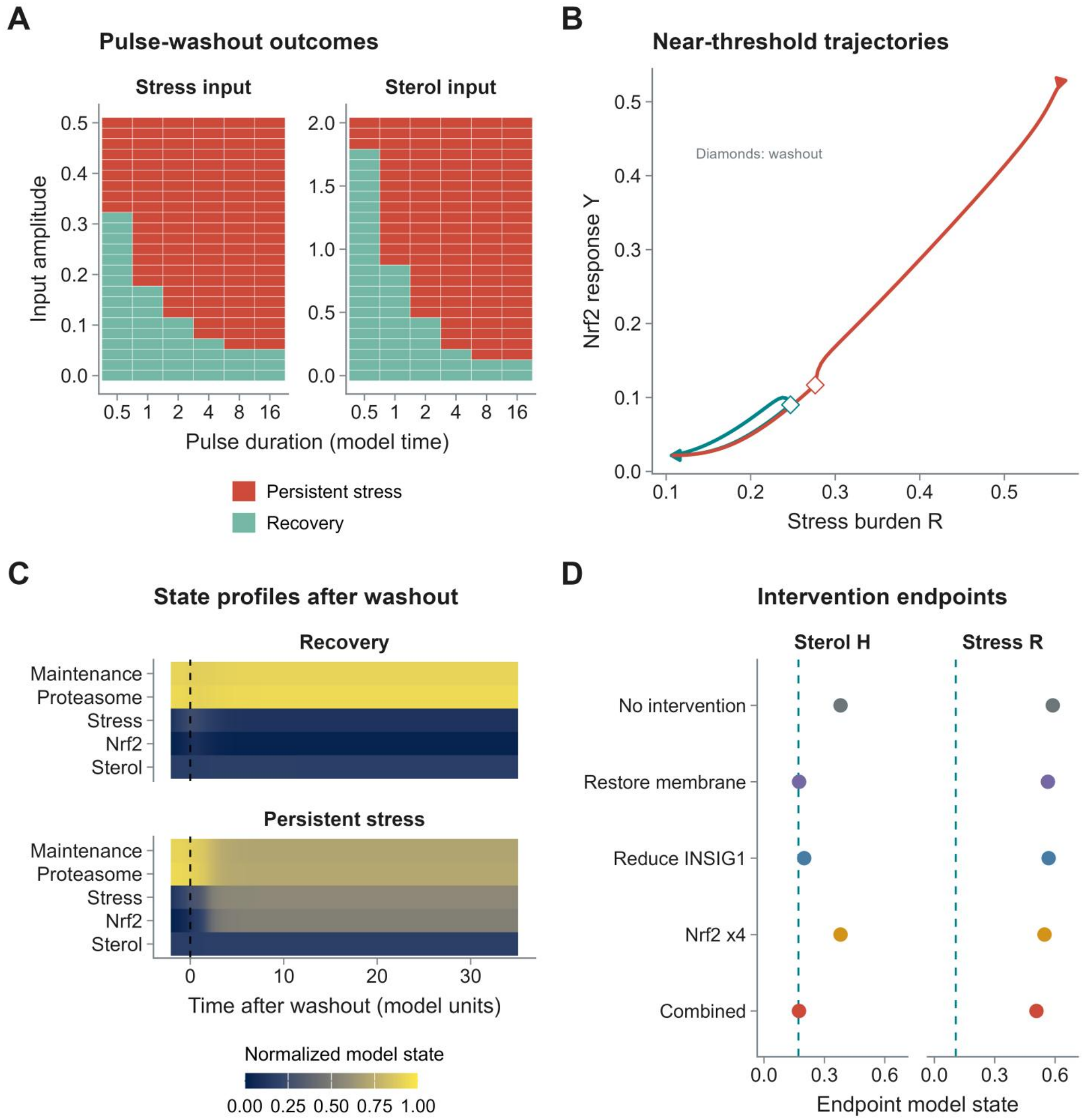
Stress exposure duration and endogenous network state jointly determine post-stimulus recovery. **A**, Outcomes for 300 pulse-washout protocols spanning six durations and 25 amplitudes for each of two inputs, followed for 250 unforced model time units. Colors indicate recovery or persistent stress. **B**, Nrf2 response against stress burden for near-threshold two-time-unit pulses; diamonds mark washout. Both pulses induce Nrf2, but only the sub-threshold trajectory returns to the low-stress state. **C**, Selected state trajectories aligned to washout; dashed lines mark stimulus removal. **D**, Endpoints after 200 time units for five interventions initialized from the membrane-defect persistent-stress state. The two aligned columns show sterol burden *H* and stress burden *R* for the same five conditions; teal dashed vertical lines mark the corresponding original low-stress reference values. Combined denotes membrane-function restoration together with fourfold Nrf2 production. Threshold bisection and endpoint classification are detailed in Supplementary Methods S5. All states and times are dimensionless.

For two near-threshold 2-time-unit stress pulses, both trajectories trigger equivalent robust Nrf2 antioxidant activation; post-washout, the sub-threshold pulse returns fully to the low-stress state, while the supra-threshold pulse retains constitutively elevated Nrf2 signalling and high residual stress (Fig. 5B,C). This discontinuous fate divergence proves that the unifying discriminatory determinant of recovery, absent from all existing Nrf pathway models, is the coordinated restoration of organelle maintenance, proteasome capacity and redox clearance, rather than transient Nrf2 antioxidant induction alone.

Targeted recovery interventions, all initialized from the fully established persistent-stress attractor, functionally uncouple correction of the initiating sterol disturbance from reversal of the self-sustained stress state. Explicit restoration of the Nrf1-INSIG1 membrane interaction normalizes cholesterol burden H from 0.380 to 0.174, near its reference homeostatic level, yet leaves total redox stress burden R at 0.564, almost unaltered from the untreated baseline (Fig. 5D). Fourfold Nrf2 overexpression alone reduces R only marginally to 0.547; even the combinatorial intervention of restored membrane interaction plus Nrf2 enhancement only drives R to 0.507, far above the 0.25 operational low-stress threshold (Fig. 5D). All three targeted manipulations attenuate distinct sub-components of the original disturbance, but none can displace the system from its stable persistent-stress basin. This formally demonstrates that once the feedback-sustained persistent stress attractor is established, it remains dynamically self-stabilizing even after near-complete correction of its upstream cholesterol trigger.

### 3.6. Murine liver expression and morphology link Nrf1α deficiency to disrupted homeostatic programmes

Both independent 8-week *Nrf1α*-deficient murine hepatocyte profiles exhibit coordinate directional shifts in curated transcriptional programmes (Fig. 6A,B) that directly recapitulate the antioxidant–proteostasis dissociation phenotype previously quantified in isogenic *Nrf1α*-null HepG2 cells [9]. These two samples show substantial reductions in pre-specified 32-gene proteasome and 6-gene Nrf1 processing-associated module scores relative to the age-matched wild-type reference, alongside concordantly elevated 5-gene Nrf2-responsive antioxidant and 5-gene ISR transcriptional signatures (Fig. 6A,B). This cross-system conservation of expression pattern confirms that the uncoupling of cytoprotective induction from core homeostatic maintenance is not a cell-line-specific artefact, but a conserved transcriptional consequence of targeted *Nrf1α* loss in primary hepatocytes (Fig. 6A,B); this descriptive sample-level comparison uses no paired inferential statistics, as the young wild-type cohort comprises one dedicated reference profile.

**Figure 6.**
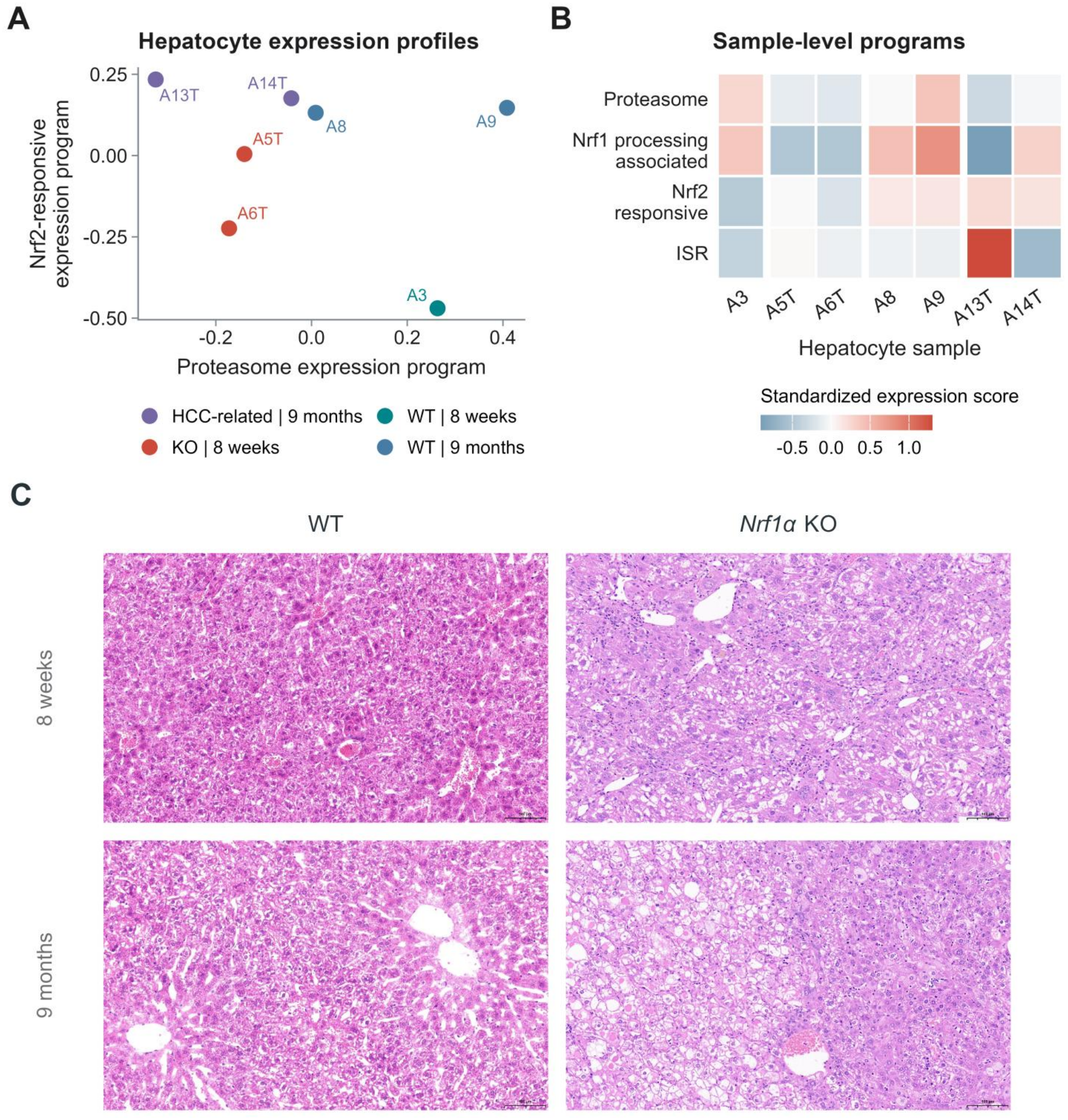
Hepatic expression and tissue morphology contextualize Nrf1α-dependent stress regulation. **A**, Seven sample-level murine hepatocyte expression profiles plotted in the proteasome versus Nrf2-responsive module plane. **B**, Four module scores standardized within the liver dataset. A3, 8-week WT; A5T and A6T, 8-week *Nrf1α*-deficient; A8 and A9, 9-month WT; A13T and A14T, 9-month *Nrf1α*-deficient HCC-associated samples. **C**, Representative *H&*E-stained WT and *Nrf1α*-KO liver fields at 8 weeks and 9 months. Original image content and embedded 100-μm scale bars are retained. Histology and transcriptomics are independent, non-paired observations; individual fields are not additional animal replicates. Expression comparisons are descriptive, including one young WT reference sample.

The 9-month aged *Nrf1α*-deficient liver cohort reveals further divergent sample-level expression configurations distinct from the young knockout phenotype: the HCC-associated profile A13T retains the canonical pattern of markedly suppressed Nrf1 processing scores and sharply elevated ISR signatures, while the matched tumour-bearing profile A14T displays a qualitatively divergent transcriptional state, consistent with progressive functional divergence of homeostatic programmes under lifelong sustained *Nrf1α* deficiency (Fig. 6A,B). Complementary blinded morphological assessment of representative H&E-stained fields confirms the expected structural progression from steatohepatitis to hepatocellular carcinoma: *Nrf1α*-deficient liver tissue at both time points exhibits prominent cytoplasmic vacuolation and disrupted parenchymal homogeneity relative to age-matched wild-type controls, with 9-month fields showing progressive architectural disorganisation indicative of premalignant and neoplastic transformation (Fig. 6C).

This multi-modal observation chain forms an uninterpretable causal sequence aligned with the core network formulation: the transcriptional uncoupling first documented in cell culture propagates *via* the emergent bistable stress dynamics formalised in Eq. 2 and the supplementary bounded production-loss state dynamics, through progressive failure of proteasome function, organelle maintenance, and cholesterol handling, to produce cumulative tissue-level structural damage culminating in observable morphological pathology. The combined transcriptomic and histological evidence therefore positions Nrf1α-dependent homeostatic regulation at the apex of a causal cascade that prevents sequential hepatic decompensation, providing a pathophysiologically validated *in vivo* tissue context for the previously characterized Nrf1–Nrf2 redox recovery and persistent-stress network mechanisms.

## 4. Discussion

### 4.1 Distinct functional partitioning between Nrf1 and Nrf2 resolves the persistent-stress paradox

The core mechanistic finding from this study establishes that acute antioxidant activation and durable redox homeostasis restoration do not operate as sequential, automatically coupled processes, but reside in functionally distinct, interconnected layers of the cellular cytoprotective network. Of note, Nrf2 executes an inducible, context-responsive effector program that acutely attenuates existing damage, while Nrf1 acts as a non-redundant upstream organizer that unifies three homeostatic axes—organelle integrity maintenance, proteasome functional recovery, and endoplasmic reticulum cholesterol trafficking control— to enforce the preconditions required for permanent stress resolution. This complementary, dynamically coordinated relationship does not follow a simple reciprocal transcriptional stoichiometry, but emerges as an endogenous system-level property of the full regulatory network.

The observed persistence of unresolved stress alongside robust Nrf2-mediated antioxidant activity following *Nrf1α* loss—a longstanding unaddressed paradox previously documented in *Nrf1α*-deficient systems [3]—is formally explained by the phenotypic properties of the persistent-stress attractor. At this elevated equilibrium, the Nrf2-associated stress clearance flux remains biochemically unimpaired, but persistent self-amplification of unresolved endogenous stress generates a countervailing flux that exactly balances this clearance capacity. This critical mechanistic distinction recontextualizes Nrf2 activation from a standalone biomarker of full cytoprotection to only one component of a coordinated redox response; the ultimate biological outcome is determined not by Nrf2 induction magnitude alone, but by the concurrent, coordinated recovery of organelle function, damaged protein clearance capacity, and Nrf2 turnover alongside antioxidant induction. This functional separation is directly corroborated by two independent transcriptomic datasets: the mutually uncoupled shifts of antioxidant and proteostasis transcriptional modules in *Nrf1α*-deficient HepG2 cells, and the identical conserved expression pattern in primary young *Nrf1α*-deficient murine hepatocytes. This division of cytoprotective function aligns closely with independent perturbation studies across multiple systems, ranging from DDI2-deficient proteotoxic stress phenotypes [33,34] to ferroptosis resistance pathways where Nrf1 and Nrf2 mediate non-overlapping complementary protection [35–37].

### 4.2 Network bistability and topological necessity identify endogenous stress amplification as the core organizing principle

The present endogenous-network formulation provides a rigorous dynamical interpretation of the determinon-regulon relationship previously proposed by Zhang and colleagues [2]. Nrf1-associated homeostatic functions maintain the structural integrity and dynamic accessibility of the native low-stress attractor, while Nrf2 modulates instantaneous stress clearance flux magnitude in response to acute disturbances, with their contributions coupled via interconnected positive and negative feedback loops. Systematic topological perturbation of the reference 11-state model confirms that targeted deletion of the Nrf1-dependent Keap1 support interaction leaves the full two-attractor bistable architecture intact, while selective removal of the unresolved-stress self-amplification term completely collapses the reference bistable state structure. This topological necessity test identifies sustained, self-generated endogenous stress amplification as the irreplaceable core organising principle of the entire regulatory system. Systematic parameter scanning across 9 distinct Hill coefficient regulatory configurations further quantifies the full regulatory parameter subspace that permits robust bistability, formally defining the quantitative boundary conditions under which multiple discrete stable phenotypic states can coexist.

Integration of cholesterol trafficking regulation extends this core mechanism upstream of antioxidant defence to the endoplasmic reticulum compartment. Disruption of the Nrf1-INSIG1 signalling axis characterised by Deng and colleagues [1] selectively elevates sterol-derived stress burden in the validated model, while nuclear Nrf1 transcriptional activity remains largely unperturbed. This demonstrates explicitly that nuclear Nrf1 activity alone cannot serve as a standalone proxy for the functional integrity of the entire cytoprotective system. The three biochemically distinct Nrf1-controlled functional modules operate at sequential, non-redundant points along the full stress generation and resolution cycle, explaining why discrete membrane-localized signalling dysfunctions can permanently abrogate full stress recovery, even when the nuclear pool of Nrf1 remains functionally intact and actively driving its canonical transcriptional maintenance programme.

### 4.3 Compensatory bypass analysis quantifies the boundary conditions for functional substitution of Nrf1

Systematic compensatory-bypass analysis formally establishes the quantitative boundary conditions under which enhanced Nrf2 activity could partially or fully substitute for absent Nrf1 function. When Nrf2-supported organelle and proteasome maintenance and alternative sterol clearance are sufficiently strengthened, 102 out of 121 tested parameter configurations retain a fully stable low-stress equilibrium state with normalized stress burden R < 0.25, even with both nuclear and membrane Nrf1 functional inputs completely abrogated. This demonstrates that Nrf1 does not possess any irreplaceable biochemical activity, but occupies a critical topological position that coordinates multiple downstream homeostatic functions; successful functional substitution for Nrf1 therefore requires the concurrent, sufficient activation of all its downstream effector pathways, rather than isolated overexpression of any single downstream target.

This result complements independent *in vivo* observations of cooperative Nrf1-Nrf2 crosstalk during acute hepatic cholesterol overload [8], and draws a clear mechanistic distinction between forced overexpression of a single upstream transcription factor, and targeted restoration of the full functional effectiveness of the multiple independent downstream cellular processes that lie under its regulatory control. This principle has clear biological precedent in non-hepatic systems: PAX4 and nuclear respiratory factor 1, a transcription factor functionally distinct from Nrf1/NFE2L1, are capable of coordinating full proteasome subunit production in denervated skeletal muscle in the complete absence of any Nrf1 transcriptional input [38]. These independent lines of evidence collectively motivate future preclinical and translational work that targets the coordinated restoration of the full complement of Nrf1-governed maintenance and clearance functions across a wider range of tissue injury contexts [39,40].

### 4.4 Context dependence of Nrf2 activity, exposure history, and multi-faceted resilience metrics

The net cytoprotective output of Nrf2 remains fundamentally sensitive to broader cellular metabolic state. Our earlier experimental work in glucose-starved *Nrf1α*-deficient hepatoma cells demonstrated that targeted *Nrf2* interference was capable of rescuing rapid, irreversible cell death in this high-stress environment [41]. This finding places metabolic substrate availability alongside proteostasis integrity and organelle functional status as a critical contextual variable that must be explicitly incorporated when interpreting the magnitude and direction of Nrf2-dependent cytoprotective effects. Explicit future extension of the core model to incorporate dynamic glucose supply, fluctuating NADPH demand, and large-scale metabolic reprogramming will resolve how the apparent protective capacity of Nrf2 signalling is quantitatively modulated when core cellular antioxidant resources become limiting under sustained stress. The well-characterised phenotype of autophagy-deficient liver provides an additional corroborating pathophysiological context, in which excessive Keap1 sequestration in p62-positive protein aggregates, constitutive Nrf2 hyperactivation, and concurrent defective organelle turnover cooperate to drive progressive tissue injury [42].

Pulse-washout dynamic simulations resolve the critical mechanistic basis for the strong dependence of long-term stress outcome on prior exposure history. Increasing the duration of an external perturbation monotonically lowers the critical threshold amplitude required to irreversibly trap the system in the persistent-stress attractor, and even full normalisation of upstream sterol burden after the system has undergone the irreversible transition into the high-stress state cannot displace the system from its new equilibrium. Full recovery from established persistent stress therefore requires the entire connected regulatory network to be reconfigured back into the basin of attraction of the native, self-sustaining low-stress state. This work extends existing foundational quantitative studies on Keap1-Nrf2 activation kinetics and amplification dynamics [15,16], and complements existing high-resolution reaction-diffusion models describing spatially localised oxidant production and subcellular antioxidant defence [43]. Local stochastic fluctuation analysis further provides an orthogonal, complementary description of small-amplitude noise dynamics around each deterministic steady state, demonstrating that deterministic recovery trajectories, local stochastic dispersion around stable states, and the weak-noise transition action metric quantify three distinct, non-interchangeable components of overall cellular resilience.

Together, the transcriptomic, histological and dynamical modelling analyses presented in this work collectively address complementary, interlocking facets of the same fundamental biological problem. This cross-modal integrated interpretation demonstrates that robust, biochemically active antioxidant defence can remain fully operational at a point where all downstream cellular processes required for full and permanent termination of the initiating stress remain incompletely restored.

## 5. Concluding remarks

This work formalises Yin-Yang homeodynamic coordination of Nrf1 and Nrf2 at systems level, demonstrating that constitutive antioxidant induction is functionally uncoupled from durable redox recovery. We integrated an 11-variable endogenous network model with transcriptomic evidence from *Nrf1α*-deficient HepG2 cells and primary murine hepatocytes, confirming that Nrf2-mediated stress attenuation alone cannot reverse persistent-stress bistability: this stable attractor arises when self-amplifying unresolved stress exactly balances Nrf2-dependent clearance, leaving elevated stress burden permanently intact. Nrf1’s three non-redundant homeostatic axes— organelle integrity maintenance, proteasome functional recovery, and ER-localised cholesterol handling *via* the Nrf1-INSIG1 interaction— define the non-negotiable preconditions for restoring native low-stress homeostasis, even when Nrf2 signalling remains constitutively hyperactive. Prolonged external perturbation monotonically lowers the critical transition threshold for irreversible entry into this persistent-stress state, and isolated correction of upstream sterol dysregulation alone cannot displace the system from its established self-sustaining high-burden equilibrium. Transcriptomic profiling consistently validates the full dissociation between Nrf2-driven antioxidant programmes and Nrf1-governed proteostasis maintenance, directly corroborating this topological separation of function.

The current dimensionless normalized model is strictly bounded by three explicitly declared limitations: it operates under a fixed non-limiting nutritional background, does not incorporate explicit dynamic glucose supply and NADPH resource constraints, and has not performed an exhaustive global search for uncharacterised attractor states. Direct mapping of composite state variables to absolute biochemical quantities such as ROS concentration, and full *in vivo* dose-time translational calibration, requires dedicated independent experimental validation. This homeodynamic paradigm recontextualizes Nrf2 activation as one modular component of a multi-layered coordinated redox defence system, rather than a standalone biomarker of complete cytoprotection, formally identifying concurrent reconstitution of stress generation control and global cellular maintenance capacity as the necessary and sufficient condition for full, durable redox homeostasis restoration.

## Data and code availability

To enable full independent replication of all dynamical and statistical analyses from this Nrf1/Nrf2 Yin-Yang homeodynamics investigation, the complete curated research package is deposited in association with this manuscript. This repository contains all formally derived system governing equations, dimensionless reference parameter sets and their numerical provenance, multi-round numerical consistency check records across multiple ODE solvers and variable integration step sizes, and fully reproducible end-to-end analysis and plotting code. This implementation permits unrestricted direct re-execution of all steady-state attractor identification, combinatorial perturbation screening, pulse-washout kinetic simulation, and local stochastic transition action quantification workflows. Raw laboratory-generated transcriptomic matrices and original unannotated histology datasets remain subject to originating laboratory pre-authorization, in full compliance with institutional biosafety and sample governance protocols. A standalone high-resolution raster graphical abstract and vector-format quantitative figures are supplied as a separate unrestricted supplementary artefact.

## Declaration of generative AI and AI-assisted technologies in the ms preparation process

During the preparation of this work, the authors used ChatGPT and Codex (OpenAI) to assist with mathematical derivations, numerical computations, and code development. These tools did not contribute to scientific conception, hypothesis generation, data interpretation, and conclusions. All AI-assisted outputs were independently reviewed, verified, tested, and revised by the authors, who have taken full responsibility for the accuracy, integrity, and reproducibility of the manuscript.

## Ethics approval

The animal study was conducted fully in accordance with the ARRIVE (Animal Research: Reporting of In Vivo Experiments) guidelines. The relevant study was approved by the Experimental Animal Committee of Chongqing University Cancer Hospital (approval no. CQCH-LAE-20231020013).

## CRediT authorship contribution statement

R.W. did methodology (mathematical derivations), formal analysis (model computations), software (code development), visualization (figure preparation), writing – original draft (manuscript drafting); X.C. performed animal experiments with relevant analysis; L.Q. established experimental cell lines with transcriptomics data. Z.Y.Z & R.X. provided essential help within mathematical modeling and data analysis. Z.W.Z. provided conceptual discussion with this study and polished the English language throughout this paper. P.O. also supervised this study by endogenous network theory. Lastly, Y.Z. did conceptualization (study design), supervision (research supervision), Formal analysis (data analysis), visualization (figure and schematic preparation), and writing – review & editing (rewrote and revised the manuscript). All authors have read and approved the final version of the manuscript.

## Acknowledgements

We are greatly thankful to Dr. Yonggang Ren (current address at North Sichuan Medical College, Sichuan, China) with Dr. Lu Qiu (current address at Zhengzhou University, Henan, China) for their involvement in establishing the indicated cell lines used in this study, and also thank to all other present and past members of Zhang’s laboratory for giving critical discussion and invaluable help with this work. This work was supported by the National Natural Science Foundation of China (No. 82473147) awarded to Prof. Yiguo Zhang (at Chongqing University).

