## Supplementary Information for "Yin-Yang homeodynamics of Nrf1 and Nrf2: coupling antioxidant defense to redox recovery bistability under persistent stress"

[1]Lab of Cell Biochemistry and Topogenetic Regulation, College of Bioengineering & Chongqing Academy of Medical Sciences, Chongqing General Hospital, Faculty of Medicine, Chongqing University, Chongqing 400044, China
[2]Department of Laboratory Medicine, Chongqing Center for Clinical Laboratory, Chongqing Academy of Medical Sciences, Chongqing General Hospital, School of Medicine, Chongqing University, Chongqing 401147, China
[3]Department of Pathology, the First Affiliated Hospital of Zhengzhou University, Zhengzhou, Henan, 450052, China,
[4]College of Biomedical Engineering, Sichuan University, Chengdu, Sichuan 610044, China
[5]Laboratory of Neuroscience, Institute of Cognitive Neuroscience and School of Pharmacy, University College London, 29-39 Brunswick Square, London WC1N 1AX, UK
§ equal contribution to this work

### Supplementary methods and computational analyses relative to this work are shown below:

#### S0 Parameter definitions and numerical specification

This supplement defines the eleven network coordinates and all 74 dynamical settings, derives the governing equations, and describes the numerical and expression analyses. S0 is the parameter catalogue; S2 contains the complete balances; S10-S11 derive the main-text equations; S12 connects parameter construction to the additional computational experiments.

Intact input multipliers equal one by normalization. Zero optional gains specify the reference topology. The remaining coefficients, thresholds and relative times define a reproducible reference scenario. Published studies determine the regulatory functions and directions; the numerical values are reference assignments explored by sensitivity analysis, rather than kinetic estimates from a fitted time series.

At fixed regulatory inputs, a node approaches $z_i^* = \alpha_i/(\alpha_i + \beta_i)$ with rate $(\alpha_i + \beta_i)/\tau_i$. An endpoint therefore constrains a rate ratio; turnover or time-course information additionally constrains response speed. S12 gives operating-point reconstruction examples and a multi-condition parameterization procedure.

Figure S7 separates normalized operating levels from turnover scales and illustrates the coefficient reconstruction in S12.1.

The derivation proceeds from parameter definitions to the eleven-state drift, from the drift to its attractors, and from each attractor to local fluctuation geometry and transition costs. Figures S9-S10 connect the landscape interpretation to the covariance and path calculations; S10 provides the underlying identities.

##### *S0.1 Response and control*

| Symbol | Value | Definition and reference setting |
|---|---|---|
| $n$ | 4 | Shared Hill exponent; steepness of the seven-function regulatory gates.<br>Reference nonlinearity; evaluated at 2, 3 and 4. |
| $K_h$ | 0.42 | Half-response input for the shared Hill gate.<br>Reference threshold; evaluated at 0.35, 0.42 and 0.50. |
| $\gamma_\mathrm{n}$ | 1 | Multiplier of effective nuclear Nrf1 establishment.<br>Intact input normalized to 1; joint-input grid 0 to 1; loss 0. |
| $\gamma_\mathrm{e}$ | 1 | Multiplier of ER-associated Nrf1 availability.<br>Intact input normalized to 1; matched joint-input grid 0 to 1; loss 0. |
| $\gamma_2$ | 1 | Multiplier of the entire Nrf2 establishment term.<br>Reference input 1; joint-input grid 0 to 4; enhanced input 4. |
| $q_2$ | 1 | Conversion of Nrf2 state into effective output.<br>Reference normalization; Nrf2-response deletion uses 0 together with zero production. |
| $a_N$ | 0 | Optional Nrf1 self-activation gain.<br>Disabled in the reference network; enabling it requires full-state root finding. |

| Symbol | Value | Definition and reference setting |
|---|---|---|
| $a_{2N}$ | 0 | Optional Nrf2-to-Nrf1 establishment gain.<br>Disabled in the reference network; no nonzero scan reported. |
| $\chi$ | 1 | Switch for sterol feedback into nuclear Nrf1 and stress.<br>1 for the coupled network; 0 for the updated seven-function control. |

#### *S0.2 Maintenance and stress*

| Symbol | Value | Definition and reference setting |
|---|---|---|
| $b_M$ | 0.05 | Basal organelle and redox maintenance input.<br>Positive basal maintenance; ensemble range 0.02 to 0.20. |
| $b_P$ | 0.05 | Basal proteasome maintenance input.<br>Positive basal maintenance; ensemble range 0.02 to 0.20. |
| $a_{2M}$ | 0.05 | Nrf2-supported organelle maintenance gain.<br>Reference compensation; ensemble 0 to 0.30; joint bypass 0 to 4. |
| $a_{2P}$ | 0.15 | Nrf2-supported proteasome maintenance gain.<br>Reference compensation; ensemble 0 to 0.60; joint bypass 0 to 4. |
| $a_R$ | 1.6 | Gain of saturating unresolved-stress amplification.<br>Reference feedback strength; ensemble 0.8 to 2; deletion 0. |
| $a_{NK}$ | 0.78 | Nrf1-dependent support of functional KEAP1 restraint.<br>Protein-level direction from [3]; gain is a reference setting; deletion 0. |
| $a_{SK}$ | 0.9 | p62-dependent weakening of KEAP1 restraint.<br>Direction from [17,19,20]; reference effective gain. |
| $a_{NS}$ | 0.3 | Nrf1-supported p62-associated clearance gain.<br>Direction from [18]; reference gain, distinct from p62 production. |

#### *S0.3 Sterol and membrane*

| Symbol | Value | Definition and reference setting |
|---|---|---|
| $\eta$ | 1 | NHB2-associated Nrf1-INSIG1 interaction multiplier.<br>Intact interaction 1; deletion 0; membrane grid 0 to 1. |
| $r_C$ | 0.75 | Sterol-dependent attenuation of nuclear Nrf1 input.<br>Direction motivated by [6]; reference gain, not a measured retained fraction. |
| $K_C$ | 0.3 | Half-response ER sterol signal.<br>Reference threshold for the membrane and nuclear retention gates. |
| $c_b$ | 0.12 | Basal target level of the ER sterol signal.<br>Reference offset in the burden-to-ER closure. |
| $\kappa$ | 0.2 | Gain from cellular sterol burden to the ER target.<br>Ensemble 0 to 0.4; decoupling 0; target remains at most 1. |
| $b_I$ | 0.2 | INSIG1 establishment coefficient.<br>Reference input; synthesis-reduction intervention uses 0.04. |
| $d_I$ | 0.08 | Basal INSIG1 loss coefficient.<br>Positive reference turnover coefficient. |
| $k_I$ | 2 | Nrf1-associated INSIG1 turnover gain.<br>Direction from [7]; reference gain; ensemble 1 to 4. |
| $f_u$ | 1 | Weight of proteasome dependence in INSIG1 turnover.<br>Reference fully dependent scenario; alternative 0 removes this dependence. |

| Symbol | Value | Definition and reference setting |
|---|---|---|
| $b_V$ | 0.02 | Basal export-program establishment.<br>Positive reference input. |
| $a_V$ | 0.8 | Maximal additional export establishment released by low INSIG1.<br>Reference gain under the selected nutritional context [7]. |
| $d_V$ | 0.2 | Export-program loss coefficient.<br>Positive reference turnover. |
| $K_I$ | 0.3 | INSIG1 half-response threshold for export restraint.<br>Reference inhibitory threshold. |
| $j_H$ | 0.15 | Constitutive cellular sterol input.<br>Reference input; sustained-input map 0.02 to 0.42. |
| $d_H$ | 0.1 | Basal sterol-removal coefficient.<br>Positive reference clearance. |
| $k_V$ | 0.8 | Export-program contribution to sterol removal.<br>Reference gain; ensemble 0.4 to 1.2. |
| $k_{2H}$ | 0 | Optional Nrf2-supported alternative sterol clearance.<br>Absent in reference; bypass grid 0 to 1. |
| $a_H$ | 0.3 | Sterol-associated stress-source gain.<br>Effective cross-module coupling; ensemble 0.1 to 0.6; deletion 0. |
| $K_H$ | 0.35 | Half-response sterol burden for stress generation.<br>Reference cross-module threshold. |

#### *S0.4 Fixed coefficients appearing in the balance laws*

The following named coefficients reproduce the constants in the original balance equations. Their numerical values remain unchanged.

| Symbol | Value | Definition and reference setting |
|---|---|---|
| $b_N$ | 1.1 | Nuclear Nrf1 establishment before input scaling.<br>Basal establishment in the $N$ equation. |
| $d_N$ | 0.13 | Nuclear Nrf1 loss.<br>Together with nuclear input sets the frozen-state $N$ level. |
| $a_{NM}$ | 1.05 | Nrf1-supported organelle maintenance.<br>Reference $N$-to-$M$ support; direction [1]. |
| $d_M$ | 0.1 | Basal organelle-maintenance loss.<br>Positive maintenance turnover. |
| $d_{RM}$ | 0.48 | Stress-dependent organelle impairment.<br>Reference $R$-to-$M$ loss gain. |
| $a_{NP}$ | 1.15 | Nrf1-supported proteasome maintenance.<br>Reference $N$-to-$P$ support; direction [5,9,10]. |
| $d_P$ | 0.08 | Basal proteasome loss.<br>Positive proteasome turnover. |
| $d_{RP}$ | 0.55 | Stress-dependent proteasome impairment.<br>Reference $R$-to-$P$ loss gain. |
| $b_R$ | 0.012 | Basal stress production.<br>Positive background input in the $R$ equation. |
| $a_{MR}$ | 0.42 | Stress input caused by incomplete organelle maintenance.<br>Multiplies the deficit 1-M. |

| Symbol | Value | Definition and reference setting |
|---|---|---|
| $d_R$ | 0.02 | Background stress relief.<br>Positive clearance independent of $P$ and Nrf2 output. |
| $k_{PR}$ | 1 | Proteasome-associated relief of damaged-protein burden.<br>Unit reference gain for the $P$ times $R$ removal term. |
| $k_{2R}$ | 0.52 | Nrf2-associated stress-relief gain.<br>Reference output-to-relief strength. |
| $b_Y$ | 0.018 | Basal Nrf2 establishment.<br>Small positive input before scaling by $\gamma_2$. |
| $a_{RY}$ | 0.95 | Stress-induced Nrf2 establishment.<br>Reference inducible response gain. |
| $d_Y$ | 0.12 | Nrf2 loss independent of the joint KEAP1-proteasome term.<br>Positive background turnover. |
| $k_{KP}$ | 1.35 | Joint KEAP1 and proteasome contribution to Nrf2 loss.<br>Multiplies $K$ times $P$; joint turnover condition [13]. |
| $b_K$ | 0.18 | Basal establishment of KEAP1 restraint.<br>Positive background support. |
| $d_K$ | 0.1 | Basal loss of KEAP1 restraint.<br>Positive background turnover. |
| $b_S$ | 0.015 | Basal p62-associated production.<br>Positive input independent of Nrf1 and Nrf2. |
| $a_{NS}^{+}$ | 0.35 | Nrf1-supported p62-associated production.<br>Production coefficient; distinct from $a_{NS}$ clearance [18]. |
| $a_{2S}$ | 0.28 | Nrf2-supported p62-associated production.<br>Reference output-dependent production. |
| $d_S$ | 0.34 | Basal p62-associated clearance.<br>Positive background clearance. |
| $\theta_e$ | 0.3 | Basal fraction in membrane Nrf1 availability.<br>The complementary sterol-responsive fraction is 0.70, not an independent parameter. |
| $n_C$ | 2 | Exponent for the ER sterol gate.<br>Fixed reference gate exponent; not varied by the shared $n$ scan. |
| $n_I$ | 4 | Exponent for INSIG1 restraint of the export program.<br>Fixed reference gate exponent; not varied by the shared $n$ scan. |
| $n_H$ | 2 | Exponent for the sterol-to-stress gate.<br>Fixed reference gate exponent; not varied by the shared $n$ scan. |

The numerical constants 0 and 1 in available-range factors and complementary gates define the normalized domain. The factor one half in the diffusion matrix and one quarter in the action follow from the chosen noise normalization; they are derived constants, not fitted gains.

#### *S0.5 Relative node time scales*

The relative times $\tau_i$ specify temporal ordering. The frozen-input relaxation time is $\tau_i/(\alpha_i+\beta_i)$; collective relaxation follows the full Jacobian. The clock experiments in S12.5 vary selected times and compare pulse outcomes.

| Symbol | Value | Associated process |
|---|---|---|
| $\tau_N$ | 1.10 | Effective nuclear Nrf1 activity |
| $\tau_M$ | 1.60 | Organelle and basal redox maintenance |

| Symbol | Value | Associated process |
|---|---|---|
| $\tau_P$ | 1.45 | Effective proteasome function |
| $\tau_R$ | 0.75 | Coupled redox and proteotoxic stress burden |
| $\tau_Y$ | 0.28 | Effective Nrf2 response |
| $\tau_K$ | 0.55 | Functional KEAP1 restraint |
| $\tau_S$ | 0.62 | p62-associated abundance |
| $\tau_C$ | 0.50 | ER free-sterol signal |
| $\tau_I$ | 0.80 | INSIG1 protein abundance |
| $\tau_V$ | 1.50 | SREBP1-supported export program |
| $\tau_H$ | 3.00 | Cellular sterol burden |

#### *S0.6 Inputs and perturbation protocols*

| Quantity | Value or protocol | Meaning and calculation |
|---|---|---|
| $u, u_H$ | 0 in the reference autonomous system | Additional stress and sterol inputs. They are pulsed separately. |
| $A_\mathrm{p}$ | Stress 0 to 0.5; sterol 0 to 2 | 25 evenly spaced amplitudes per input in the displayed pulse grid. |
| $T_\mathrm{p}$ | 0.5, 1, 2, 4, 8, 16 | Six exposure durations per input; 300 protocols in total. |
| Threshold brackets | Stress [0,1]; sterol [0,3] | 16 bisections after checking the upper bracket switches state. |
| Washout duration | 250 | Unforced integration after each pulse. |
| Joint input map | 21 by 21 grid; 250 time units | $\gamma_\mathrm{n} = \gamma_\mathrm{e}$ from 0 to 1; $\gamma_2$ from 0 to 4. |
| Membrane input map | 21 by 21 grid | $\eta$ from 0 to 1; $j_H$ from 0.02 to 0.42. |
| Robustness ensemble | 128 Sobol settings; seed 2201 | Nine parameters varied over the ranges in S0.1-S0.3. |
| Hill sensitivity | 3 by 3 combinations | $n \in \{2,3,4\}$ and $K_h \in \{0.35, 0.42, 0.50\}$. |
| Bypass grid | 11 by 11 combinations | $\gamma_\mathrm{n} = \gamma_\mathrm{e} = 0$, $\gamma_2 = 4$; joint $a_{2M} = a_{2P}$ from 0 to 4 and $k_{2H}$ from 0 to 1. |
| Recovery starting state | 250 time units at interaction loss | Begin at the endpoint reached with $\eta = 0$ from the reference low-stress state. |
| Recovery interventions | 200 time units per intervention | No change; restore $\eta = 1$; set $b_I = 0.04$; set $\gamma_2 = 4$; or restore $\eta = 1$ and set $\gamma_2 = 4$. |
| Ensemble state criterion | $R_0 < 0.25$, $R_1 > 0.40$ | Operational category for the change in the lowest detected stable burden after interaction loss. |

#### *S0.7 Stochastic and numerical settings*

| Symbol or setting | Value or definition | Purpose and scope |
|---|---|---|
| $\varepsilon$ | $10^{-4}$ for probability ellipses | Illustrative scale selected by the coordinatewise boundary criterion in S6. Covariance tables report covariance divided by $\varepsilon$. |
| $\Omega$ | $10^{4}$ for the selected illustration | Inverse illustrative noise scale in the effective-event comparison, defined by $\varepsilon = 1/\Omega$ in S10.6. |
| $\mathbf{D}_0, \mathbf{D}_v$ | $\mathbf{D}_0 = \mathbf{1}_{11}$; $\mathbf{D}_v = \frac{1}{2}\mathrm{diag}(a_i^+ + a_i^-)$ | Isotropic and effective-flux local diffusion conventions. Transition-action calculations use the isotropic convention. |
| Local ellipse | Probability 0.683; squared radius 2.2957489 | Bivariate Gaussian marginal in stress and proteasome coordinates. |

| Symbol or setting | Value or definition | Purpose and scope |
|---|---|---|
| Root search | 1024 reduced starts; 256 full starts | Scrambled Sobol sampling plus 5 and 7 structured starts respectively. Primary seed 20261014; full check 71; repeats 17,71,307. |
| Root solver | Tolerance $10^{-10}$; evaluations 400 or 700 | Reduced or full hybr solves. Accept full drift infinity norm below $10^{-8}$. |
| Root domain and identity | Domain tolerance $10^{-7}$; merge distance $10^{-5}$ | Checks bounded states and removes numerically duplicate roots. |
| Eigenvalue classification | Real-part margin $10^{-7}$ | All below its negative value for stability; positive values above it count unstable directions. |
| Jacobian differentiation | Complex step $10^{-28}$ | Cross-check with central difference $10^{-6}$ at 100 states; check seed 41. |
| Saddle branch perturbation | Magnitude $10^{-5}$; duration 180 | Both signs of the unique unstable eigenvector. |
| Adaptive integration | Relative $2 \times 10^{-9}$; absolute $2 \times 10^{-11}$ | DOP853; maximum step 1; failed solutions raise an error. |
| Fixed-step comparison | 0.05 and 0.025 | Fourth-order Runge-Kutta; eight endpoints independently checked with DOP853. |
| Fold derivative step | $10^{-4}$ | Central differences for the directional curvature and parameter transversality checks. |
| Action horizons and segments | 12/60, 24/120, 48/240, 48/480, 96/960 | Finite-time midpoint discretizations; curved restart at 48/240. |
| Action minimizer | L-BFGS-B; coordinate bounds [0,1] | Function tolerance $10^{-13}$; gradient tolerance $10^{-7}$; history 20; line searches 40. Iteration limits follow the archived stages. |
| Action-gradient check | 20 coordinates; step $10^{-6}$; seed 24 | Central-difference comparison with the analytic discrete gradient. |
| Expression transformation | log2(expression + 1) | Average duplicate genes before transformation; exclude zero-variance genes. |
| Expression standardization | Sample standard deviation; 15 or 7 profiles | Separate scaling within HepG2 and liver datasets. Module means use retained genes. |
| Exact tests and adjustment | 20 allocations; 16 tests; $q$ threshold 0.05 | Two-sided permutation comparisons and Benjamini-Hochberg adjustment; minimum possible exact $P$ is 0.10. |

#### *S0.8 States and mathematical symbols*

| State | Definition | Domain |
|---|---|---|
| $N$ | Effective nuclear Nrf1 activity | [0,1] |
| $M$ | Organelle and basal redox maintenance | [0,1] |
| $P$ | Effective proteasome function | [0,1] |
| $R$ | Coupled redox and proteotoxic stress burden | [0,1] |
| $Y$ | Effective Nrf2 response | [0,1] |
| $K$ | Functional KEAP1 restraint | [0,1] |
| $S$ | p62-associated abundance | [0,1] |
| $C$ | ER free-sterol signal | [0,1] |
| $I$ | INSIG1 protein abundance | [0,1] |
| $V$ | SREBP1-supported export program | [0,1] |
| $H$ | Cellular sterol burden | [0,1] |

| Symbol | Definition |
|---|---|
| $\mathbf{z}, \mathbf{Z}$ | Deterministic state vector and stochastic state vector in the same eleven-coordinate order. |
| $h, h_C, h_I, h_H$ | Shared regulatory Hill gate and the ER-sterol, INSIG1 and burden-specific gates. |
| $o, e, p_u, c_t$ | Effective Nrf2 output, membrane Nrf1 availability, mixed proteasome factor and ER sterol target; algebraic functions, not additional states. |
| $\alpha_i, \beta_i$ | State-dependent establishment and loss coefficients. |
| $v_i^+, v_i^-, a_i^+, a_i^-$ | Unscaled node fluxes and time-scaled rates, with $a_i^\pm = v_i^\pm/\tau_i$. |
| $f_i, \mathbf{f}, \mathbf{J}$ | Coordinate drift, full drift vector and its full eleven-dimensional Jacobian. |
| $J_+, J_-$ | Stress-source and stress-relief fluxes. These are the two fluxes named in main-text Eq. (2). |
| $\mathbf{W}, \mathbf{B}, \mathbf{D}$ | Standard Wiener process vector, noise amplitude shape and diffusion shape; $\mathbf{D}$ equals B times its transpose. |
| $\mathbf{Q}, \varphi$ | Antisymmetric mobility component and scalar potential in the stated decomposition. |
| $\boldsymbol{\xi}, \mathbf{C}_0, \mathbf{\Sigma}, \mathbf{U}$ | Local deviation; covariance per unit noise; actual covariance $\mathbf{\Sigma} = \varepsilon \mathbf{C}_0$; precision $\mathbf{U} = \mathbf{C}_0^{-1}$. |
| $\varphi_\ell$ | Local quadratic potential, with an arbitrary additive constant. |
| $\mathcal{S}, \mathcal{A}$ | Symmetric dissipative and antisymmetric transverse matrices; calligraphic $\mathcal{S}$ differs from scalar p62 state S. |
| $\mathcal{I}_T$ | Finite-time weak-noise path action; calligraphic $\mathcal{I}$ differs from scalar INSIG1 state I. |
| $\mathbf{\Lambda}, \delta, m, \mathbf{r}_j, \mathbf{J}_j$ | Inverse diffusion weight $\mathbf{\Lambda} = \mathbf{D}^{-1}$, time increment $\delta = T/m$, number of path segments, midpoint drift residual and midpoint Jacobian in S11.3. |
| $s, t, t_0, T_i, x_i, x_i^\circ$ | Physical time, normalized time, reference time, process scale, physical functional readout and its reference range. No physical reference values were assigned in this realization. |
| $P_\dagger, o_+, \mathbf{r}$ | Specified proteasome target, maximal accessible Nrf2 output and residual of the reverse-adjoint uphill equation. |

Scalars are italic, vectors and matrices bold, and operators, differentials and transpose symbols upright. Subscripts identify nodes or coefficients. The calligraphic dissipative matrix and action are distinct from the scalar p62 and INSIG1 states.

### Main text and supplementary equation guide

| Main text | Mathematical content | Detailed derivation |
|---|---|---|
| Eq. (1) | Bounded balance and Hill response | S2; S10.2; reference catalogue S0 |
| Eq. (2) | Stress source and relief | S10.3, Eqs. (S34)-(S42) |
| Eq. (3) | Nuclear Nrf1 balance | S2 Eq. (S5); S10.2; S11.1 |
| Eqs. (4a)-(4b) | Organelle and proteasome maintenance | S2 Eqs. (S6)-(S7); bounds in S10.3 |
| Eqs. (5a)-(5c) | Nrf2 response, KEAP1 restraint and p62 balance | S2 Eqs. (S10)-(S12); S11.1 |
| Eqs. (6a)-(6b) | Membrane availability and INSIG1 turnover | S10.4, Eqs. (S43)-(S46); sign derivation S11.1 |
| Eq. (7) | A-type stochastic formulation | S6; S10.6-S10.8; numerical gradient S11.3 |
| Eq. (8) | Summary of Eqs. (1)-(2) | S2 (S2), (S4), (S8)-(S9); S10.2-S10.3 |

### S1 Biological organization of the network

Nrf1 denotes the CNC transcription factor encoded by NFE2L1; Nrf2 denotes NFE2L2. The Yin-Yang framework integrates Nrf1-associated basal redox homeostasis, organelle integrity and proteasome control with inducible Nrf2 defense [40] (Figure S1). The model resolves these functions into coupled maintenance, stress-response and sterol-handling processes.

Nuclear Nrf1 supports organelle maintenance [1], proteasome recovery [5,9,10] and functional KEAP1 restraint [3]. KEAP1 and proteasome function jointly regulate Nrf2 turnover [13]. p62 weakens KEAP1 restraint [17,19,20], while Nrf1 contributes separately to p62 production and clearance [18]. Nonzero basal and Nrf2-supported maintenance are retained [21].

The sterol branch links ER cholesterol sensing [6] to Nrf1-associated INSIG1 turnover and the SREBP1-supported export program [7]. The export contribution is defined at a fixed nutritional background. Nuclear Nrf1 input, membrane availability and interaction strength are independently adjustable. Sterol-associated stress and unresolved-stress amplification connect this branch to the redox network and are examined by parameter variation and deletion.

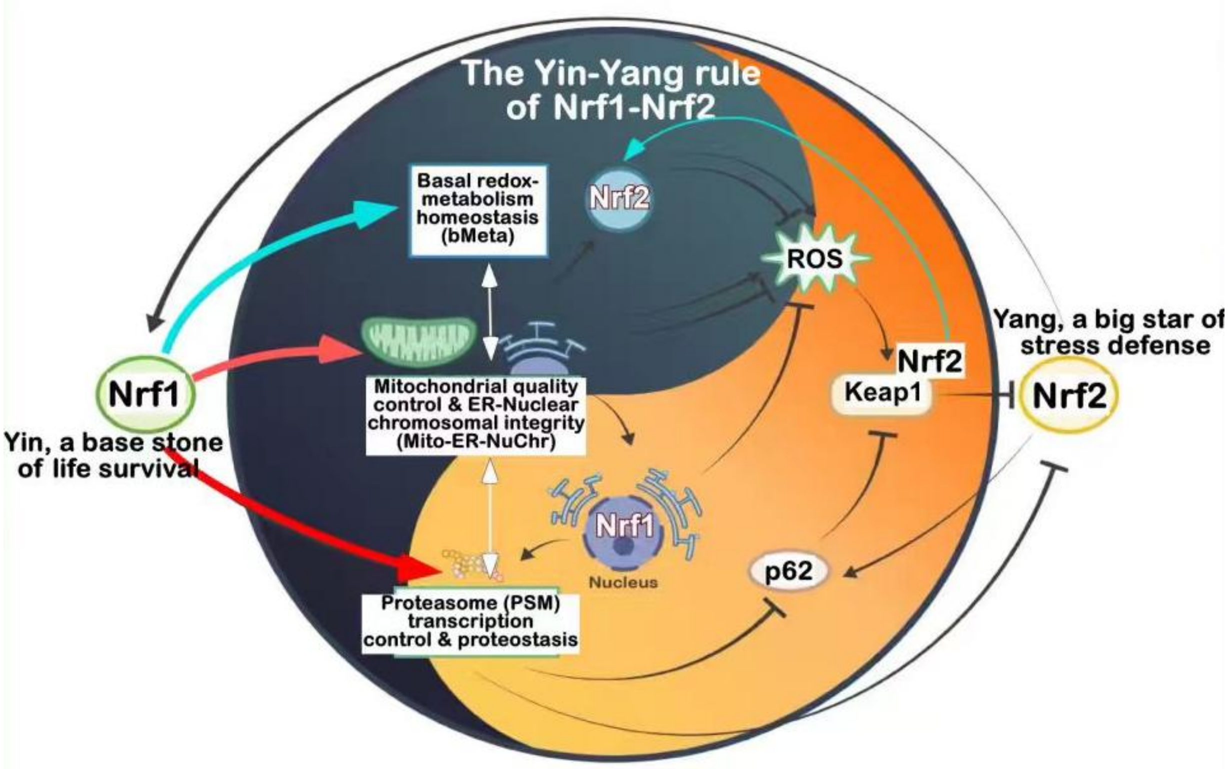


Figure S1. Yin-Yang organization of Nrf1 and Nrf2. The supplied schematic connects Nrf1-associated basal redox homeostasis, organelle integrity and proteasome control to Nrf2-mediated inducible defense, ROS, KEAP1 and p62. This conceptual framework is discussed in the review by Chen et al. [40].

### S2. State variables and complete equations

All calculations use the same coordinate order, with state meanings defined in S0:

$$\mathbf{z} = (N, M, P, R, Y, K, S, C, I, V, H)^{\mathrm{T}} \quad \text{(S1)}$$

Define the common Hill gate, Nrf2 output and three process-specific gates. The ER sterol gate $c_f$ is $h_C(C)$; its exponent, like the INSIG1 and burden exponents, is listed separately in S0.4.

$$h(x) = \frac{x^n}{K_h^n + x^n}, \quad o = q_2 h(Y), \quad h_j(x) = \frac{x^{n_j}}{K_j^{n_j} + x^{n_j}}, \quad j \in \{C, I, H\} \quad \text{(S2)}$$

Membrane Nrf1 availability $e$, proteasome dependency $p_u$ and the ER sterol target $c_t$ are algebraic functions:

$$e = \gamma_{\mathrm{e}}[\theta_e + (1 - \theta_e) h_C(C)], \quad p_u = (1 - f_u) + f_u P, \quad c_t = c_b + \kappa H \quad \text{(S3)}$$

The algebraic factor $e$ is membrane Nrf1 availability, controlled by $\gamma_e$; $I$ is INSIG1 abundance. The interaction increases INSIG1 loss. Setting $f_u = 0$ removes proteasome dependence from this channel, and $\kappa = 0$ removes burden-to-ER feedback. S10.4 derives the interpolation.

The production-loss formulation follows the endogenous-network strategy of representing regulated establishment and removal [15,16]. The available-range and occupied-range factors give the bounded functional-state balance derived in S10.2.

The positive and negative fluxes below precede division by $\tau_i$. Their difference gives the deterministic drift in main-text Eq. (1):

$$\tau_i \frac{\mathrm{d}z_i}{\mathrm{d}t} = v_i^+ - v_i^-, \qquad f_i = \frac{v_i^+ - v_i^-}{\tau_i} \tag{S4}$$

Nuclear Nrf1. Sterol-dependent retention attenuates establishment; current nuclear activity undergoes proportional loss. Both optional transcriptional feedback gains are zero at reference.

ER retention provides the biological motivation for the sterol-dependent input factor [6]; Eq. (S5) is its effective network realization.

$$v_N^+ = \frac{\gamma_{\mathrm{n}}[b_N + a_N h(N) + a_{2N} o]}{1 + \chi r_C h_C(C)}(1 - N), \quad v_N^- = d_N N \tag{S5}$$

Organelle maintenance. Basal and Nrf1/Nrf2-supported establishment oppose basal and stress-dependent loss.

Nrf1-dependent mitochondrial and redox homeostasis motivates this maintenance arm [1].

$$v_M^+ = [b_M + a_{NM} h(N) + a_{2M} o](1 - M), \quad v_M^- = [d_M + d_{RM} h(R)]M \tag{S6}$$

Proteasome function. Basal and both transcription-factor inputs promote establishment; stress increases functional loss.

Nrf1-mediated proteasome recovery and Nrf2-supported proteasome expression motivate the parallel inputs [5,21].

$$v_P^+ = [b_P + a_{NP} h(N) + a_{2P} o](1 - P), \quad v_P^- = [d_P + d_{RP} h(R)]P \tag{S7}$$

Stress burden. Five source terms act on the available burden range. Three relief terms act on the existing burden. Their difference is main-text Eq. (2).

$$v_R^+ = [b_R + u(t) + a_{MR}(1 - M) + a_R h(R) + \chi a_H h_H(H)](1 - R) \tag{S8}$$

$$v_R^- = [d_R + k_{PR} P + k_{2R} o]R \tag{S9}$$

Nrf2 response. Basal and stress-induced establishment oppose basal turnover and joint KEAP1-proteasome-dependent turnover.

KEAP1-dependent regulation and proteasomal turnover are supported by quantitative KEAP1-Nrf2 modeling [13,14].

$$v_Y^+ = \gamma_2[b_Y + a_{RY} h(R)](1 - Y), \quad v_Y^- = [d_Y + k_{KP} K P]Y \tag{S10}$$

KEAP1 restraint. Basal and Nrf1-supported establishment oppose basal loss and p62-associated loss of effective restraint.

Nrf1-loss-associated changes in KEAP1 protein and the p62-KEAP1 interaction motivate the effective restraint balance [3,17].

$$v_K^+ = [b_K + a_{NK} h(N)](1 - K), \quad v_K^- = [d_K + a_{SK} h(S)]K \tag{S11}$$

p62-associated abundance. Production supported by Nrf1 and Nrf2 is distinct from Nrf1-supported clearance. Their net Nrf1 dependence is derived in Eq. (S42).

Nrf1-dependent p62 induction and aggrephagy motivate distinct production and clearance contributions [18].

$$v_S^+ = [b_S + a_{NS}^+ h(N) + a_{2S} o](1 - S), \quad v_S^- = [d_S + a_{NS} h(N)]S \tag{S12}$$

ER sterol signal. Complementary establishment and loss coefficients give relaxation toward the burden-dependent target:

$$v_C^+ = c_t(1 - C), \qquad v_C^- = (1 - c_t)C, \qquad \tau_C \frac{\mathrm{d}C}{\mathrm{d}t} = c_t - C \tag{S13}$$

INSIG1 and export. Interaction-linked turnover requires membrane Nrf1 availability and the proteasome-dependency factor. Lower INSIG1 releases export-program establishment.

Nrf1-INSIG1 interaction, proteasomal destabilization of INSIG1 and SREBP1-supported secretion motivate this serial branch [7].

$$v_I^+ = b_I(1 - I), \qquad v_I^- = [d_I + \eta k_I e p_u]I \tag{S14}$$

$$v_V^+ = [b_V + a_V(1 - h_I(I))](1 - V), \quad v_V^- = d_V V \tag{S15}$$

Sterol burden. Constitutive and pulsed inputs oppose basal, export-associated and optional Nrf2-supported clearance.

$$v_H^+ = [j_H + u_H(t)](1 - H), \qquad v_H^- = [d_H + k_V V + k_{2H} o]H \tag{S16}$$

Nonnegative rates, positive times and a target within the unit interval give inward-pointing boundary drift. S10.2 proves deterministic invariance; trajectories were checked without clipping.

Figure 2D reports each source and relief contribution divided by $\tau_R$. The proteasome-associated term represents clearance of the damaged-protein component of the coupled burden.

The eleven-state equations combine published biological interactions with a common bounded response architecture. Literature specifies the supported directions of organelle maintenance, proteasome control, KEAP1-mediated turnover and membrane sterol handling [1,3,5,6,7,13,17,18,21]. S10 derives the normalization, invariance, response slopes and algebraic closures of this particular network. S0 specifies its numerical realization, while S12 uses operating states, turnover and perturbations to organize parameter construction and uncertainty analysis. The predictive content is obtained by solving the assembled network and varying these assignments.

### S3. Parameterization and relationship to the legacy seven-variable model

Substitution of S0 values into S2 reproduces the archived eleven-state implementation. The companion parameter ledger links each mathematical symbol to its code field or fixed coefficient.

Nrf1 autoregulation and Nrf2-to-Nrf1 transcriptional support are available as optional terms but are disabled in the reference network. These optional terms permit separate evaluation of additional transcriptional feedbacks. When Nrf1 autoregulation is enabled, the code bypasses the reduced equilibrium solver and uses the full eleven-dimensional solver.

The updated seven-function control sets the sterol-feedback switch to zero, leaving the four sterol states downstream. The original seven-variable model also differs in basal production, Nrf2 support and p62 clearance. Separate root inventories distinguish these changes from addition of sterol feedback.

### S4. Fixed points, stability and folds

With $a_N$ equal to zero, $C$, $N$, $M$, $P$, $S$, $K$, $I$ and $V$ can be solved algebraically at equilibrium for supplied $R$, $Y$ and H. The remaining three equilibrium equations are solved numerically and mapped back to eleven dimensions. This exact elimination is used for equilibrium searches; trajectories are integrated in the full eleven-dimensional state space.

The reference reduced search used 1024 scrambled Sobol starts plus five structured initial states. SciPy hybr used xtol = $10^{-10}$ and a maximum of 400 function evaluations. The full eleven-dimensional cross-check used 256 Sobol starts plus seven structured starts and at most 700 evaluations. Roots were accepted only if the full drift infinity norm was below $10^{-8}$ and coordinates were in the unit interval to a tolerance of $10^{-7}$. Roots within Euclidean distance $10^{-5}$ were deduplicated. Reduced searches were repeated with seeds 17, 71 and 307. The accepted multistart root inventories are compared across seeds and between the reduced and full-state searches.

The Jacobian used complex-step differentiation with step $10^{-28}$ and was compared against central differences with step $10^{-6}$ at 100 random states. A stable root required all eigenvalue real parts below $-10^{-7}$. Positive real parts above $10^{-7}$ were counted as unstable directions. Opposite perturbations of size $10^{-5}$ along the reference saddle's unique unstable eigenvector were integrated for 180 model time units. Distances to the terminal attractors are retained in data/summary.json.

Adaptive trajectories used DOP853, relative tolerance $2 \times 10^{-9}$, absolute tolerance $2 \times 10^{-11}$ and maximum step one. Failed solves raise errors and are not classified as biological outcomes.

The membrane map sampled 21 $\eta$ values from zero to one and 21 sustained sterol inputs from 0.02 to 0.42. Each grid cell used 16 Sobol starts plus roots from the previous cell. Fold candidates jointly solved the three reduced equilibrium equations and a zero determinant of their equilibrium Jacobian. Full-state eigenvectors were then used for nondegeneracy checks:

$$\mathbf{J}\mathbf{v} = 0, \quad \mathbf{w}^{\mathrm{T}}\mathbf{J} = 0, \quad \mathbf{w}^{\mathrm{T}}\mathbf{v} = 1 \tag{S17}$$

$$\mathbf{w}^{\mathrm{T}} \frac{\partial \mathbf{f}}{\partial j_H} \neq 0, \quad \mathbf{w}^{\mathrm{T}}\mathbf{f}_{zz}[\mathbf{v}, \mathbf{v}] \neq 0 \tag{S18}$$

The second directional derivative used central differences with step $10^{-4}$. The fold input values at $\eta$ = 0, 0.25, 0.50, 0.75 and 1 were 0.081214, 0.168474, 0.219769, 0.236410 and 0.242101, respectively. The residual, zero-eigenvalue and nondegeneracy checks characterize these five local folds.

In the landscape interpretation, a stable equilibrium is the center of a locally restoring state. A perturbation changes the state reached at washout or changes the drift itself. These are different operations: the former probes the reference attraction geometry, whereas the latter can move equilibria and folds. The full-state calculations in S4-S5 distinguish them.

### S5. Perturbation, robustness and recovery protocols

The 128-scenario ensemble sampled a nine-dimensional Sobol sequence with seed 2201. The ranges were: $a_R$, 0.8-2; $a_H$, 0.1-0.6; $k_I$, 1-4; $k_V$, 0.4-1.2; $\kappa$, 0-0.4; $a_{2P}$, 0-0.6; $a_{2M}$, 0-0.3; and $b_P$ and $b_M$, each 0.02-0.20. The intact and $\eta$ = 0 networks were solved separately. Comparisons used the lowest-$R$ stable state in each system. The stored field low_branch_lost is an operational category: intact minimum $R$ < 0.25 and perturbed minimum $R$ > 0.40. This category records a change in the minimum stable burden under the stated thresholds. Hill sensitivity used $n$ = 2, 3 or 4 and $K_h$ = 0.35, 0.42 or 0.50.

The compensatory-bypass scan set $\gamma_{\mathrm{n}}$ and $\gamma_{\mathrm{e}}$ to zero and $\gamma_2$ to four. Joint $a_{2M}$ and $a_{2P}$ values ranged from zero to four in eleven steps, and alternative sterol-clearance gain $k_{2H}$ ranged from zero to one in eleven steps. Of these 121 combinations, 102 contained a stable state with $R$ < 0.25. The reported count summarizes stable-state existence across the specified bypass-strength combinations.

The joint-input map set $\gamma_{\mathrm{n}}$ = $\gamma_{\mathrm{e}}$ to 21 levels between zero and one and $\gamma_2$ to 21 levels between zero and four. All 441 trajectories began at the reference low-load attractor and were integrated for 250 units using fourth-order Runge-Kutta at steps 0.05 and 0.025. Eight endpoints were checked independently with DOP853. The maximum step-halving discrepancy was $3.91 \times 10^{-14}$, the maximum final drift residual was $1.54 \times 10^{-14}$, and the maximum DOP853 discrepancy was $2.93 \times 10^{-10}$. No extension beyond 250 units was needed.

External pulses were applied to either $u$ or $u_H$, never both simultaneously. Duration was 0.5, 1, 2, 4, 8 or 16. The directly evaluated grid used 25 amplitudes from zero to 0.5 for stress input and from zero to two for sterol input, giving 300 protocols. Each was followed by 250 unforced time units. Endpoint assignment used Euclidean distance in the full eleven-dimensional state space to the two reference attractors. Step halving produced zero classification changes; the maximum endpoint distance to its assigned attractor was $4.58 \times 10^{-12}$. For display in Figure 5C, saved trajectories were linearly interpolated onto a common time grid.

Threshold searches independently bracketed a switching protocol between zero and an amplitude of one for stress or three for sterol, confirmed a high-load endpoint at the upper bracket, then performed 16 bisections. The retained upper bounds are:

| Duration | Stress-input threshold | Sterol-input threshold |
|---|---|---|
| 0.5 | 0.329910 | 1.773468 |
| 1 | 0.181335 | 0.884995 |
| 2 | 0.107513 | 0.448471 |
| 4 | 0.071365 | 0.240143 |
| 8 | 0.054367 | 0.146210 |
| 16 | 0.047058 | 0.108490 |

Rescue simulations first imposed $\eta$ = 0 for 250 units from the reference low state. Five protocols then ran for 200 units: no change; $\eta$ restored to one; $b_I$ reduced from 0.20 to 0.04; $\gamma_2$ increased from one to four; or $\eta$ restoration combined with $\gamma_2$ enhancement. Results are compared directly in normalized sterol burden $H$ and redox stress burden $R$ at the specified intervention strengths.

### S6. A-type interpretation and local stochastic construction

The formal Ao representation separates gradient and rotational contributions [22,23,25,26]. The attractor-local construction follows the linear fixed-point analysis of Kwon, Ao and Thouless [24]:

The random state is $\mathbf{Z}$, $\mathbf{W}$ is a vector of independent Wiener processes, and $\mathbf{B}\mathbf{B}^{\mathrm{T}} = \mathbf{D}$. Noise increments have covariance $2\varepsilon\mathbf{D}\,\mathrm{d}t$. The asterisk denotes A-type integration:

$$\mathrm{d}\mathbf{Z} = \mathbf{f}(\mathbf{Z})\mathrm{d}t + \sqrt{2\varepsilon}\mathbf{B}(\mathbf{Z}) * \mathrm{d}\mathbf{W}, \qquad BB^{\mathrm{T}} = \mathbf{D} \tag{S19}$$

$$\mathbf{f} = -(\mathbf{D} + \mathbf{Q})\nabla\varphi, \quad \mathbf{D}^{\mathrm{T}} = \mathbf{D}, \quad \mathbf{Q}^{\mathrm{T}} = -\mathbf{Q} \tag{S20}$$

The potential is $\varphi$, and $\mathbf{Q}$ is antisymmetric. With density $\rho$ and $\partial_i = \partial/\partial z_i$, repeated coordinate indices are summed:

$$\partial_t\rho = \partial_i\{(\mathbf{D}_{ij} + \mathbf{Q}_{ij})[\partial_j\varphi\,\rho + \varepsilon\partial_j\rho]\} \tag{S21}$$

For regular coefficients, normalizable density and compatible boundaries, $\rho \propto \exp(-\varphi/\varepsilon)$ is stationary. The equivalent Itô drift is:

$$\mathrm{d}Z_i = [f_i + \varepsilon\partial_j(\mathbf{D}_{ij} + \mathbf{Q}_{ij})]\mathrm{d}t + \sqrt{2\varepsilon}\mathbf{B}_{i\alpha}\mathrm{d}W_\alpha, \qquad BB^{\mathrm{T}} = \mathbf{D} \tag{S22}$$

The divergence term specifies the drift correction in the Itô representation. The construction used here evaluates the drift Jacobian and diffusion at each stable point, giving a constant-coefficient local stochastic system.

Writing the displacement from a stable point as $\boldsymbol{\xi}$, the resulting local process is:

$$\mathrm{d}\boldsymbol{\xi} = \mathbf{J}\boldsymbol{\xi}\mathrm{d}t + \sqrt{2\varepsilon}\mathbf{B}\mathrm{d}\mathbf{W} \tag{S23}$$

Its covariance coefficient and local potential are obtained from:

$$\mathbf{J}\mathbf{C}_0 + \mathbf{C}_0\mathbf{J}^{\mathrm{T}} + 2\mathbf{D} = 0, \qquad \boldsymbol{\Sigma} = \varepsilon\mathbf{C}_0 \tag{S24}$$

$$\mathbf{U} = \mathbf{C}_0^{-1}, \qquad \mathbf{Q} = -\mathbf{J}\mathbf{C}_0 - \mathbf{D}, \qquad \varphi_\ell(\boldsymbol{\xi}) = \frac{1}{2}\boldsymbol{\xi}^{\mathrm{T}}\mathbf{U}\boldsymbol{\xi} \tag{S25}$$

The Lyapunov equation guarantees antisymmetry of $\mathbf{Q}$ and the identity $\mathbf{J} = -(\mathbf{D} + \mathbf{Q})\mathbf{U}$. The symmetric and antisymmetric parts of the inverse mobility are:

$$\mathbf{G} = (\mathbf{D} + \mathbf{Q})^{-1}, \qquad \mathcal{S} = \frac{\mathbf{G}+\mathbf{G}^{\mathrm{T}}}{2}, \qquad \mathcal{A} = \frac{\mathbf{G}-\mathbf{G}^{\mathrm{T}}}{2} \tag{S26}$$

The Lyapunov identity makes the transverse component antisymmetric. Constant local coefficients have no convention-dependent drift correction. The quadratic potential decreases along local deterministic relaxation; S10.6-S10.8 derive the density conversion, covariance and action relationships.

Two diffusion conventions were evaluated, using time-scaled fluxes $a_i = v_i/\tau_i$. The symbol $\mathbf{1}_{11}$ denotes the eleven-dimensional identity matrix, distinct from the scalar INSIG1 state I.

$$\mathbf{D}_0 = \mathbf{1}_{11}, \qquad \mathbf{D}_v = \frac{1}{2}\mathrm{diag}(a_i^+ + a_i^-) \tag{S27}$$

The effective-flux convention scales local fluctuation intensity with the sum of effective production and loss rates. Isotropic and effective-flux diffusion are evaluated for local deviations around each attractor, with $\varepsilon$ setting the common noise scale. This comparison tests the sensitivity of local fluctuation structure to the chosen diffusion model.

| State | Diffusion | Trace of covariance divided by epsilon |
|---|---|---|
| Low load | Isotropic | 14.068863 |
| High load | Isotropic | 12.009374 |
| Low load | Effective flux | 1.702073 |
| High load | Effective flux | 2.544608 |

Figure S3A uses the stress/proteasome covariance submatrix at $\varepsilon = 10^{-4}$ to draw a 68.3% contour of the local Gaussian approximation. The reversed covariance-trace ranking shows dependence on diffusion choice. S11.4 derives the contour radius.

The illustration scale is selected from the archived covariance per unit noise, $C_0$, by requiring $3\sqrt{\varepsilon(C_0)_{ii}} \leq min(z_i^*, 1 - z_i^*)$ for every coordinate at both attractors under both diffusion conventions. The most restrictive ceiling is $1.6538 \times 10^{-4}$; the common choice $\varepsilon = 10^{-4}$ satisfies this coordinatewise screening rule. It sets the displayed width while leaving the covariance-per-noise and potential-curvature calculations unchanged. Joint confinement and nonlinear fluctuations depend additionally on the surrounding drift and boundary dynamics.

Figure S8 maps the deterministic, local stochastic and transition-action calculations to their derivations.

#### *S6.1 From local restoring dynamics to landscape geometry*

The landscape construction assigns a scalar cost to deviations around a stable network state [24-26]. The deterministic Jacobian describes restoring and coupled motion; diffusion specifies how fluctuations enter. Their Lyapunov balance determines covariance, whose inverse supplies the curvature of the local quadratic potential in Eq. (S25). Noise intensity sets the probability scale through the ratio of potential to noise, rather than changing that quadratic curvature. The potential is dimensionless and is interpreted as a stochastic state cost, not a molecular energy measured in joules.

Figure S9 displays the stress/proteasome marginal geometry obtained from the corresponding covariance submatrix, following S11.4. The inverse of this marginal covariance defines the plotted quadratic cost; it is not obtained by selecting a block of the full precision matrix. Each surface is centered at its own attractor with zero cost there, and all panels use the same coordinate window and vertical scale. The change in geometry between isotropic and effective-flux diffusion shows why restoring drift alone does not determine fluctuation width. Relative well populations additionally depend on normalization and the connections between states.

### S7. Eleven-dimensional candidate transition action

The finite-time variational formulation follows the minimum-action approach of E, Ren and Vanden-Eijnden [41]. The geometric reformulation [27] is related theoretical context rather than the discretization executed here.

For isotropic diffusion and noise amplitude $\sqrt{2\varepsilon}$, the finite-time action is:

$$\mathcal{I}_T[\mathbf{z}] = \frac{1}{4}\int_0^{\mathrm{T}}[\dot{\mathbf{z}} - \mathbf{f}(\mathbf{z})]^{\mathrm{T}}\mathbf{D}^{-1}[\dot{\mathbf{z}} - \mathbf{f}(\mathbf{z})]\mathrm{d}t \quad \text{(S28)}$$

Fixed-endpoint paths connect each stable point to the saddle. A regular order-epsilon drift correction does not change the leading weak-noise action. S10.8 derives its relation to a compatible potential difference.

The eleven-coordinate endpoints were fixed. Interior path nodes were constrained to [0,1]. Midpoint quadrature and an analytic discrete gradient were used; the gradient was checked against central differences at 20 randomly selected coordinates. L-BFGS-B used ftol = $10^{-13}$, gtol = $10^{-7}$, maxcor = 20 and maxls = 40. Path stages used $T$ = 12, 24 and 48 with 60, 120 and 240 segments, followed by a curved alternative initial path, 480 segments at $T$ = 48, and 960 segments at $T$ = 96. Stage iteration limits and solver flags remain in the source scripts and output tables.

| Discretization | Low load to saddle | High load to saddle |
|---|---|---|
| $T$ = 12, 60 segments | 0.001581648 | 0.025111027 |
| $T$ = 24, 120 segments | 0.001441315 | 0.024820056 |
| $T$ = 48, 240 segments | 0.001437079 | 0.024811739 |
| $T$ = 48, curved initial path | 0.001437093 | 0.024811750 |
| $T$ = 48, 480 segments | 0.001436693 | 0.024767138 |
| $T$ = 96, 960 segments | 0.001436689 | 0.024767131 |
| $T$ = 48 path, eight-point Gauss check | 0.001436640 | 0.024765305 |

The table reports finite-time optimized actions. Horizon extension, mesh refinement, alternative initialization and independent quadrature assess numerical consistency. The executed solver uses discrete finite-time minimization; the geometric method [27] supplies related theoretical context.

#### *S7.1 From local curvature to direction-dependent transition cost*

Local curvature describes small fluctuations within a state; leaving that state requires a path through the nonlinear network. The action in Eq. (S28) accumulates the diffusion-weighted discrepancy between path velocity and deterministic drift [41]. Figure S10 shows the stored eleven-dimensional paths through their stress/proteasome projection and accumulated costs. At horizon 96 with 960 segments, the candidate costs to the reference saddle are 0.001436689 from low stress and 0.024767131 from persistent stress. These results distinguish the tested directions of departure under isotropic diffusion.

For a compatible potential representation, the square-completion identity in S10.8 relates action to potential difference plus a nonnegative residual. Equality requires the stated uphill connection. Thus path refinement and horizon extension connect the numerical calculation to the landscape interpretation, while the finite-time candidates retain their defined endpoints and horizon. Their values do not by themselves specify transition-time prefactors or stationary population ratios.

### S8. Expression analysis and source images

The laboratory HepG2 matrix contained 16,660 gene rows and 15 FPKM samples. Symbols were trimmed and converted to uppercase. Duplicate symbols were averaged on the original expression scale before log2(expression + 1). Each gene was standardized within its dataset using the sample standard deviation (ddof = 1). Zero-variance genes were excluded. A module score was the mean of available gene $z$-scores. Four nonoverlapping sets were specified:

| Module | Genes |
|---|---|
| Proteasome | PSMA1-7; PSMB1-7; PSMC1-6; PSMD1, PSMD2, PSMD3, PSMD4, PSMD6, PSMD7, PSMD8, PSMD9, PSMD11, PSMD12, PSMD13, PSMD14 |
| Nrf2-responsive output | NQO1, GCLC, GCLM, TXNRD1, SLC7A11 |
| Processing-associated | DDI2, NGLY1, VCP, UFD1, NPLOC4, SYVN1 |

| Module | Genes |
| --- | --- |
| ISR | DDIT3, TRIB3, ASNS, PPP1R15A, CHAC1 |

All 32 proteasome genes, five Nrf2-output genes and five ISR genes were retained in both datasets. Five of the six processing-associated genes were retained in HepG2 (UFD1 was not matched), and all six were retained in liver. All scores and statistical comparisons reported here use these four predefined, nonoverlapping modules. Gene coverage is saved in data/expression_module_coverage.csv, with source provenance and gene-level tables alongside it. The scores summarize transcriptional programs and are compared with the functional directions of the network.

Four contrasts were specified: Nrf1α deficiency versus control; additional siNrf2 versus Nrf1α deficiency; Nrf2 DeltaTA versus control; and caNrf2 DeltaN versus control. For each contrast and module, all 20 allocations of six samples into groups of three were enumerated. The two-sided statistic was the absolute difference in means. The minimum possible exact $P$ value was 0.10; BH adjustment covered all 16 tests. No comparison reached $q < 0.05$. Connecting segments in Figure 1C are effect-size guides, not uncertainty intervals.

The liver .xls source is actually tab-delimited text and contains 30,156 gene rows across seven sample-level average-expression profiles. The unit analyzed here is the supplied sample-level hepatocyte mean, and module scores are calculated across the seven profiles. A3 is the single 8-week normal mean; A5T/A6T are 8-week knockout means; A8/A9 are 9-month normal means; and A13T/A14T are 9-month HCC-related means. The two tumor-related profiles are presented individually to retain their sample-specific expression patterns.

The four H&E fields in Figure 6C retain their original pixels and embedded scale bars. Only panel placement and external labels are adjusted. Histology and sample-level expression are separate, nonpaired observations.

### S9. Supplementary figures and reproduction record

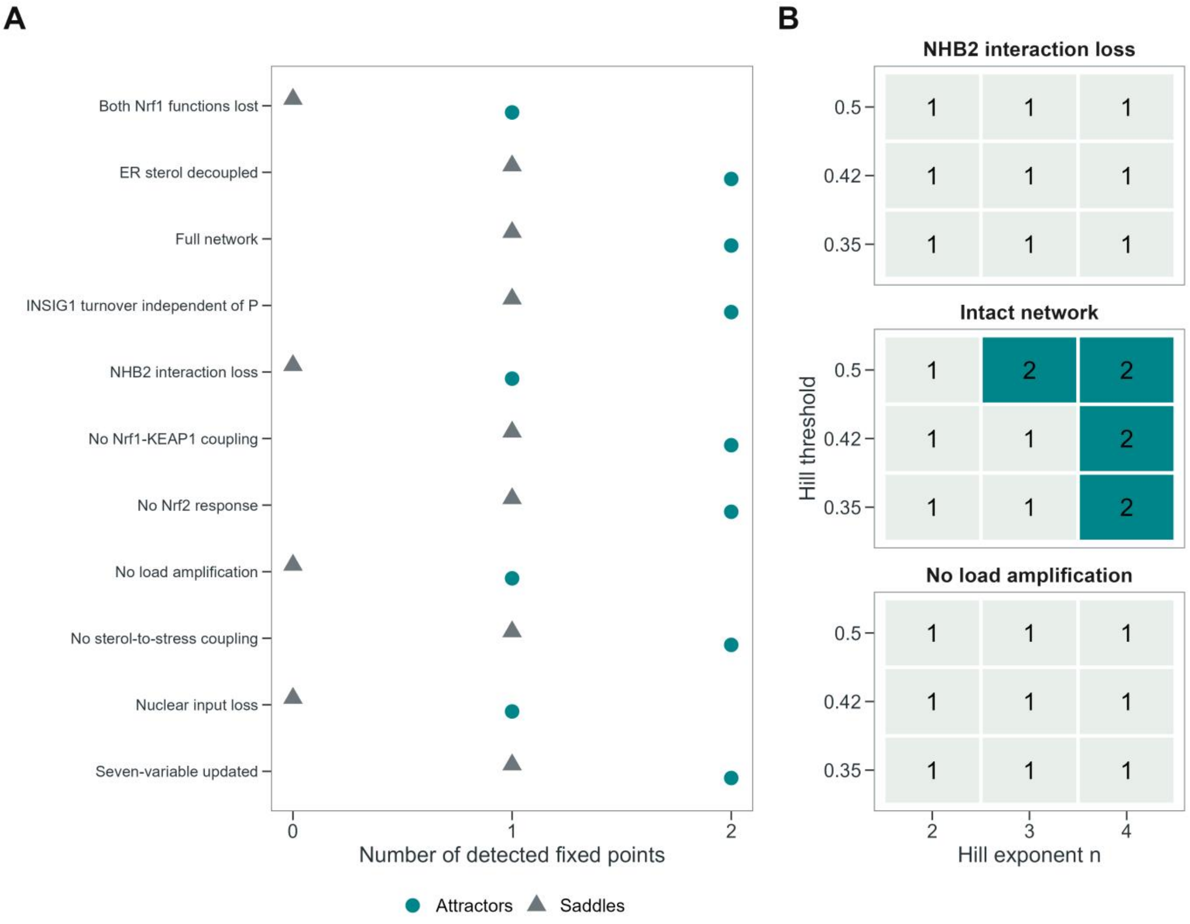


Figure S2. Topological and parameter sensitivity. A, Detected stable points and index-one saddles under the indicated network changes. B, Stable-point counts across nine Hill parameter settings. Searches use the residual, domain and eigenvalue criteria in S4; the source tables retain coordinates and eigenvalues.

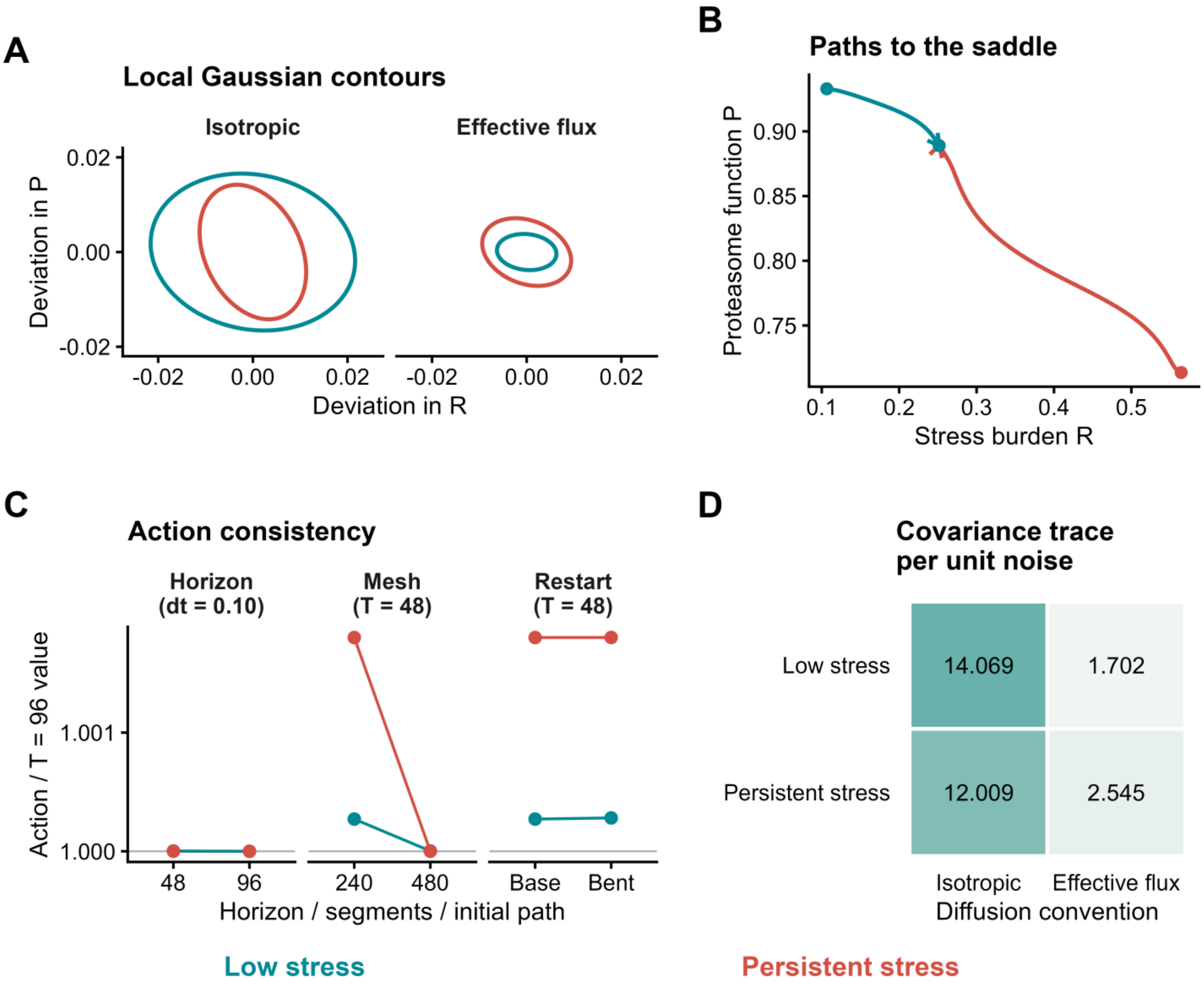


Figure S3. Local fluctuations and separate action-consistency checks. A, 68.3% contours of the local Gaussian stress/proteasome marginals at $\varepsilon = 10^{-4}$, selected by the S6 boundary screening rule. B, Archived eleven-dimensional attractor-to-saddle paths projected onto stress burden and proteasome function, at horizon 96 with 960 segments. C, Candidate actions divided by the corresponding direction-specific value at horizon 96. Horizon extension compares 48/480 and 96/960 time/segment pairs at time step 0.10; mesh refinement compares 240 and 480 segments at horizon 48; restart compares the base and bent initial paths at horizon 48 with 240 segments. All optimization stages remain listed in S7. D, Full covariance trace per unit noise under each diffusion convention.

The companion source tables retain the joint-input, pulse, flux, equilibrium and action calculations. Quantitative figures are redrawn from these tables; histology remains raster image data. S12 lists the additional computational designs and their reconstruction files.

### S10. Derivation, mathematical conditions and scope of the network formulation

#### *S10.1. Biological closure and the meaning of the state space*

The eleven coordinates separate defense activation from maintenance, turnover and stress-source control. Seven retain the redox/proteostasis core; four resolve ER sterol signaling, INSIG1, export and sterol burden. Individual coordinates and full-state dynamics describe recovery without an averaged capacity index.

The closure retains the feedbacks and time scales relevant to the chosen cellular and nutritional context. Other processes enter effective rates or the specified noise. A consequential omitted slow variable would require an additional state or a memory-dependent description [42].

#### *S10.2. Deriving bounded production-loss dynamics*

To derive main-text Eq. (1), first define a functional activity $x_i$ relative to a fixed reference range $x_i^\circ$ and a physical time $s$ relative to a reference time $t_0$. A characteristic process time $T_i$ defines the dimensionless relative time $\tau_i$:

$$z_i = x_i/x_i^\circ, \quad t = s/t_0, \quad \tau_i = T_i/t_0, \quad T_i \frac{\mathrm{d}z_i}{\mathrm{d}s} = \tau_i \frac{\mathrm{d}z_i}{\mathrm{d}t} \tag{S29}$$

The reference ranges define functional coordinates and the reference clock defines model time. Establishment acts on the available range, while loss acts on the occupied range:

$$\tau_i \dot{z}_i = \alpha_i(\mathbf{z})(1 - z_i) - \beta_i(\mathbf{z}) z_i \tag{S30}$$

$$v_i^+ = \alpha_i(\mathbf{z})(1 - z_i), \quad v_i^- = \beta_i(\mathbf{z}) z_i, \quad a_i^\pm = v_i^\pm/\tau_i, \quad f_i = a_i^+ - a_i^- \tag{S31}$$

Hill response functions provide a compact representation of saturable regulatory input [43]. Here the exponent specifies response steepness; the gate is differentiated and scanned within the stated rate-law family.

These definitions give the eleven balances in S2 and the time-scaled rates used for deterministic integration and effective-event noise. The common response gate follows from normalized inactive and active weights:

$$w_0 = 1, \qquad w_1 = (x/K_h)^n, \qquad h(x) = \frac{w_1}{w_0 + w_1} = \frac{x^n}{K_h^n + x^n} \tag{S32}$$

Here $n$ describes effective nonlinearity. At unit input, $h(1) = 1/(1 + K_h^n)$. Additive inputs describe parallel contributions; products describe joint dependencies.

At $z_i = 0$, drift is nonnegative; at $z_i = 1$, it is nonpositive. Locally Lipschitz rates and $\tau_i > 0$ therefore make the unit hypercube forward invariant. The domain requires $c_b \geq 0$, $\kappa \geq 0$, $c_b + \kappa \leq 1$, and $0 \leq f_u, \theta_e \leq 1$. Positive thresholds and the evaluated integer exponents ensure regularity; nonnegative pulses preserve the boundary signs.

When the two rates are frozen, the single-node solution is:

$$z_i(t) = z_i^* + [z_i(0) - z_i^*]\exp[-(\alpha_i + \beta_i)t/\tau_i], \qquad z_i^* = \frac{\alpha_i}{\alpha_i + \beta_i} \tag{S33}$$

The frozen rate ratio determines the operating level; the rate sum divided by the node time determines relaxation. In the network, inputs vary together and the full Jacobian determines local modes. Positive time changes preserve equilibrium positions; individual changes can alter stability, whereas a common change rescales the clock.

#### *S10.3. Protective output, persistent stress and an analytical compensation bound*

Main-text Eq. (2) is the $R$ component of Eq. (1), not a separate model. Identify its establishment coefficient by adding the five source contributions, and its loss coefficient by adding the three relief contributions. Multiplication by 1-$R$ and $R$, respectively, gives:

$$J_+ = [b_R + u(t) + a_{MR}(1 - M) + a_R h(R) + \chi a_H h_H(H)](1 - R) \tag{S34}$$

$$J_- = [d_R + k_{PR}P + k_{2R}o]R, \quad f_R = (J_+ - J_-)/\tau_R \tag{S35}$$

At a steady state the two fluxes are equal; their equality specifies a balance, while the value of $R$ specifies the remaining burden. Thus a large protective flux can coexist with a large source at a persistent-stress equilibrium.

Writing the burden equation as source minus relief distinguishes direct cytoprotection from whole-network recovery. For $o = q_2 h(Y)$, its direct partial effect is:

$$\frac{\partial f_R}{\partial o} = -\frac{k_{2R}R}{\tau_R} \leq 0 \tag{S36}$$

This partial derivative fixes other coordinates. Along a differentiable equilibrium branch, the complete response to a parameter follows by differentiating the zero-drift condition:

$$\frac{\mathrm{d}\mathbf{z}^*}{\mathrm{d}\mu} = -\mathbf{J}^{-1}\mathbf{f}_\mu \tag{S37}$$

The inverse Jacobian transmits the perturbation through all feedbacks. This identity requires a nonsingular Jacobian and is not evaluated at a fold.

Stress-dependent impairment of maintenance and proteasome function forms reinforcing double-negative loops. Stress-induced Nrf2 relief forms a buffering loop. Their gains and response ranges determine whether the full network admits alternative stable states.

The model also admits a sufficient condition for failure to restore a prespecified functional target without setting Nrf2 compensation to zero. With $\gamma_{\mathrm{n}}$=0, any equilibrium has $N^* = 0$. Therefore:

$$P^* = \frac{b_P + a_{2P} o^*}{b_P + a_{2P} o^* + d_P + d_{RP} h(R^*)} \tag{S38}$$

Since $0 \le Y \le 1$ and $h(R^*) \ge 0$, define $o_+ = q_2/(1 + K_h^n)$ and obtain:

$$P^* \le \frac{b_P + a_{2P} o_+}{b_P + a_{2P} o_+ + d_P}, \quad o_+ = \frac{q_2}{1 + K_h^n} \tag{S39}$$

Consequently, no increase in the Nrf2 production input $\gamma_2$ alone can produce a steady $P$ at or above a prespecified $P_\dagger$ in (0,1) if:

$$b_P + a_{2P} o_+ < \frac{d_P P_\dagger}{1 - P_\dagger} \tag{S40}$$

This condition permits nonzero basal synthesis and Nrf2-dependent maintenance. Its violation is necessary, but not sufficient, for reaching the target: stress-dependent loss and other constraints remain. An analogous bound for organelle maintenance is:

$$M^* \le \frac{b_M + a_{2M} o_+}{b_M + a_{2M} o_+ + d_M} \tag{S41}$$

At the reference settings, the proteasome and maintenance bounds are 0.710 and 0.496, compared with low-stress levels 0.933 and 0.911. Increasing Nrf2 production alone cannot overcome these bounds. Raising downstream support or basal establishment changes the bounds; the bypass experiment in S12.4 tests this possibility.

Separate p62 production and clearance also matter. The instantaneous derivative of the $S$ equation with respect to the Nrf1 Hill input is:

$$\frac{\partial f_S}{\partial h(N)} = \frac{a_{NS}^{+} - (a_{NS}^{+} + a_{NS}) S}{\tau_S} \tag{S42}$$

Nrf1-supported production and clearance therefore give a state-dependent net effect on p62 abundance [18].

#### *S10.4. Conditions on the sterol extension*

The algebraic membrane closure in main-text Section 2.1 begins with a saturating sterol gate $c_f$. Interpolate between a basal membrane availability $e_0$ and a sterol-enhanced availability $e_1$. With $e_0$=0.3 $\gamma_{\mathrm{e}}$ and $e_1$=$\gamma_{\mathrm{e}}$, the closure is:

$$c_f = h_C(C), \quad e = e_0(1 - c_f) + e_1 c_f, \quad e_0 = \theta_e \gamma_e, \quad e_1 = \gamma_{\mathrm{e}} \tag{S43}$$

The interpolation is nonnegative and bounded by $\gamma_e$. It describes fast membrane availability relative to the retained state dynamics. Nuclear activity retains its own equation and input.

Substitute the membrane availability and proteasome factor into the establishment-loss balance for INSIG1. The complete turnover coefficient is:

$$p_u = (1 - f_u) + f_u P, \quad \beta_I = d_I + \eta k_I e p_u, \quad \tau_I \dot{I} = b_I(1 - I) - \beta_I I \tag{S44}$$

Setting INSIG1 establishment equal to turnover gives its frozen-input stationary level:

$$I^* = \frac{b_I}{b_I + d_I + \eta k_I e p_u} \tag{S45}$$

INSIG1 restraint on the export-supporting program is represented by the complementary gate 1-$I^4/(K_I{}^4+I^4)$. The resulting program contributes $k_V\ V$ to the sterol-clearance coefficient. Combining synthesis and clearance gives:

$$\tau_H \dot{H} = [j_H + u_H(t)](1 - H) - [d_H + k_V V + k_{2H} o]H \tag{S46}$$

The interaction parameter $\eta$, membrane input $\gamma_e$ and nuclear input $\gamma_n$ specify distinct perturbations. The proteasomal contribution to INSIG1 turnover is supported by [7]; $f_u$ varies its dependence, with $f_u = 1$ as the reference setting.

The export term describes the SREBP1-supported export contribution under a fixed nutritional background. ER free sterol, cellular burden and secretion are distinct observables. Their observation maps must respect the definitions of C, H and V.

#### *S10.5. Exact equilibrium elimination and state boundaries*

With $a_N$=0, let $\mathbf{q} = (R, Y, H)$ and $\mathbf{w} = (C, N, M, P, S, K, I, V)$. The equations for $w$ can be solved sequentially at equilibrium as $\mathbf{w} = \psi(\mathbf{q})$, using positive sums of establishment and loss rates. The remaining equilibrium problem is:

$$\mathbf{f}_w(\boldsymbol{\psi}(\mathbf{q}), \mathbf{q}) = 0, \quad \mathbf{g}(\mathbf{q}) = \mathbf{f}_q(\boldsymbol{\psi}(\mathbf{q}), \mathbf{q}) = 0 \tag{S47}$$

For the reference case $a_N$=0, the elimination is explicitly evaluated in the following order. All quantities on the right are evaluated at the same candidate equilibrium; the supplied coordinates are $R$, $Y$ and H.

$$C = c_b + \kappa H, \quad A_N = \frac{\gamma_{\mathrm{n}}(b_N + a_{2N} o)}{1 + \chi r_C c_f}, \quad N = \frac{A_N}{A_N + d_N} \tag{S48}$$

$$A_M = b_M + a_{NM} h(N) + a_{2M} o, \quad M = \frac{A_M}{A_M + d_M + d_{RM} h(R)} \tag{S49}$$

$$A_P = b_P + a_{NP} h(N) + a_{2P} o, \quad P = \frac{A_P}{A_P + d_P + d_{RP} h(R)} \tag{S50}$$

$$A_S = b_S + a_{NS}^{+} h(N) + a_{2S} o, \quad S = \frac{A_S}{A_S + d_S + a_{NS} h(N)} \tag{S51}$$

$$A_K = b_K + a_{NK} h(N), \quad K = \frac{A_K}{A_K + d_K + a_{SK} h(S)} \tag{S52}$$

$$I = \frac{b_I}{b_I + d_I + \eta k_I e p_u}, \quad A_V = b_V + a_V[1 - h_I(I)], \quad V = \frac{A_V}{A_V + d_V} \tag{S53}$$

The auxiliary $A_N$, $A_M$, $A_P$, $A_S$, $A_K$ and $A_V$ denote algebraic establishment coefficients only. They are not additional state variables. Substitution into the $R$, $Y$ and $H$ residuals completes the three-coordinate equilibrium problem; all trajectories retain eleven coordinates.

With positive rate sums, every admissible reduced root reconstructs a full equilibrium and vice versa. This is equilibrium elimination, not a reduction of trajectory dimension. Nonzero Nrf1 autoregulation requires full-state root finding.

Differentiating the implicit relation for $w$ and applying the block determinant identity gives:

$$\mathbf{g}_q = \mathbf{J}_{qq} - \mathbf{J}_{qw}\mathbf{J}_{ww}^{-1}\mathbf{J}_{wq}, \quad \det\mathbf{J} = \det\mathbf{J}_{ww}\det\mathbf{g}_q \tag{S54}$$

For completeness, differentiating $f_w(\psi(q), q) = 0$ first gives $\mathbf{J}_{ww}\psi_q + \mathbf{J}_{wq} = 0$; hence $\psi_q = -\mathbf{J}_{ww}^{-1}\mathbf{J}_{wq}$. Applying the chain rule to $g(q) = f_q(\psi(q), q)$ then gives the first identity, and block Gaussian elimination gives the determinant identity. This connects the search method in main-text Section 2.3 to the full eleven-dimensional Jacobian.

The reduced residual locates zero-eigenvalue candidates because its eliminated block is invertible. Stability is evaluated with the full eleven-dimensional spectrum.

An index-one saddle has a one-dimensional unstable manifold and a ten-dimensional stable manifold. The two unstable branches identify destinations. A local basin-separating stable manifold requires the corresponding basin geometry. Fold checks additionally require a simple zero eigenvalue and the nonzero coefficients in Eq. (S18).

The input in main-text Section 2.4 is a rectangular pulse. For one selected source at a time, write:

$$u(t) = A_{\mathrm{p}}\,\mathbf{1}_{[0,T_{\mathrm{p}}]}(t), \qquad u(t) = 0 \quad t > T_{\mathrm{p}} \tag{S55}$$

Here $A_{\mathrm{p}}$ is the dimensionless pulse amplitude, $T_{\mathrm{p}}$ its duration, and the indicator is one during exposure and zero otherwise. For a sterol pulse replace $u$ by $u_H$. The other pulsed input remains zero. After washout the parameters and autonomous drift return to the reference values, but the initial condition for recovery is the state reached at pulse termination.

$$d_k = \|\mathbf{z}(T_{\mathrm{p}} + 250) - \mathbf{z}_k^*\|_2, \qquad k \in \{0,1\} \tag{S56}$$

Assign the endpoint to the smaller of the two distances and check that the endpoint is close to its assigned fixed point and has small drift. In the stored pulse grid the largest assigned distance was $4.58 \times 10^{-12}$, and halving the integration step did not alter assignments. The 16-step bisection retains an upper switching bracket; it is a numerical threshold bracket, not an exact analytical dose threshold.

Recovery depends on the full state at washout, not only on existence of a low-stress equilibrium. The pulse waveform, starting state, duration and coefficients define each switching threshold.

#### *S10.6. A-type, Itô and effective-event noise*

The A-type-to-Itô conversion follows Shi and colleagues [44], complementing the decomposition framework [25].

We use the descending-potential convention $\mathbf{f} = -(\mathbf{D} + \mathbf{Q})\nabla\varphi$, with $\mathbf{D}$ symmetric and $\mathbf{Q}$ antisymmetric [22,23,24,25,26]. The A-type density equation in S6 is equivalent to an Itô SDE whose drift is:

$$b_i = f_i + \varepsilon\partial_j(\mathbf{D}_{ij} + \mathbf{Q}_{ij}) \tag{S57}$$

To see the drift conversion explicitly, insert $f_i = -(\mathbf{D}_{ij} + \mathbf{Q}_{ij})\partial_j\varphi$ into the A-type density equation. The divergence of an antisymmetric tensor contracted with symmetric second derivatives vanishes. Expanding the remaining diffusion product yields the usual Itô forward operator:

$$\partial_t\rho = -\partial_i\{[f_i + \varepsilon\partial_j(\mathbf{D}_{ij} + \mathbf{Q}_{ij})]\rho\} + \varepsilon\partial_i\partial_j(\mathbf{D}_{ij}\rho) \tag{S58}$$

Antisymmetry cancels the contraction with symmetric second derivatives. The drift correction depends on spatial variation of both $\mathbf{D}$ and $\mathbf{Q}$. For compatible boundaries the stationary density can carry the Itô current $j_i = \varepsilon\partial_j(Q_{ij}\rho_{\mathrm{ss}})$, so stationarity does not require detailed balance.

Along deterministic trajectories of this representation:

$$\dot{\varphi} = -\nabla\varphi^{\mathrm{T}}\mathbf{D}\nabla\varphi \le 0 \tag{S59}$$

Positive-definite diffusion makes the mobility invertible, relating drift fixed points to potential critical points. A single smooth descending potential under these conditions precludes nontrivial periodic motion in its domain.

An effective-event comparison makes the flux noise convention explicit. Suppose each independent establishment or loss event changes $z_i$ by $1/\Omega$ and occurs at rate $\Omega a_i^+$ or $\Omega a_i^-$, respectively. Conditional Poisson moments give:

$$\mathrm{E}[\Delta z_i|z] = (a_i^+ - a_i^-)\Delta t, \qquad \mathrm{Var}[\Delta z_i|z] = \frac{a_i^+ + a_i^-}{\Omega}\Delta t \tag{S60}$$

When events are sufficiently numerous while propensities vary slowly over the chosen interval, a Gaussian approximation gives an Itô diffusion [45]. With $\varepsilon = 1/\Omega$ and the $\sqrt{2\varepsilon}$ amplitude convention:

$$\mathbf{D}_v = \frac{1}{2}\mathrm{diag}(a_i^+ + a_i^-), \qquad a_i^\pm = v_i^\pm/\tau_i \tag{S61}$$

Variance is formed from time-scaled event rates before taking the noise square root. Shared events would add off-diagonal diffusion. The event-derived Itô process and A-type process require the drift conversion above; their frozen constant-coefficient local forms agree.

Local Gaussian fluctuations require amplitudes small relative to distance from the boundary. A global stochastic simulation additionally needs a boundary-compatible process.

#### *S10.7. Local potential, covariance and the canonical decomposition*

For a stable fixed point, linearization and frozen diffusion give $\mathrm{d}\boldsymbol{\xi} = \mathbf{J}\boldsymbol{\xi}\mathrm{d}t + \sqrt{2\varepsilon}\mathbf{B}\mathrm{d}W$. Itô's product rule yields the covariance equation. The covariance equations below use centered fluctuations; their stationary linear mean is zero. If $\mathbf{J}$ is Hurwitz and $\mathbf{D}$ is positive definite:

$$\mathbf{J}\mathbf{C}_0 + \mathbf{C}_0\mathbf{J}^{\mathrm{T}} + 2\mathbf{D} = 0, \qquad \mathbf{C}_0 = 2\int_0^\infty \mathrm{e}^{Jt}\,\mathbf{D}\mathrm{e}^{\mathrm{J}^{\mathrm{T}}t}\mathrm{d}t, \qquad \boldsymbol{\Sigma} = \varepsilon\mathbf{C}_0 \tag{S62}$$

$$\frac{\mathrm{d}\boldsymbol{\Sigma}}{\mathrm{d}t} = \mathbf{J}\boldsymbol{\Sigma} + \boldsymbol{\Sigma}\mathbf{J}^{\mathrm{T}} + 2\varepsilon\mathbf{D}, \qquad \boldsymbol{\Sigma} = \mathrm{E}[\boldsymbol{\xi}\boldsymbol{\xi}^{\mathrm{T}}] \tag{S63}$$

More explicitly, $\mathrm{d}(\boldsymbol{\xi}\boldsymbol{\xi}^{\mathrm{T}}) = (\mathrm{d}\boldsymbol{\xi})\boldsymbol{\xi}^{\mathrm{T}} + \boldsymbol{\xi}(\mathrm{d}\boldsymbol{\xi})^{\mathrm{T}} + (\mathrm{d}\boldsymbol{\xi})(\mathrm{d}\boldsymbol{\xi})^{\mathrm{T}}$. Taking expectations removes the mean-zero Brownian terms and retains their quadratic variation. Stability ensures decay of the initial covariance and convergence of the integral in Eq. (S62).

The Brownian quadratic variation supplies $2\varepsilon\mathbf{D}$; stationarity and $\boldsymbol{\Sigma} = \varepsilon\mathbf{C}_0$ give the Lyapunov equation.

The integral converges and is positive definite. Set $\mathbf{U}=\mathbf{C}_0^{-1}$ and $\mathbf{Q} = -\mathbf{J}\mathbf{C}_0 - \mathbf{D}$. The Lyapunov equation gives $\mathbf{Q} + \mathbf{Q}^{\mathrm{T}} = 0$ and $\mathbf{J} = -(\mathbf{D} + \mathbf{Q})\mathbf{U}$, establishing the local construction [24]. The quadratic potential is $\varphi_\ell = \boldsymbol{\xi}^{\mathrm{T}}\mathbf{U}\boldsymbol{\xi}/2$; the corresponding quadratic form using $\boldsymbol{\Sigma}^{-1}$ is $\varphi_\ell/\varepsilon$, not $\varphi_\ell$. These two scales should not be interchanged.

Define $\mathbf{G} = (\mathbf{D} + \mathbf{Q})^{-1}$, $\mathcal{S} = (\mathbf{G} + \mathbf{G}^{\mathrm{T}})/2$ and $\mathcal{A} = (\mathbf{G} - \mathbf{G}^{\mathrm{T}})/2$. Multiplication shows $GDG^{\mathrm{T}} = \mathcal{S}$, connecting the mobility form to the dissipative/transverse form of the Ao decomposition. The calligraphic notation avoids confusing this matrix with p62 state S.

Local dispersion depends jointly on drift, diffusion and coupling. The separate local potentials have independent additive constants. Marginal precision is the inverse of the selected covariance submatrix.

The symmetric diffusion component determines dissipation of the local potential, while the antisymmetric component can drive motion along equal-cost contours [24,25]. Landscape descent therefore organizes recovery without requiring the full drift to be a gradient. This distinction also explains the direction-dependent uphill paths entering the action construction.

#### *S10.8. Linking action to potential differences*

For noise covariance 2 $\varepsilon$ $\mathbf{D}$, the leading weak-noise Gaussian path cost is:

$$\mathcal{I}_T[\mathbf{z}] = \frac{1}{4}\int_0^T[\dot{\mathbf{z}} - \mathbf{f}(\mathbf{z})]^{\mathrm{T}}\mathbf{D}^{-1}(\mathbf{z})[\dot{\mathbf{z}} - \mathbf{f}(\mathbf{z})]\mathrm{d}t \tag{S64}$$

The action applies to absolutely continuous paths with regular coefficients and positive-definite diffusion. Smooth bounded order-epsilon drift corrections do not change its leading weak-noise form.

For a compatible decomposition $\mathbf{f} = -(\mathbf{D}+\mathbf{Q})\nabla\varphi$, define $\mathbf{r} = \dot{\mathbf{z}} - (\mathbf{D}-\mathbf{Q})\nabla\varphi$. Expansion of the two quadratic forms, using $\mathbf{Q}^{\mathrm{T}} = -\mathbf{Q}$, gives the square-completion identity:

$$\mathcal{I}_T[\mathbf{z}] = \varphi(\mathbf{b}) - \varphi(\mathbf{a}) + \frac{1}{4}\int_0^{\mathrm{T}} \mathbf{r}^{\mathrm{T}}\mathbf{D}^{-1}\mathbf{r}\,\mathrm{d}t \tag{S65}$$

In detail, write $p = \nabla\varphi$ and $v = \dot{z}$. Antisymmetry implies $p^{\mathrm{T}}Qp = 0$ and gives the pointwise difference of the two quadratic costs:

$$[\mathbf{v} + (\mathbf{D}+\mathbf{Q})\mathbf{p}]^{\mathrm{T}}\mathbf{D}^{-1}[\mathbf{v} + (\mathbf{D}+\mathbf{Q})\mathbf{p}] - [\mathbf{v} - (\mathbf{D}-\mathbf{Q})\mathbf{p}]^{\mathrm{T}}\mathbf{D}^{-1}[\mathbf{v} - (\mathbf{D}-\mathbf{Q})\mathbf{p}] = 4\mathbf{v}^{\mathrm{T}}\mathbf{p} \tag{S66}$$

Dividing by four and integrating uses the chain rule, $\int_0^{\mathrm{T}} \dot{z}^{\mathrm{T}}\,\nabla\varphi\,\mathrm{d}t = \varphi(b) - \varphi(a)$, to obtain the identity above. The nonnegative residual vanishes only for the specified uphill equation.

The residual is nonnegative. Equality requires an admissible uphill connection $\dot{\mathbf{z}} = (\mathbf{D}-\mathbf{Q})\nabla\varphi$, generally with infinite-time departure from equilibrium. When $\mathbf{Q} \neq 0$, this is not simple reversal of the deterministic drift [26].

S7 computes constant-isotropic, finite-time, fixed-endpoint candidate costs. The discretization and numerical gradient are given in S11.3.

#### *S10.9. Coordinate consistency and verification*

If normalized variables are related to physical coordinates by a fixed affine map $\mathbf{z} = L\mathbf{x} + \mathbf{c}$, consistent drift and diffusion transformations are $f_z = Lf_x$ and $\mathbf{D}_z = LD_xL^{\mathrm{T}}$. The action quadratic form is then invariant. Assigning identity diffusion anew after an arbitrary rescaling changes the noise model. All comparisons therefore use matched coordinate and diffusion conventions.

The derivations establish invariance, equilibrium elimination, conditional bounds and matrix identities under the stated assumptions. The numerical checks below test their implementation.

Checks used 512 Sobol states per boundary face, repeated multistart root searches, 256 elimination tests and full-state Schur checks. At both attractors under both diffusion conventions, Lyapunov, antisymmetry, drift-reconstruction and inverse-mobility identities were checked. Maximum errors were $4.44 \times 10^{-16}$ for elimination, $8.88 \times 10^{-16}$ for the Schur identity and drift reconstruction, and $7.29 \times 10^{-15}$ relative error for action square completion. The archived record retains software versions, source checksum and tolerances.

### S11 Additional derivations and implementation crosswalk

#### *S11.1 From biological processes to the main equations*

Main-text Eqs. (3)-(5) follow by assigning nuclear Nrf1 support, Nrf2 induction and response turnover to establishment or loss. Parallel contributions add; jointly required functions multiply. S2 provides the explicit balances and S0 their coefficients.

Main-text Eq. (2) collects stress sources and relief. Eq. (6) describes membrane Nrf1-supported INSIG1 loss. Lower INSIG1 releases export, reducing sterol burden.

At fixed $P$ and intact interaction, INSIG1 decreases as membrane Nrf1 availability rises. At fixed remaining coefficients, the export program decreases as INSIG1 rises. The conditional signs are:

$$\frac{\partial I^*}{\partial e} = -\frac{b_I\eta k_I p_u}{(b_I+d_I+\eta k_I e p_u)^2} \leq 0, \quad \frac{\partial v_V^+}{\partial I} = -a_V h_I'(I)(1-V) \leq 0 \tag{S67}$$

These signs justify inhibition of INSIG1 abundance and inhibition of export by INSIG1. The graphical abstract instead names INSIG1 degradation, a process promoted by its upstream inputs.

#### *S11.2 Why a reference value is not uniquely determined by a steady state*

The stationary state fixes the ratio of establishment to loss. Their common scale remains free until dynamic information or a clock convention is added. In fact, the following transformation leaves the entire node drift unchanged for any positive constant assigned to that node:

$$\alpha_i \mapsto c_i\alpha_i, \quad \beta_i \mapsto c_i\beta_i, \quad \tau_i \mapsto c_i\tau_i, \quad c_i > 0 \tag{S68}$$

This coefficient-function transformation leaves drift and effective event rates unchanged. It is admissible only where the implemented family permits all associated rates to scale together. Fixed normalizations and shared coefficients restrict this freedom.

#### S11.3 Discrete path action and its gradient

Let a path with fixed endpoints contain $m$ segments, time increment $\delta = T/m$, midpoint $\bar{\mathbf{z}}_j = (\mathbf{z}_j + \mathbf{z}_{j+1})/2$, and velocity $\mathbf{v}_j = (\mathbf{z}_{j+1} - \mathbf{z}_j)/\delta$. Define the drift residual and a constant symmetric weight matrix:

$$\mathbf{r}_j = \mathbf{v}_j - \mathbf{f}(\bar{\mathbf{z}}_j), \quad \boldsymbol{\Lambda} = \mathbf{D}^{-1}, \quad \mathcal{J}_{T,m} = \frac{\delta}{4}\sum_{j=0}^{m-1} \mathbf{r}_j^{\mathrm{T}}\boldsymbol{\Lambda}\mathbf{r}_j \tag{S69}$$

The matrix $\boldsymbol{\Lambda}$ weights path deviations by inverse diffusion. The executed action calculation uses identity diffusion, so this weight is the identity. Differentiation of the segment residual with respect to its left and right nodes gives:

$$\partial_{\mathbf{z}_j}\mathbf{r}_j = -\mathbf{1}_{11}/\delta - \mathbf{J}_j/2, \quad \partial_{\mathbf{z}_{j+1}}\mathbf{r}_j = \mathbf{1}_{11}/\delta - \mathbf{J}_j/2 \tag{S70}$$

For the segment cost $\ell_j = \delta\mathbf{r}_j^{\mathrm{T}}\boldsymbol{\Lambda}\mathbf{r}_j/4$, the two column gradients are therefore:

$$\nabla_{\mathbf{z}_j}\ell_j = -\frac{1}{2}\boldsymbol{\Lambda}\mathbf{r}_j - \frac{\delta}{4}\mathbf{J}_j^{\mathrm{T}}\boldsymbol{\Lambda}\mathbf{r}_j \tag{S71}$$

$$\nabla_{\mathbf{z}_{j+1}}\ell_j = \frac{1}{2}\boldsymbol{\Lambda}\mathbf{r}_j - \frac{\delta}{4}\mathbf{J}_j^{\mathrm{T}}\boldsymbol{\Lambda}\mathbf{r}_j \tag{S72}$$

Each interior-node gradient is the sum of contributions from its adjacent segments. The fixed endpoints are excluded from the optimization vector. This is the gradient implemented in the action solver. A state-dependent diffusion would require additional derivatives of its inverse; those terms are absent because the executed path calculation has constant identity diffusion.

#### S11.4 Reading a local probability ellipse

Let $\boldsymbol{\Sigma}_{RP}$ be the two-coordinate covariance submatrix selected from the full local covariance. The displayed ellipse is a probability contour of the local Gaussian marginal, not a deterministic basin boundary:

$$\mathbf{x}^{\mathrm{T}}\boldsymbol{\Sigma}_{RP}^{-1}\mathbf{x} \le r^2, \quad \Pr(\chi_2^2 \le r^2) = 0.683, \quad r^2 = -2\ln(1 - 0.683) = 2.2957489 \tag{S73}$$

A covariance submatrix defines the marginal distribution; a submatrix of the full precision defines a different conditional distribution.

#### S11.5 Mathematical notation and implementation fields

The companion parameter ledger contains the complete 74-entry mathematical-to-code crosswalk. The full parameter definitions and values remain in S0; code identifiers are kept out of the mathematical equations.

### S12 Mechanism-constrained parameterization and ensemble analysis

#### S12.1 Operating levels turnover and regulatory response

Ao and colleagues constructed metabolic kinetics by specifying operating concentrations and fluxes, then matching unknown coefficients through the rate laws [46]. Their one-substrate example illustrates the principle:

$$v(a) = V_{\max}\frac{a}{K_a + a}, \qquad V_{\max} = v_0\left(1 + \frac{K_a}{a_0}\right) \tag{S74}$$

Choosing $K_a = a_0$ gives $V_{\max} = 2v_0$. For this functional network, the transferable idea is to separate working level, turnover and regulatory shape. At an interior operating point, establishment and loss balance:

$$J_i^* = \alpha_i^*(1 - z_i^*) = \beta_i^* z_i^*, \quad r_i^* = \frac{J_i^*}{\tau_i} = \lambda_i z_i^*(1 - z_i^*), \quad \frac{\alpha_i^*}{\tau_i} = \lambda_i z_i^*, \quad \frac{\beta_i^*}{\tau_i} = \lambda_i(1 - z_i^*) \tag{S75}$$

The normalized level alone fixes a ratio. A matched turnover or frozen-input rate supplies the additional scale. The S0 values define the reference realization; computed working states explain those values and check their implementation. At the low-stress point, $P^* = 0.932871$, establishment is 1.143209 and loss is 0.082265. With $\tau_P = 1.45$, the frozen-input rate is 0.845155.

For proteasome establishment and Nrf2 turnover, solving the stationary balances gives:

$$a_{NP} = \frac{\tau_P\lambda_P P^* - b_P - a_{2P}o^*}{h(N^*)}, \qquad d_P + d_{RP}h(R^*) = \tau_P\lambda_P(1 - P^*) \tag{S76}$$

$$k_{KP} = \frac{\gamma_2[b_Y + a_{RY}h(R^*)](1 - Y^*)/Y^* - d_Y}{K^*P^*} \tag{S77}$$

The first denominator requires $h(N^*) > 0$; the second requires $Y^* > 0$ and $K^*P^* > 0$. At the reference low-stress point, these reconstruct $a_{NP} = 1.15$ and $k_{KP} = 1.35$. At high stress, the Nrf2-supported proteasome input increases to 0.105169, but stress-dependent loss rises to 0.420863; proteasome function falls to 0.713664 despite the compensatory input.

Reconstructed gains must also be nonnegative to belong to the parameter domain in S10.2. Nonzero denominators ensure that the algebraic inversions are defined; nonnegativity ensures that the resulting coefficients retain their stated establishment or removal roles.

The membrane and export coefficients follow from the corresponding balances:

$$k_I = \frac{b_I(1-I^*)/I^* - d_I}{\eta \mathrm{e}^* p_u^*} \tag{S78}$$

$$k_V = \frac{j_H(1-H^*)/H^* - d_H - k_{2H} o^*}{V^*} \tag{S79}$$

At low stress, $\mathrm{e}^* = 0.446258$, $p_u^* = P^* = 0.932871$ and $I^* = 0.179759$ recover $k_I = 2$. The export level $V^* = 0.784634$ and burden $H^* = 0.170900$ recover $k_V = 0.80$. The same four reconstructions at three reference equilibria agree with the stored values to within $1.14 \times 10^{-14}$. All 33 node-level flux balances agree to within $2.00 \times 10^{-15}$.

Figure S7 summarizes the operating-point parameterization and its reconstruction checks; Figure S8 shows how the resulting drift enters the dynamical analyses.

#### *S12.2 Constructing compatible parameter sets*

RACIPE motivates systematic variation of rates and thresholds within a regulatory topology [47,48]. Structural Kinetic Modeling separates turnover scales from local response slopes [49]. Ensemble Modeling retains several kinetic realizations compatible with operating states and perturbations [50]. These approaches inform parameter construction without requiring a unique coefficient set.

Observation-conditioned parameterization can combine qualitative directions and quantitative readouts through an explicit observation relationship [51].

First specify the coordinate definitions, reference clock and observation maps. Functional assays may constrain state levels or rates; expression profiles can constrain directions or intervals through a stated observation relationship. Next propose gate shapes and structural factors within the nonnegative domain of S10.2. For an active input $x_0$ at gate occupancy $g_0$, the threshold is:

$$K = x_0\left(\frac{1-g_0}{g_0}\right)^{1/n} \tag{S80}$$

Here $x_0 > 0$, $n > 0$ and $0 < g_0 < 1$, with $g_0 = h(x_0)$. Occupancies at zero or one are limiting cases rather than finite positive-threshold assignments. These conditions make the inversion in Eq. (S80) well defined and preserve the positive-threshold response family.

At fixed gates and candidate operating states, amplitudes that enter the balances affinely can be collected across conditions into one constrained system:

$$\mathbf{B}\boldsymbol{\vartheta} = \mathbf{d}, \qquad \boldsymbol{\ell} \le \boldsymbol{\vartheta} \le \mathbf{u} \tag{S81}$$

Each shared parameter occupies one column. For proteasome establishment the columns are $1$, $h(N^*)$ and $o^*$; independent variation and total-establishment information are needed to distinguish the three gains. Small singular values or a broad profile likelihood identify weakly constrained combinations [52,53,54]. Uncertain operating states require repeating the construction over their allowed ranges or solving the resulting nonlinear problem.

The exact regulatory elasticities connect rate laws to local feedback strength:

$$\frac{\partial \ln h(x)}{\partial \ln x} = n[1-h(x)], \qquad \frac{\partial \ln[1-h(x)]}{\partial \ln x} = -nh(x) \tag{S82}$$

$$J_{ij} = \lambda_i z_i^*(1-z_i^*)\frac{E_{ij}^{\alpha} - E_{ij}^{\beta}}{z_j^*} - \lambda_i \delta_{ij} \tag{S83}$$

Here $E_{ij}^{\alpha}$ and $E_{ij}^{\beta}$ are logarithmic rate elasticities at an interior equilibrium. The full Jacobian spectrum determines stability; constructing a balanced operating point does not predetermine that spectrum. With positive total rates, the continuous map below maps the unit cube to itself, so Brouwer's theorem supplies at least one equilibrium:

$$T_i(\mathbf{z}) = \frac{\alpha_i(\mathbf{z})}{\alpha_i(\mathbf{z}) + \beta_i(\mathbf{z})}, \qquad \mathbf{z}^* = \mathbf{T}(\mathbf{z}^*) \quad \Leftrightarrow \quad \mathbf{f}(\mathbf{z}^*) = 0 \tag{S84}$$

A data-conditioned ensemble is the next step: retain candidates satisfying independently specified observations and mathematical constraints, freeze each accepted vector, and then classify steady states, compensation targets and washout outcomes. Single-state cases, successful bypasses and unresolved numerical cases remain in the outcome record. Bistability and failed rescue are outputs, not acceptance requirements.

#### *S12.3 Response shape and existing ensemble comparisons*

Four archived 256-candidate designs detected at least two stable equilibria in 165 nine-dimensional linear proposals, 149 mixed-log proposals, 113 expanded fixed-exponent proposals and 39 corrected fifteen-dimensional proposals. Each count belongs to its own ranges and sampling rule. The corrected design uses exponents 1-4; its per-exponent counts are 0, 0, 10 and 29 among 64 candidates each. The earlier rounded design that included exponent 5 is excluded from this comparison.

The matched response-shape experiment fixes all other S0 coefficients and evaluates four shared exponents (1, 2, 3, 4) at eight thresholds (0.30, 0.35, 0.40, 0.42, 0.45, 0.50, 0.55, 0.60). Dedicated sterol and INSIG1 gates are unchanged. At exponents 1 and 2, one stable point was detected at every threshold. At exponent 3, two stable points and an index-one saddle were found at thresholds 0.50 and 0.55. At exponent 4, this structure occurred at 0.35, 0.40, 0.42, 0.45 and 0.50. Thus switching depends jointly on response steepness and operating range.

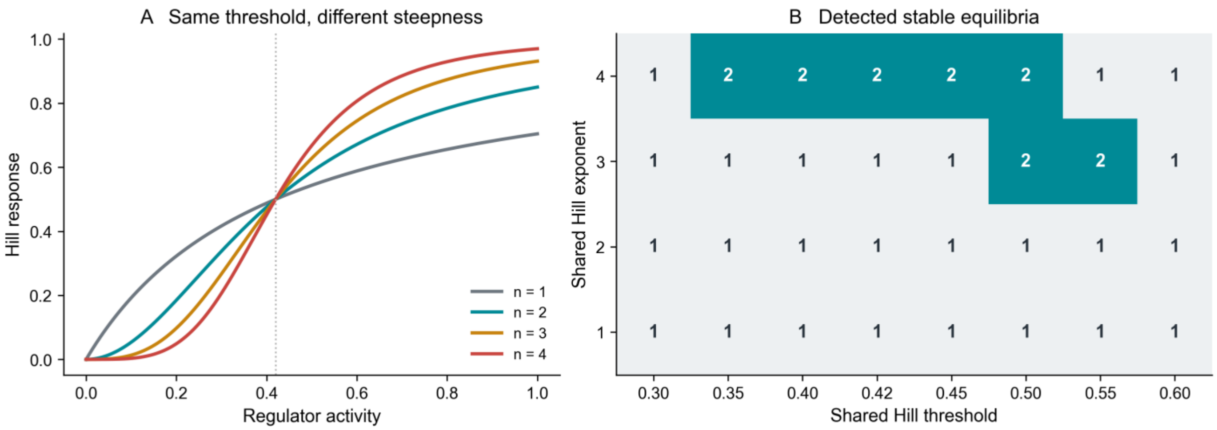


Figure S4. Gate shape and equilibrium multiplicity. Left, response curves at the reference threshold 0.42. Right, detected stable-state counts in all 32 matched cases. Other coefficients remain at S0; dedicated sterol and INSIG1 gates are unchanged.

#### *S12.4 Expanded maintenance and Nrf2 compensation*

We compared nuclear-input loss with simultaneous nuclear and membrane-input loss. Both basal maintenance coefficients were multiplied by 1, 2, 4, 8, 16 or 32; both Nrf2 maintenance gains by 0, 1, 2, 4, 8, 16 or 32; and Nrf2 production by 1 or 4. All 168 interventions began from the corresponding unrescued persistent-stress equilibrium and ran for 500 time units. Target recovery required proteasome and maintenance levels at least 0.932871 and 0.911334, respectively, with stress no greater than 0.106507 (tolerance $10^{-6}$).

With membrane Nrf1 retained, fourfold Nrf2 production and 32-fold basal maintenance allowed all seven support-gain settings to reach the three targets. None did so at reference Nrf2 production. With both Nrf1 inputs absent, two of 42 fourfold-production cases met the maintenance targets, but none met the stress target. The successful nuclear-loss case with zero Nrf2 maintenance gain reached P = 0.951672, M = 0.940572 and R = 0.091787. By contrast, dual-input loss with 32-fold basal and regulator-dependent support reached P = 0.957979 and M = 0.935050 while R remained 0.232645. The comparison separates maintenance restoration from control of the continuing sterol-associated source.

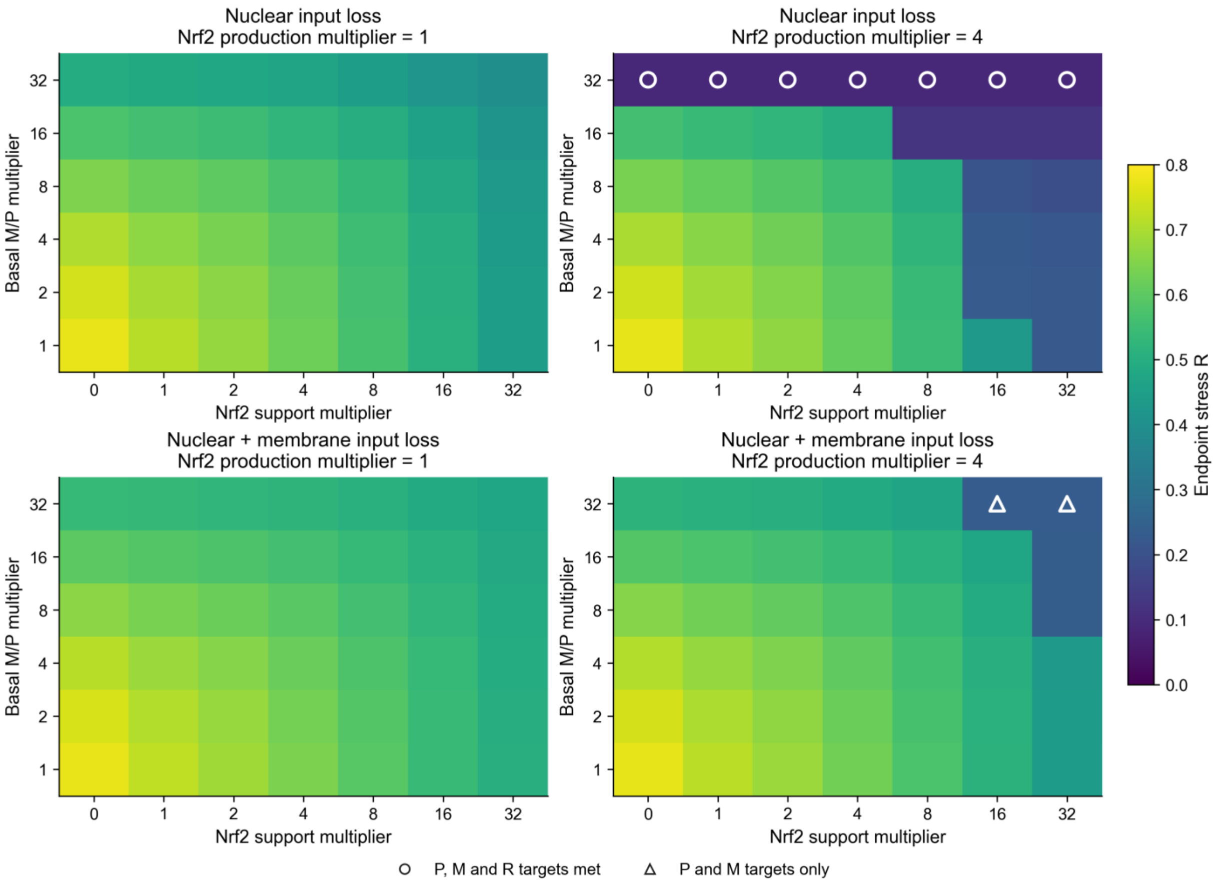


Figure S5. Compensation under expanded support. Squares show stress after 500 time units. Rows vary basal maintenance; columns vary Nrf2 maintenance gains. Open circles meet all three functional targets; triangles meet only proteasome and maintenance targets. All 168 virtual interventions are retained, including successful bypasses.

#### *S12.5 Relative clocks and stress exposure*

Six clock settings were compared: reference; Nrf2 time multiplied by 0.25 or 4; both maintenance and proteasome times multiplied by 0.25 or 4; and all times multiplied by 2. Every setting retained the three reference equilibria and their full-Jacobian stability classes. From the low-stress state, six amplitudes (0, 0.04, 0.08, 0.16, 0.32, 0.64) and three durations (0.5, 2, 8) produced 108 stress-pulse protocols, each followed for 500 unforced units.

All endpoints were within $10^{-5}$ of a reference attractor. High-stress outcomes occurred in 8/18 reference protocols, 8/18 for each Nrf2 clock change, 9/18 for faster maintenance, 8/18 for slower maintenance and 7/18 for uniform slowing. Twelve-step bisection refined the duration-two threshold, with independent Radau checks at both bracket endpoints:

| Clock setting | Lower switching bound | Upper switching bound |
|---|---|---|
| Reference | 0.107500 | 0.107656 |
| Nrf2 time x 0.25 | 0.107500 | 0.107656 |
| Nrf2 time x 4 | 0.107188 | 0.107344 |
| M/P times x 0.25 | 0.100938 | 0.101094 |
| M/P times x 4 | 0.114375 | 0.114531 |
| All times x 2 | 0.181250 | 0.181406 |

Changing a node time accelerates both establishment and loss. Faster maintenance dynamics can therefore also accelerate stress-induced deterioration during exposure. For a common rescaling, let $c > 0$ multiply every node time while retaining the same initial condition. The external waveform is evaluated at the rescaled time, so a rectangular pulse and its comparison time both have their durations multiplied by $c$. The resulting drift and trajectory obey:

$$\mathbf{f}_c = \frac{\mathbf{f}}{c}, \qquad u_c(t) = u(t/c), \qquad \mathbf{z}_c(t) = \mathbf{z}(t/c) \tag{S85}$$

For a twofold common scale change, doubling pulse and washout durations reproduced the reference trajectory with maximum coordinate error $1.48 \times 10^{-8}$. Holding the pulse duration fixed instead changes exposure relative to the network clock.

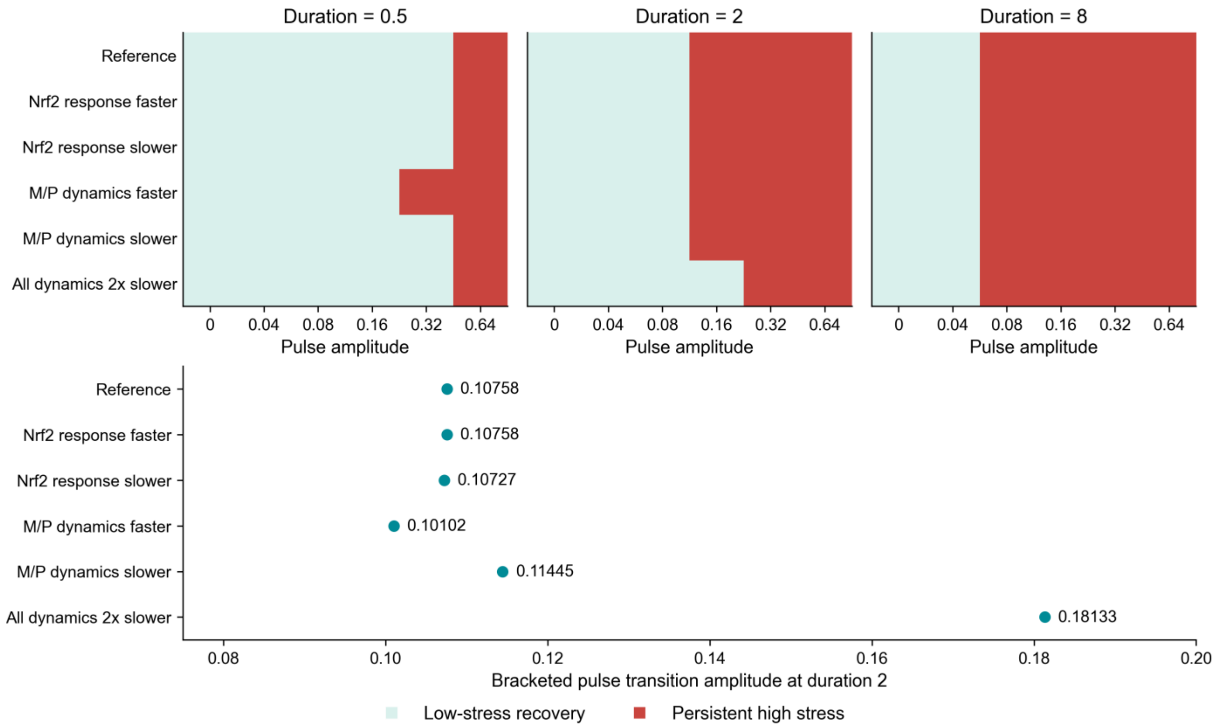


Figure S6. Clock dependence of recovery. Upper maps retain all 108 pulse protocols. Lower markers show refined duration-two transition brackets; error bars span the brackets. The six clock settings retain equilibrium positions and stability while altering finite-exposure responses.

#### *S12.6 Numerical checks and reconstruction files*

The matched experiments used 128 scrambled Sobol starts (seed 20261025) plus seven structured starts. Roots required full-state residual below $10^{-8}$, domain tolerance $10^{-7}$, merge distance $10^{-5}$ and stability margin $10^{-7}$. All 27,405 attempts across 203 searches are retained, including 7,276 solver failure flags; acceptance was assessed using residual and domain checks.

Trajectories used DOP853 with relative tolerance $10^{-8}$, absolute tolerance $10^{-10}$ and maximum step 0.5. Seventeen systematically spaced cases were repeated with tighter Radau tolerances; maximum endpoint difference was $2.30 \times 10^{-8}$. The refined pulse search used tolerances $10^{-9}$ and $10^{-11}$, maximum step 0.25, and twelve additional Radau endpoint checks. The elasticity Jacobian agreed with complex-step differentiation to $3.89 \times 10^{-15}$. Both attractors under both diffusion conventions gave positive covariance matrices, Lyapunov residuals below $2.36 \times 10^{-14}$ and drift-factorization residuals below $2.99 \times 10^{-15}$.

The companion tables provide all gate, compensation and pulse cases, equilibrium attempts, transition brackets, independent solver checks, operating values and coefficient reconstructions. The matched-design files specify the intervention settings before execution. The existing exploratory ensembles and these completed experiments remain distinct from the proposed observation-conditioned parameter construction in S12.2.

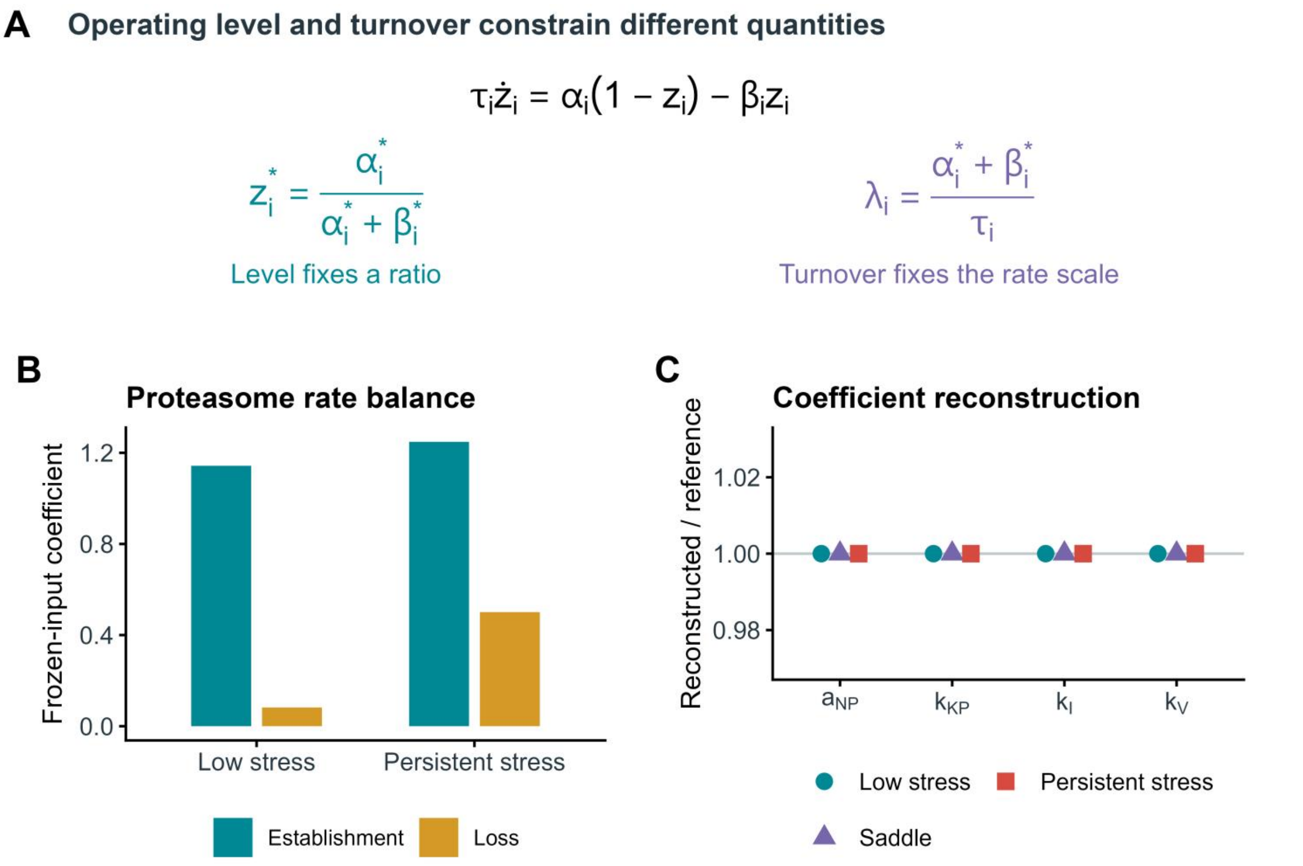


Figure S7. Operating levels, turnover and coefficient reconstruction. A, At an interior operating point, the normalized level fixes the establishment-to-loss ratio, whereas the frozen-input rate also involves their common scale and the relative node time. The coefficients are evaluated with regulatory inputs held fixed; the collective network relaxation is determined by the full Jacobian. B, Archived establishment and loss coefficients for proteasome function at the low-stress and persistent-stress equilibria. C, Four coefficients reconstructed independently at each of the three reference equilibria, divided by their assigned values. The Nrf1-supported proteasome, joint KEAP1-proteasome, membrane-dependent INSIG1 turnover and sterol export coefficients have reference values 1.15, 1.35, 2 and 0.80, respectively. Symbols correspond to S0 and Eqs. (S76)-(S79). The reconstruction uses model-computed equilibria and checks algebraic consistency; it does not estimate coefficients from independent experimental measurements. Source values are provided in the accompanying operating-point and coefficient-reconstruction tables.

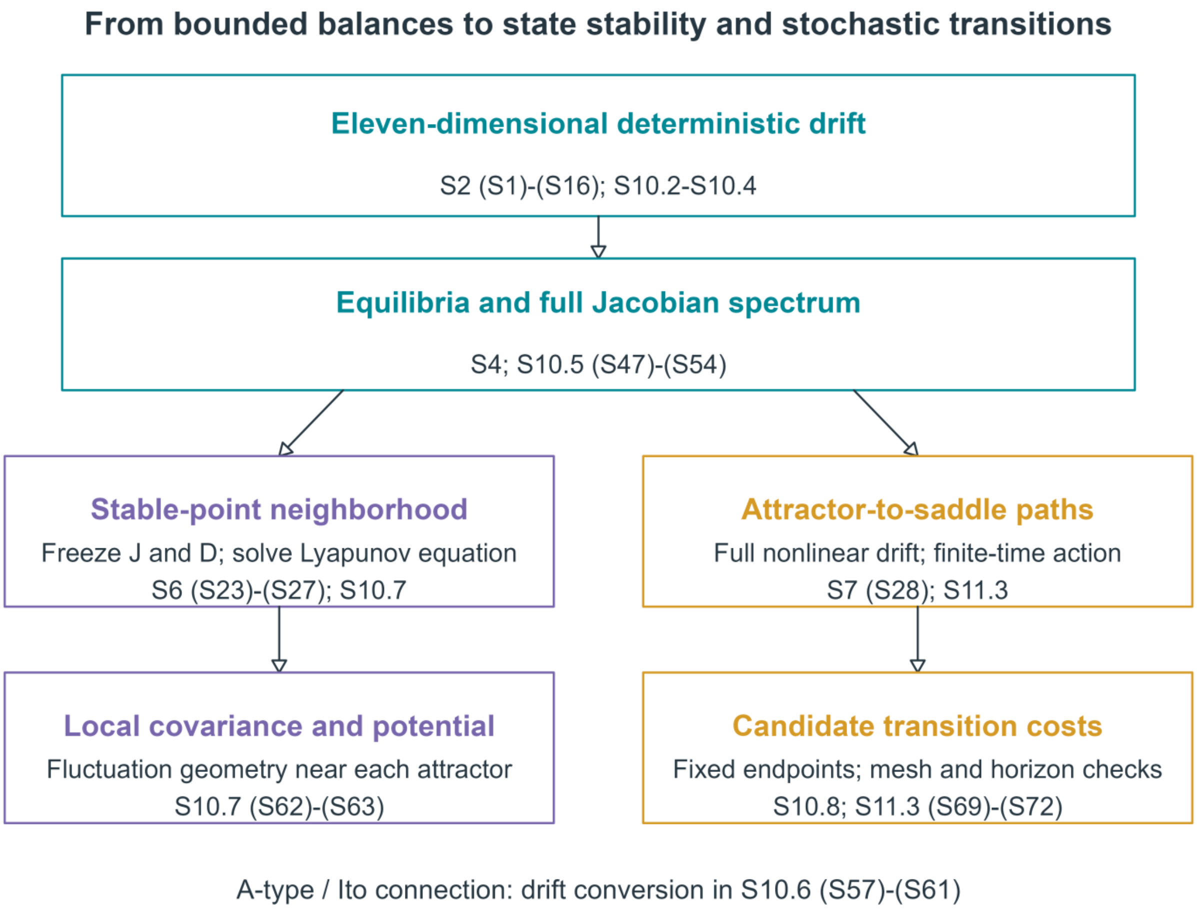


Figure S8. Mathematical organization of the deterministic and stochastic analyses. The complete eleven-dimensional balances define the deterministic drift. Equilibrium elimination assists root finding, while stability is evaluated with the full Jacobian. At each stable point, frozen drift and diffusion coefficients lead through the Lyapunov equation to local covariance and a quadratic local potential. Separately, the nonlinear drift enters the fixed-endpoint, finite-time action calculation from each attractor to the saddle. The diagram identifies the sections and numbered equations for each construction. The A-type-to-Itô conversion is given in S10.6; the local constant-coefficient construction and finite-time nonlinear path calculation are distinct operations.

Together, these analyses separate three properties of redox recovery: existence and deterministic accessibility of low-stress states, local fluctuation curvature around each attractor, and action accumulated along a candidate transition path. Parameter and clock perturbations in S12 test the first property; the two diffusion conventions test the second; path and horizon checks test the third. These quantities provide complementary descriptions of Nrf1-Nrf2 network resilience.

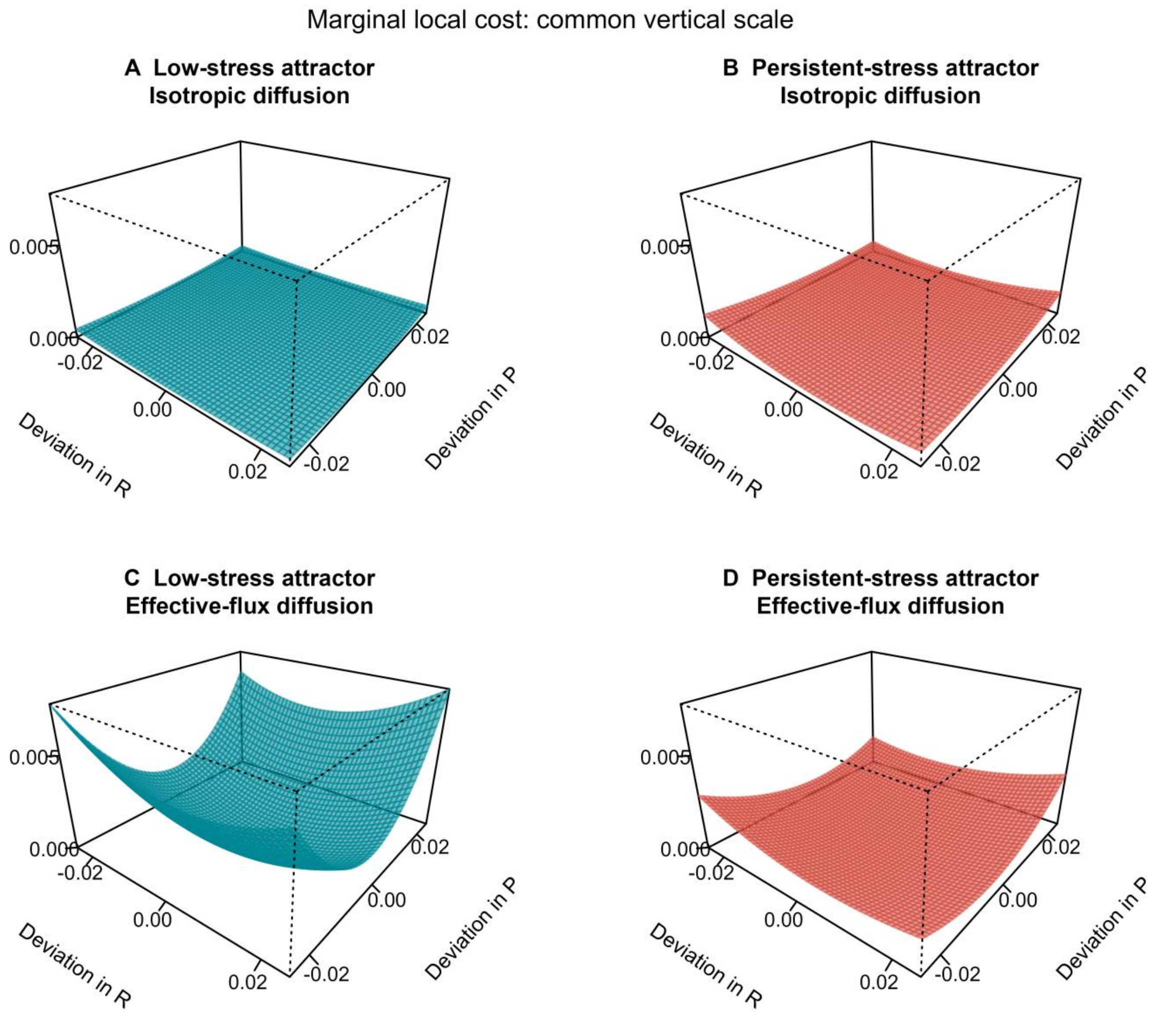


Figure S9. Local landscape geometry near the two reference attractors. A-B, Isotropic diffusion; C-D, effective-flux diffusion. Each surface is the stress/proteasome marginal quadratic cost, calculated as one half of the displacement vector contracted with the inverse of the archived covariance-per-unit-noise submatrix. Its associated Gaussian marginal density is proportional to the exponential of minus this cost divided by noise intensity. The panels use deviations from the respective attractor, not absolute state coordinates, with a common range of -0.025 to 0.025 and a shared vertical scale. Each minimum is assigned zero independently. The displayed geometry follows S6 and S10.7; marginalization follows S11.4. Source covariance coefficients and their Lyapunov checks are supplied with the figure.

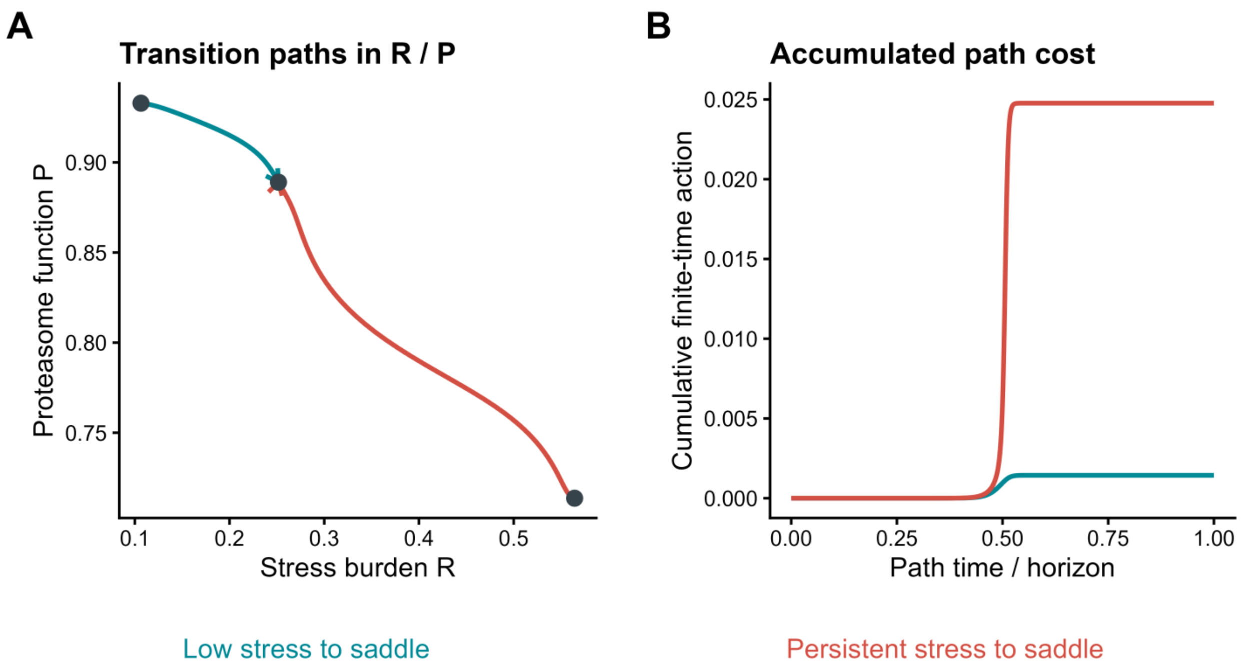


Figure S10. Transition paths and cumulative action link state switching to the landscape formulation. A, Archived eleven-dimensional finite-time optimized paths from each reference attractor to the saddle, projected onto stress burden and proteasome function. Arrows indicate progression toward the saddle; the projection does not define a two-dimensional basin boundary. B, Stored cumulative action along the same paths, with path time divided by the horizon. Both paths use isotropic diffusion, horizon 96 and 960 segments. Their terminal costs are 0.001436689 and 0.024767131 for low-stress and persistent-stress departure, respectively. S7 specifies the numerical protocol and S10.8 relates path cost to a compatible potential difference.
